# Compressive forces induce epigenetic activation of aged human dermal fibroblasts through ERK signaling pathway

**DOI:** 10.1101/2024.11.04.621794

**Authors:** Hui Liu, Luezhen Yuan, Lucrezia Baldi, Trinadha Rao Sornapudi, G.V. Shivashankar

## Abstract

Age-related changes in human dermal fibroblasts (HDFs) contribute to impaired wound healing and skin aging. While these changes result in altered mechanotransduction, the epigenetic basis of rejuvenating aging cells remains a significant challenge. This study investigates the effects of compressive forces on nuclear mechanotransduction and epigenetic rejuvenation in aged HDFs. Using a compressive force application model, the activation of HDFs through alpha-smooth muscle actin (ɑ-SMA) is demonstrated. Sustained compressive forces induce significant epigenetic modifications, including chromatin remodeling and altered histone methylation patterns. These epigenetic changes correlate with enhanced cellular migration and rejuvenation. Small-scale drug screening identifies the extracellular signal-regulated kinase (ERK) signaling pathway as a key mediator of compression-induced epigenetic activation. Furthermore, implanting aged cell spheroids to an aged skin model and subjecting the tissue with compressing forces resulted in increased collagen I protein levels. Collectively, these findings demonstrate that applying compressive force to aged fibroblasts activates global epigenetic changes through the ERK signaling pathway, ultimately rejuvenating cellular functions with potential applications for wound healing and skin tissue regeneration.

**Significance Statement:** Partial rejuvenation of aging cells is desirable but is still a major challenge. In this paper, we demonstrate that aged human dermal fibroblasts, embedded in a 3D collagen hydrogel matrix as spheroids, subjected to external static compressive force exhibit partial rejuvenation. Through immunofluorescence, small-scale inhibitor screen and gene expression analysis, we identify some of the critical mechanotransduction pathways in this process. Collectively, our results provide compelling evidence that tissue compression results in the activation of potential rejuvenation pathways in aging cells.

## Introduction

Cellular aging is accompanied by various changes in the characteristics of cells, such as genomic instability, telomere attrition, epigenetic changes, loss of proteostasis, deregulated nutrient sensing, mitochondrial dysfunction, cellular senescence, stem cell exhaustion, and altered intercellular communication (1). For example, dermal fibroblasts secrete various extracellular matrix proteins (ECM) into the dermal compartment and contribute to matrix stiffness of the skin (2). Many studies have shown that aging leads to accumulation of senescent fibroblasts resulting in decreased ECM production thereby leading to a loss of skin tissue integrity and of wound healing properties (3). A major challenge in the field is how such aging cells can be activated or rejuvenated. Current strategies to combat aging include induced pluripotent stem cells (iPSC) and mechanical reprogramming (4, 5), metabolic manipulation (daily or intermittent caloric restriction), blood transfusion, small molecule drugs (Rapamycin, Metformin, Ascorbate and Aspirin) and senescent cell ablation (Senolytics) (6). Mechanical forces (stretch, shear, compression) have also been shown to activate major signaling pathways, cytoskeleton/chromatin remodeling and gene expression (7–10). Since cells sense extracellular mechanical cues in tissue microenvironments, we hypothesized that compressive forces could enable the activation/rejuvenation of aging cells. Recent literature has also shown that cancer cells under tissue compression could get activated to a metastatic phenotype (11). Based on previous studies, including our own, on the effects of compressive force on cellular function (12, 13), we designed an engineered tissue embedded with aging cells and revealed that tissue compression could provide important avenues for cell activation/rejuvenation.

In this paper, we develop a force application device, which includes human dermal fibroblasts (derived from a 75 year old healthy male donor) embedded in a 3D collagen hydrogel matrix and subjected to external loading in the form of static compressive force. Using a fibroblast spheroid model, we show that HDFs can be activated by compressive force, as evidenced by increased levels of ɑ Smooth Muscle actin (ɑSMA) and the accompanying cellular memory responses. In particular, we measure the levels of phosphorylated myosin light chain (pMLC) levels, cytoskeletal remodeling, chromatin modifications, reduced DNA damage, global gene expression, and cell migration to demonstrate the activation of HDFs. A key element of the activation process involves the clumping of aged cells into spheroids before the application of force, as single cells embedded in a collagen matrix under compressive force do not undergo activation. We also validated our findings in an artificial aged skin model and observed an increase trend in collagen 1protein levels in the spheroid injection group compared to the single-cell injection group. Collectively, our results provide compelling evidence that tissue compression results in the activation of potential rejuvenation of aging cells.

## Results

### Compressive force induces transient activation of fibroblasts and promotes rejuvenation

We established two models for 3D cell culture: one involves embedding aged fibroblasts as single cells in a collagen hydrogel (hereafter called the single cell model), and the other involves aged fibroblasts as spheroids embedded in a collagen hydrogel (called spheroid model) (Figure 1A, Figure S1E). To achieve the spheroid model, GM08401 fibroblasts (75 year old donor, old group) and GM09503 fibroblasts (10 year old donor, young group) were cultured on fibronectin-coated micropatterns overnight to form spheroids with diameters ranging from 50 μm to 150 μm (Figure S1B), resulting in spheroids with cell numbers ranging from 30 to 100 (Figure S1C). We added collagen hydrogel 1 mg/ml concentration on top of the spheroids and applied a metal or glass ring to confine the 3D matrix, preventing collagen hydrogel shrinkage during the cell culture process (see Methods). Finally, a compressive force ∼5% strain (referred as 1xload) and ∼15% strain (referred as 2xload) was added on top of the collagen hydrogel (Figure S1A, S1D and S1E) followed by culturing for 48 hours.

**Figure 1.**
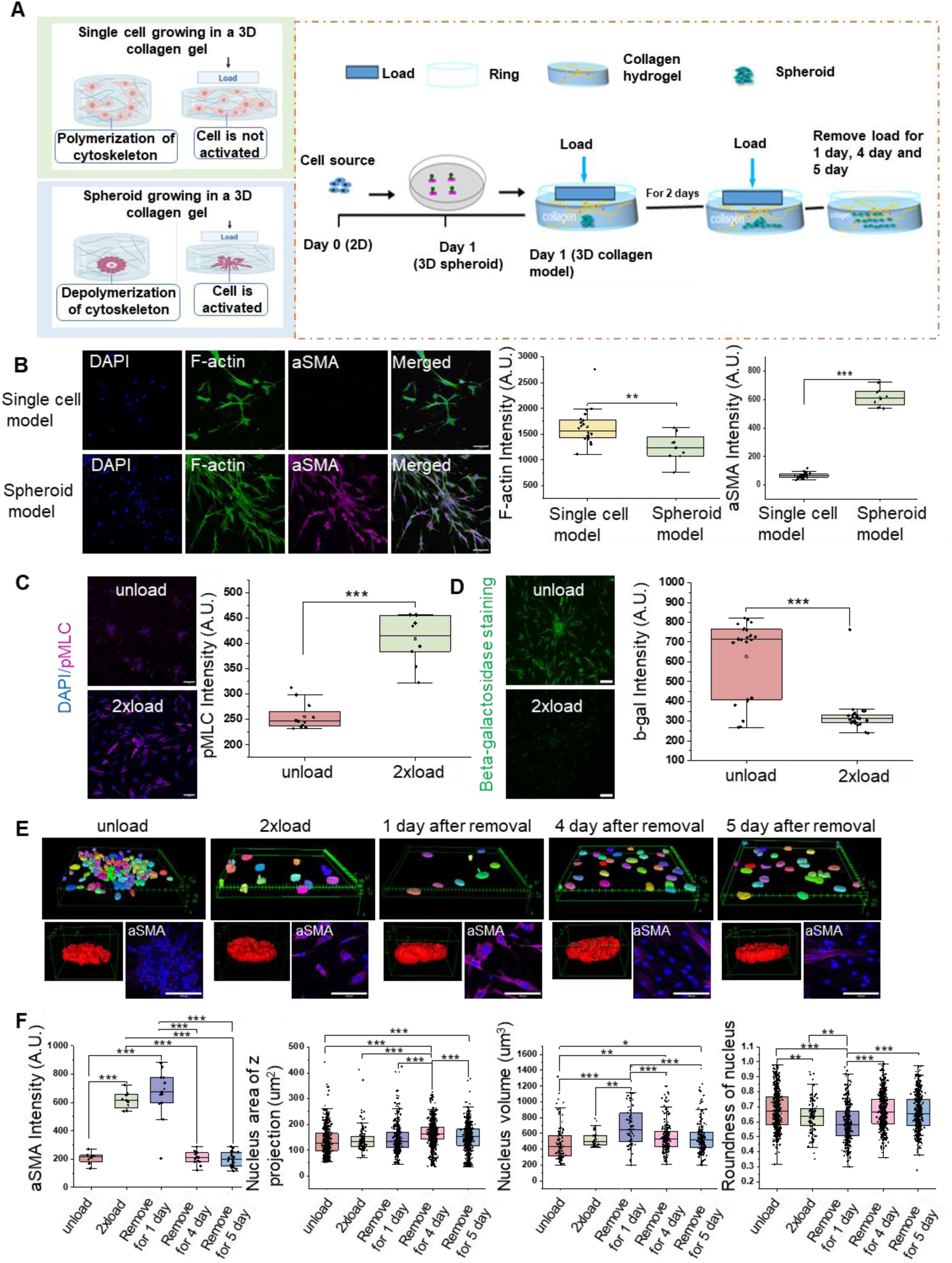
Establishment of 3D collagen hydrogel in-vitro model upon compressive forces and characterization activation, rejuvenation and memory properties of the regenerated phenotypes. (A) Schematic of 3D in-vitro collagen model (single cell embedding and spheroid embedding) and illustration of the cell culture process. (B) Representative ɑSMA immunofluorescence confocal images and quantification data per image of mean intensity in single cell model and spheroid. Nucleus is labeled in blue. (Scale bar, 100 μm). (C) Representative pMLC immunofluorescence confocal images and quantification data per image of mean intensity. Nucleus is labeled in blue. (Scale bar, 100 μm). (D) Representative Beta-galactosidase staining confocal images and quantification data per image of mean intensity. (Scale bar, 100 μm). (E) 3D nucleus construction and representative ɑSMA immunofluorescence confocal images under load and load removal condition. (Scale bar, 100 μm). (Unit in green box is μm). (F) Quantification data of ɑSMA mean intensity per image, nucleus volume, Z project area of nucleus and roundness of nucleus. All the experiments were repeated at least three times independently with similar results. P values in Figure (B-D) were calculated by unpaired, two-tailed Student’s t test. P values in Figure (F) were calculated by the one-way ANOVA method with Tukey’s post hoc test. *P<0.05; **<0.01; ***P<0.001; No asterisks means not significant. Source data are provided as a Source Data file.

After two days of culture, we evaluated fibroblast activation using immunofluorescence markers such as ɑSMA and pMLC (Figure 1B and C). We found that cells in the single cell 3D model showed low expression of ɑSMA, when compared to the 3D spheroid model (Figure 1B), although the F-actin expression level is higher in the single cell model as shown in Figure 1B. These results suggest that the activation of fibroblasts was more pronounced in spheroid models, highlighting the importance of compressive forces in 3D spheroid to regulate cellular function. In subsequent studies, we will therefore use the spheroid model to assess its applications to cellular rejuvenation.

Under compressive force conditions, both old and young fibroblasts became activated, as indicated by the increased level of ɑSMA compared to the unload group (Figure S3A and Figure S3B). The increased level of pMLC also implies that these cells are in an active state (Figure 1C). Since senescence is a hallmark of aging, we sought to determine if the application of compressive force alters senescence-associated properties. Towards this we performed beta-galactosidase staining and found fewer positively stained cells in the 2xload group, suggesting that the applied load reduces senescence of aged cells as shown in Figure 1D. Next, we measured the persistence of the activated state of the fibroblasts in the 2xload group. To do this we removed the load after culturing the cells in 3D under load condition for 48 hours and continued to culture for 5 days as shown in Figure 1E and 1F. We found that the ɑSMA level increased at day 1 and then decreased at day 4 and 5 (Figure 1F) suggesting that fibroblasts were transiently activated. Further, many of the cellular and nuclear morphometric parameters that changed with compressive force were also reversed upon the removal of compressive force (Figure 1F and Figure S3C and S3D). In summary, the increased cell contractility, as evidenced by elevated ɑSMA and pMLC levels, along with reduced senescence under compressive force conditions and the ability to revert back to a non-activated state after removing the load, suggesting that our spheroid model with load application has the potential to induce rejuvenation properties in aged cells.

### Mechanical force stabilizes microtubules and facilitates chromatin remodeling

Since cellular aging is accompanied by alterations in cytoskeletal and chromatin remodeling, we next evaluate the role of compressive forces on such cytoskeletal and chromatin remodeling. Upon application of compressive forces, microtubule reorganization was more evident as shown in Figure 2A, compared to F-actin (Figure S2E). α-tubulin intensity, a component of microtubules, is higher after the application of compressive load, and the microtubule network is much more complex compared to the unload group (Figure S9A-C). In addition, Lamin A/C, a nuclear protein playing a key role in force transmission from the cytosol into the nucleus (14), showed not much significant differences between the two groups (Figure 2A). Lamin B interacts with chromatin and contributes to nuclear organization, by anchoring heterochromatin to the nuclear periphery (15). Compared to unload group, Lamin B intensity increased in 2xload group (Figure 2A and Figure S6C).

**Figure 2.**
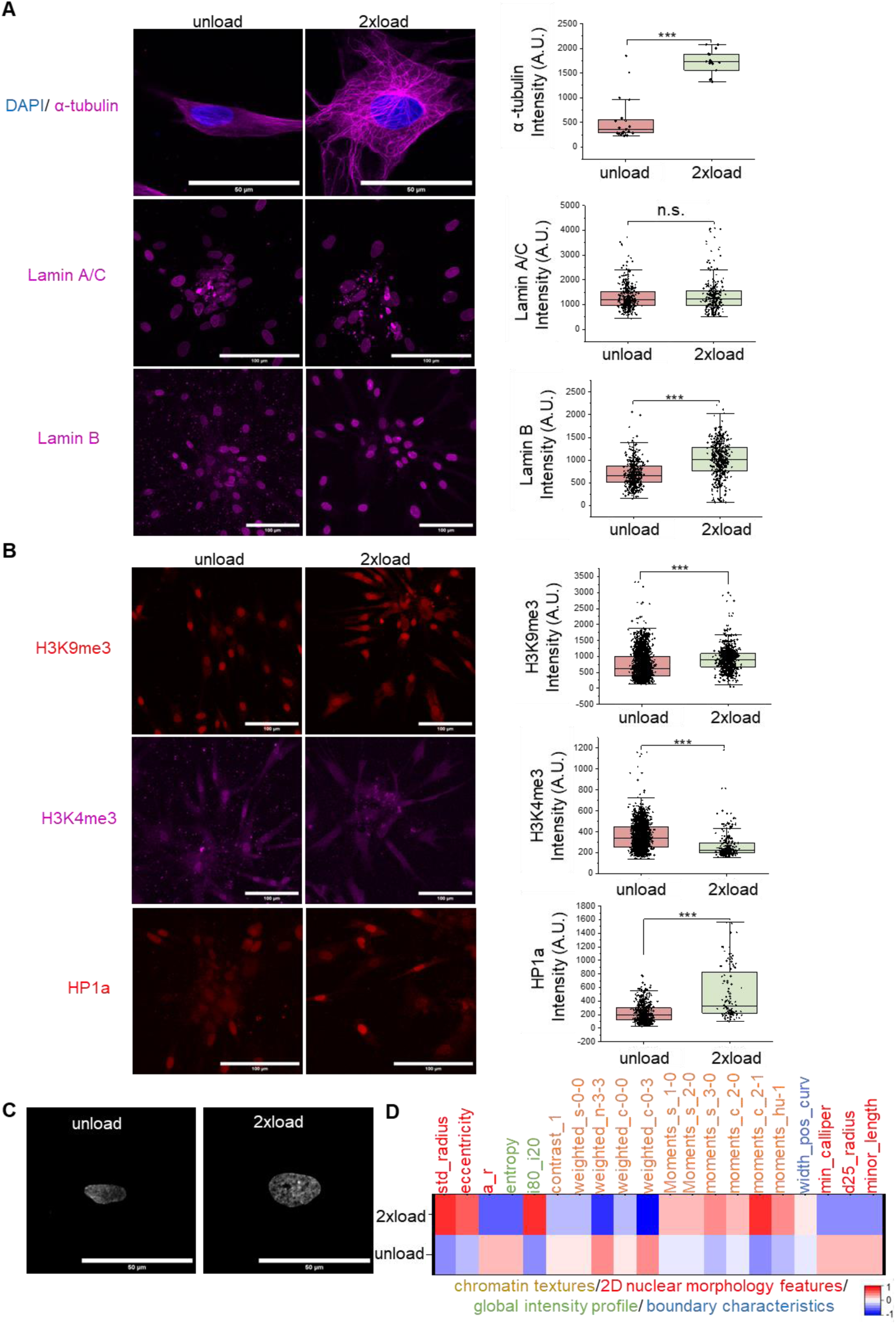
Mechanical force stabilizes microtubule and facilitates chromatin remodeling. (A) Representative α-tubulin, Lamin A/C and Lamin B immunofluorescence confocal images and quantification data per cell of mean intensity. Nucleus is labeled in blue. (Scale bar, 50 or 100μm). (B) Representative H3K9me3, H3K4me3 and HP1a immunofluorescence confocal images and quantification data per nucleus of mean intensity. (Scale bar, 100μm). (C) Representative gray images from DAPI in unload condition and load condition. (Scale bar, 50 m). (D) Heatmap of chromatin and nucleus morphology analysis. All the experiments were repeated at least three times independently with similar results. P values in Figure (A and B) were calculated by unpaired, two-tailed Student’s t test. *P<0.05; **<0.01; ***P<0.001; No asterisks means not significant. Source data are provided as a Source Data file.

Cells undergo chromatin remodeling in response to mechanical forces as a protective mechanism to maintain genome integrity (16). Our subsequent investigation focused on chromatin organization, highlighting key players such as H3K9me3 (Histone 3 Lysine 9 Trimethylation), H3K4me3 (Histone 3 Lysine 4 Trimethylation) and HP1a (Heterochromatin Protein 1 alpha). H3K9me3 and H3K4me3 are commonly associated with distinct chromatin regions, with H3K9me3 linked to heterochromatin and H3K4me3 associated with euchromatin (17, 18), while HP1a, binding to H3K9me3, plays a crucial role in forming and maintaining heterochromatin structures (19). In line with these findings, compressive forces on HDFs resulted in increased levels of H3K9me3, and HP1a, and decreased levels of H3K4me3, as indicated in Figure 2B and Figure S6A, S6B. Interestingly, upon removal of the load, HP1a level exhibited an increase trend as shown in Figure S6B. Consistent with Figure 2B, high resolution DAPI-stained images of nuclei showed increased puncta like structure in 2xload group and an analysis of nuclear morphology and chromatin intensity features demonstrated condensed chromatin under mechanical load (Figure 2C) and an increased ratio of heterochromatin to euchromatin (Figure 2D, Figure S6D and TableS4). Taken together, these findings suggest that mechanical force not only stabilizes microtubules but also contributes to increased chromatin condensation possibly contributing to genome stability.

### Mechanical force enhances cell migration primarily through nucleus-cytoskeleton axis

Upon observing the activation of fibroblasts and increased microtubule organization with compressive forces, we subsequently investigated its role on cell migration. Since cellular aging results in reduced cell migration, the goal of these experiments was to assess if mechanical forces increased cell migration, as possible routes to cellular rejuvenation. In both old and young groups, under compressive force conditions, cell migration was significantly higher than in the unloaded groups, as shown in Figure 3A and Figure S2A. In addition, cell migration speed in the aged cells increases with the applied load, as depicted in Figure S2B. Figure S2C and S2D also shows cell migration enhanced with compressive load and after removal of the load. Since cellular perception of compressive forces are transduced via membrane proteins such as G protein-coupled receptors (GPCRs), Piezo, integrins, and calcium channels, and transduce these signals into the nucleus via cytoskeletal components including actin, microtubules, and intermediate filaments (20, 21), we carried out a small-scale drug screen to identify critical signaling intermediates in our compressive force induced HDF activation and migration.

**Figure 3.**
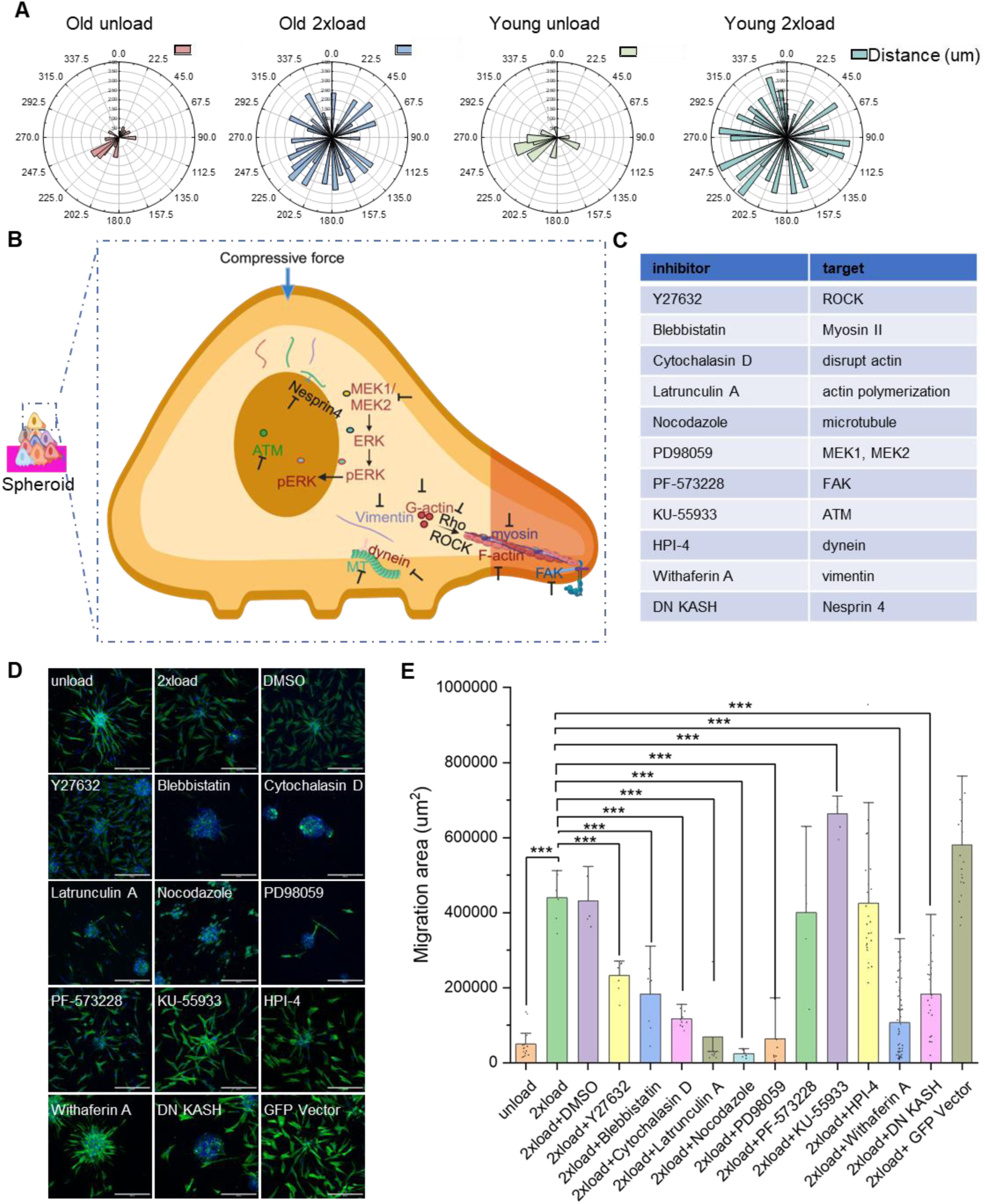
Mechanical load enhances cell migration via nucleus-cytoskeleton axis. (A) Windrose plots displaying the distance of the migrated cell nucleus to the center of one spheroid. (B) Schematic illustration of inhibitor targets. (C) Table for inhibitors’ description. (D) Representative immunofluorescence confocal images to check spheroid spreading. Nucleus is labeled in blue. F-actin is labeled in green. (Scale bar, 300 mm). (E) Quantification data of the spread area of the spheroid. All the experiments were repeated at least three times independently with similar results. P values in Figure (E) were calculated by unpaired, two-tailed Student’s t test. Other groups are compared to the 2xload group. *P<0.05; **<0.01; ***P<0.001; No asterisks means not significant. Source data is provided as a Source Data file.

In Figure 3B-E (large area as shown in Figure S4A-C), we applied several inhibitors to perturb possible intermediators shown in Figure 3C. We found that Latrunculin A, Nocodazole, and PD98059 are the three most effective inhibitors which inhibit cell migration among these interventions. Apart from the above-mentioned inhibitors, the Y27632 (ROCK inhibitor) group reduces the cell migration area but the cell number increases. On the other hand, in the PF-573228 (FAK inhibitor (22)) group, cell migration was not significantly affected compared to the 2xload group. Collectively, we identified that inhibition of actin, microtubules and ERK pathway had a critical role in force induced HDF activation and migration. Given the specific roles of ERK pathway in aging and rejuvenation, we next assessed the interplay between compressive forces, chromatin organization, DNA damage and transcription control with a particular focus on ERK inhibition.

### Mechanical load orchestrates chromatin reorganization, and diminishes DNA damage through ERK Signaling

Based on our findings showing that PD98059 inhibits cell migration to a greater extent, our next step is to investigate the effect of ERK inhibitor on chromatin remodeling and gene expression. As depicted in Figure 4A, immunofluorescence of pERK reveals that ERK undergoes phosphorylation and translocates into the nucleus upon compressive forces. After adding drug PD98059, pERK level decreased in the nucleus as shown in Figure 4B (images are shown in Figure S8).

**Figure 4.**
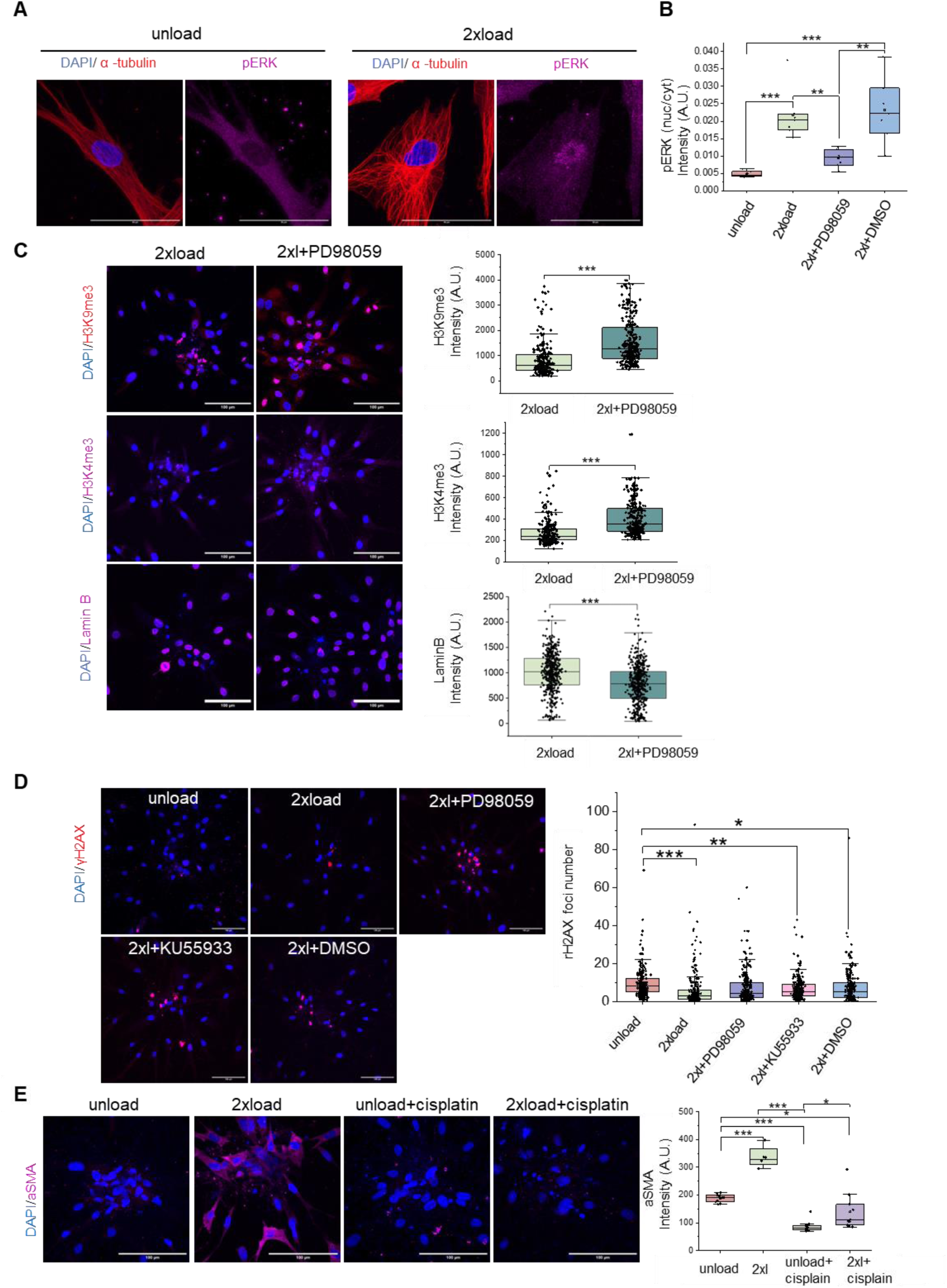
ERK role in chromatin remodeling, DNA damage response and gene expression regulation. (A) Representative ERK phosphorylation immunofluorescence confocal images and quantification data of mean intensity per cell (some images shown in Figure S8). Nucleus is labeled in blue. α-tubulin is labeled in red. pERK is labeled in magenta (Scale bar, 50μm). (B) Representative H3K9me3, H3K4me3 and Lamin B immunofluorescence confocal images and quantification data of mean intensity per nucleus. Nucleus is labeled in blue. (Scale bar, 100μm). (D) Representative γH2AX immunofluorescence confocal images and quantification data of foci number per nucleus. (Scale bar, 100μm). (E) Representative ɑSMA immunofluorescence confocal images and quantification data of mean intensity per image. Nucleus is labeled in blue. (Scale bar, 100μm). All the experiments were repeated at least three times independently with similar results. P values in Figure (B, D, E) were calculated by the one-way ANOVA method with Tukey’s post hoc test. P values in Figure (C) were calculated by unpaired, two-tailed Student’s t test. *P<0.05; **<0.01; ***P<0.001; No asterisks means not significant. Source data is provided as a Source Data file. “2xL” is an abbreviated notation for “2x load.”

Next, we measured H3K9me3 and H3K4me3 levels and observed an increase of these markers in the PD98059 group as shown in Figure 4C and Figure S6A. Increase in H3K9me3 and H3K4me3 suggests that ERK plays a role in the regulation of chromatin organization through histone modification. Lamin B expression level decreased after adding PD98059 in Figure 4C and Figure S6C. This suggests that the nuclear translocation of pERK may potentially affect chromatin structure and the activity of transcriptional regulators.

We then examined whether cells experienced DNA damage by assessing γH2AX, a well-established DNA damage marker (23). Surprisingly, our observations in Figure 4D revealed a notable mitigation of cellular DNA damage under mechanical load as indicated in foci number. In Figure S7A, where the fixed unload group and fixed load group serve as control groups, DRAQ7 staining data revealed the appearance of dead cells in the spheroid center under compressive force conditions. In Figure S7B, upon addition of the drug PD98059, we observed an increase in the level of γH2AX in the unload condition, indicating an elevation in DNA damage. To further investigate the impact of mechanical load on DNA damage response, we introduced cisplatin, a known inducer of DNA damage. Remarkably, in the 2xload+cisplatin group, cellular damage decreased compared to the unload+cisplatin group (Figure S7C).

The observed protective effects against DNA damage prompts the hypothesis that compressive force promotes DNA repair. Next, we sought to investigate whether ATM, a key protein kinase, is involved in the cellular response to DNA damage under force conditions. Surprisingly, in our study (Figure 4D), the administration of KU55933 (an ATM inhibitor) did not significantly affect DNA damage compared to 2xload group, and cell migration remained unaffected (Figure 3D and 3E). Similarly, there were no statistically significant changes observed in the γH2AX levels between the 2xl+PD98059 group and the load groups. Moreover, the γH2AX level in the 2xl+PD98059 group shows a non-significant decreasing trend compared to the control group (Figure 4D). This prompts the hypothesis that compressive force prevents DNA damage not via an ATM-dependent ERK signaling pathway, but possibly through a mechanism involving physical force-induced chromatin interactions. Next, we investigated whether DNA damage affects the activation properties under mechanical force using immunofluorescence levels of ɑSMA. As shown In Figure 4E, the immunofluorescence images from all four conditions showed lower levels of ɑSMA in cisplatin groups compared to compressive load conditions. These findings underscore the intricate involvement of ERK in governing cellular processes, ranging from chromatin organization to DNA damage response.

### Coupling between ERK signaling and differential gene expression upon compressive forces

To further characterize transcription profiles and gene expression changes associated with enhanced migration, rejuvenation, and ERK signaling pathways, we conducted global RNA sequencing analysis under different conditions as shown in Figure S10A-D. From the RNA-Seq analysis, we observed significant upregulation of 278 genes in the 2xl group compared to the unload condition (fold change > 2, adjusted p value < 0.1) (Figure 5A), and 612 genes were upregulated in the 2xl group compared to the PD98059 group (Figure 5B). Gene Ontology (GO) biological process analysis of these differentially expressed genes (278 and 612 DEGs) revealed enrichment in cell migration along with other important cellular processes (Figure 5A and 5B).

**Figure 5.**
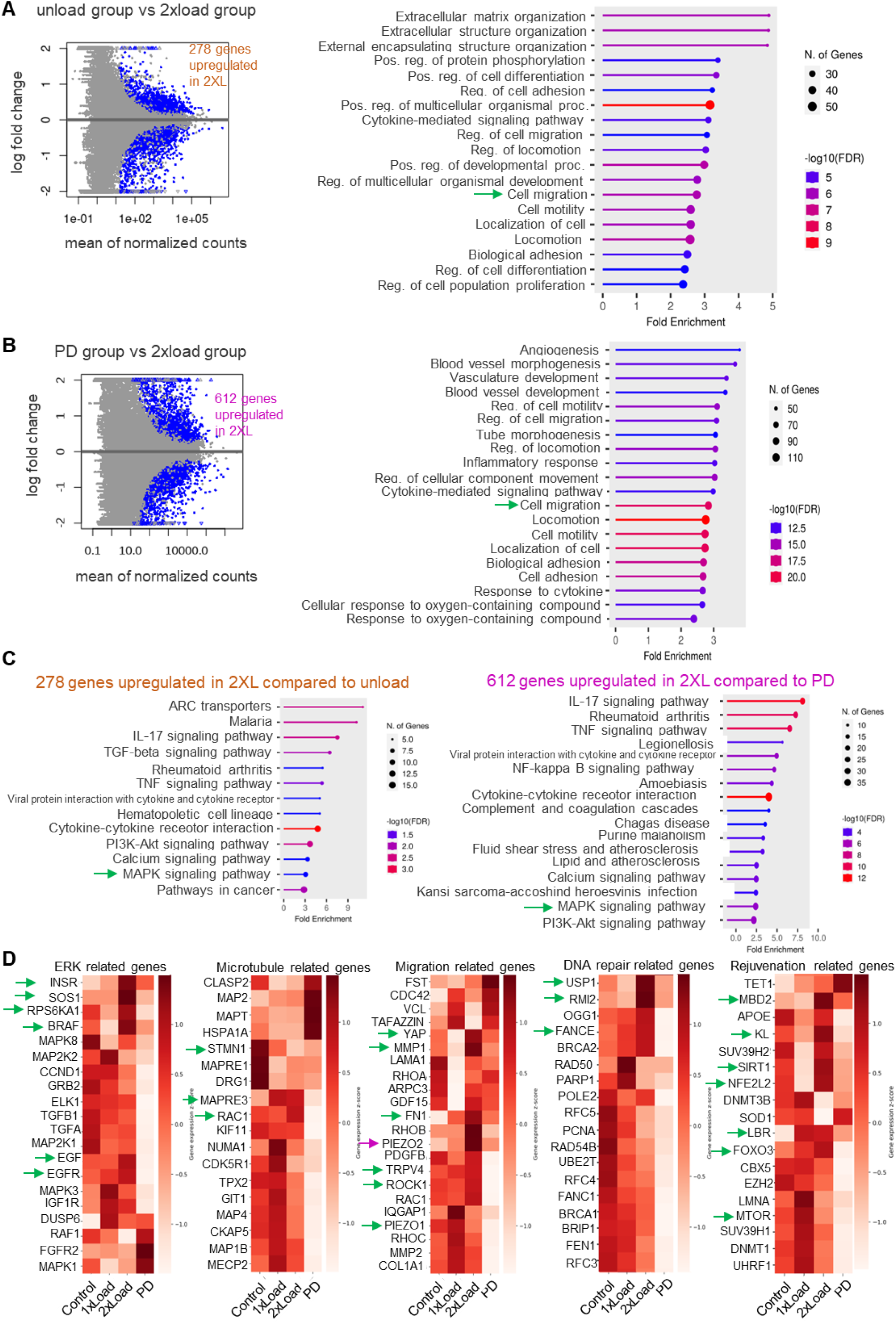
RNAseq analysis. (A) Volcano plot of significant genes in 2xload group compared to unload condition and GO Biological process analysis. (fold change > 2, adjusted p value < 0.1). 278 genes upregulated in 2xload compared to unload. (B) Volcano plot of significant genes in 2xload group compared to PD condition and GO Biological process analysis. PD condition means PD98059 inhibitors plus load condition. (fold change >1, adjusted p value < 0.1). 612 genes upregulated in 2xload compared to PD. (C) KEGG pathway analysis in above two DEG lists (278 DEG list and 612 DEG list). (D) The heatmaps show gene expression level in different groups such as ERK-related genes, microtubule-related genes, migration-related genes, DNA repair-related genes and rejuvenation-related genes.

Kyoto Encyclopedia of Genes and Genomes (KEGG) pathway analysis of the 278 DEGs highlighted the involvement of Mitogen-activated protein kinases (MAPKs) during this process (Figure 5C and Figure S12A). GO cellular component analysis of the 278 DEGs revealed that they are associated with extracellular matrix, while GO molecular function analysis indicated enrichment in growth factor activity, extracellular matrix structural constituent, and metallopeptidase activity (Figure S10E). Similarly, KEGG analysis of the 612 DEGs validated the importance of MAPKs in this process (Figure 5C and Figure S12B). GO cellular component analysis for the 612 DEGs showed enrichment in the extracellular matrix, while GO molecular function analysis highlighted their association with growth factor receptor activity (Figure S10F). The overlap of 119 genes between the 278 DEGs and 612 DEGs showed consistent results in terms of GO biological process analysis, GO cellular component analysis, GO molecular function analysis, and KEGG pathway analysis (Figure S10D and S10G). Additionally, FOXO signaling pathway involvement was noted in KEGG pathway analysis, underscoring its significance in rejuvenation.

Subsequent analysis of gene expression using DE-seq normalized results revealed changes in several genes. We divided these into groups based on their enrichment in the functional pathways such as ERK-related genes, microtubule-related genes, migration-related genes, DNA repair-related genes, and rejuvenation-related genes (Figure 5D). Among the ERK-related genes, downregulation was observed in most genes in the PD98059 group, while upregulation of Insulin Receptor (*INSR*), Son of Sevenless homolog 1 (*SOS1*), Ribosomal Protein S6 Kinase A1 (*RPS6KA1*), B-Raf Proto-Oncogene (*BRAF*), Epidermal Growth Factor (*EGF*), and Epidermal Growth Factor Receptor (*EGFR*) in the 2xload group suggested their involvement in ERK activation, directly or indirectly. Microtubule-related genes exhibited diverse expression patterns, notably characterized by the downregulation of *STMN1*, which is associated with microtubule destabilization, and the upregulation of *MAPRE3* and *RAC1* in the 2xload group, influencing microtubule dynamics and organization. Migration-related gene analysis, including ECM-related genes, mechanosensor-related genes, and Rho signaling pathway-related genes, using qRT-PCR, revealed consistent trends with part of RNA-seq data analysis (Figure S11). DNA repair-related mechanisms encompassing base excision repair, nucleotide excision repair, homologous recombination, mismatch repair, Fanconi anemia pathway, and non-homologous end-joining were examined (KEGG pathway information reviewed in Figure S13 A-F). Notably, upregulation of *USP1*, *RMI2*, and *FANCE* in the 2xload group, associated with the Fanconi anemia pathway, was observed among DNA repair-related genes. Rejuvenation-related gene analysis revealed the upregulation of *MBD2*, *KL*, *SIRT1*, *NFE2L2*, *LBR*, and *FOXO3*, alongside the downregulation of *MTOR* in 2xload group, all of which are associated with longevity (KEGG pathway information reviewed in Figure S14 A-D).

In summary, our RNA-seq results align with the observations of enhanced migration, involvement of the ERK signaling pathway, and cellular rejuvenation.

### Compressive forces on implanted spheroids in an FT AGED skin model show aged fibroblast activation

To explore the potential applications in translational medicine, we utilized an artificial aged skin tissue model to investigate whether aged fibroblasts could be activated (Figure S15). We injected either single cells or spheroids into the skin tissue, followed by the application of compressive force or no force as a control. Cells were localized around the injection site, as shown in Figures S16A and S16B. Compared to the control group (without cell injection), we observed relatively higher levels of αSMA and collagen I protein in the experimental groups with cell injections (Figures S17 and S18). Under compressive force conditions, collagen I expression was higher in the spheroid group compared to the single-cell group (Figure 6A). Similarly, elastin protein levels in the spheroid group under compressive force showed an increasing trend compared to the single-cell group under the same conditions (Figure S18). However, under the current experimental conditions, the levels of fibronectin and elastin were much less upregulated with compressive force compared to collagen 1 secretion (Figures S17 and S18).

**Figure 6.**
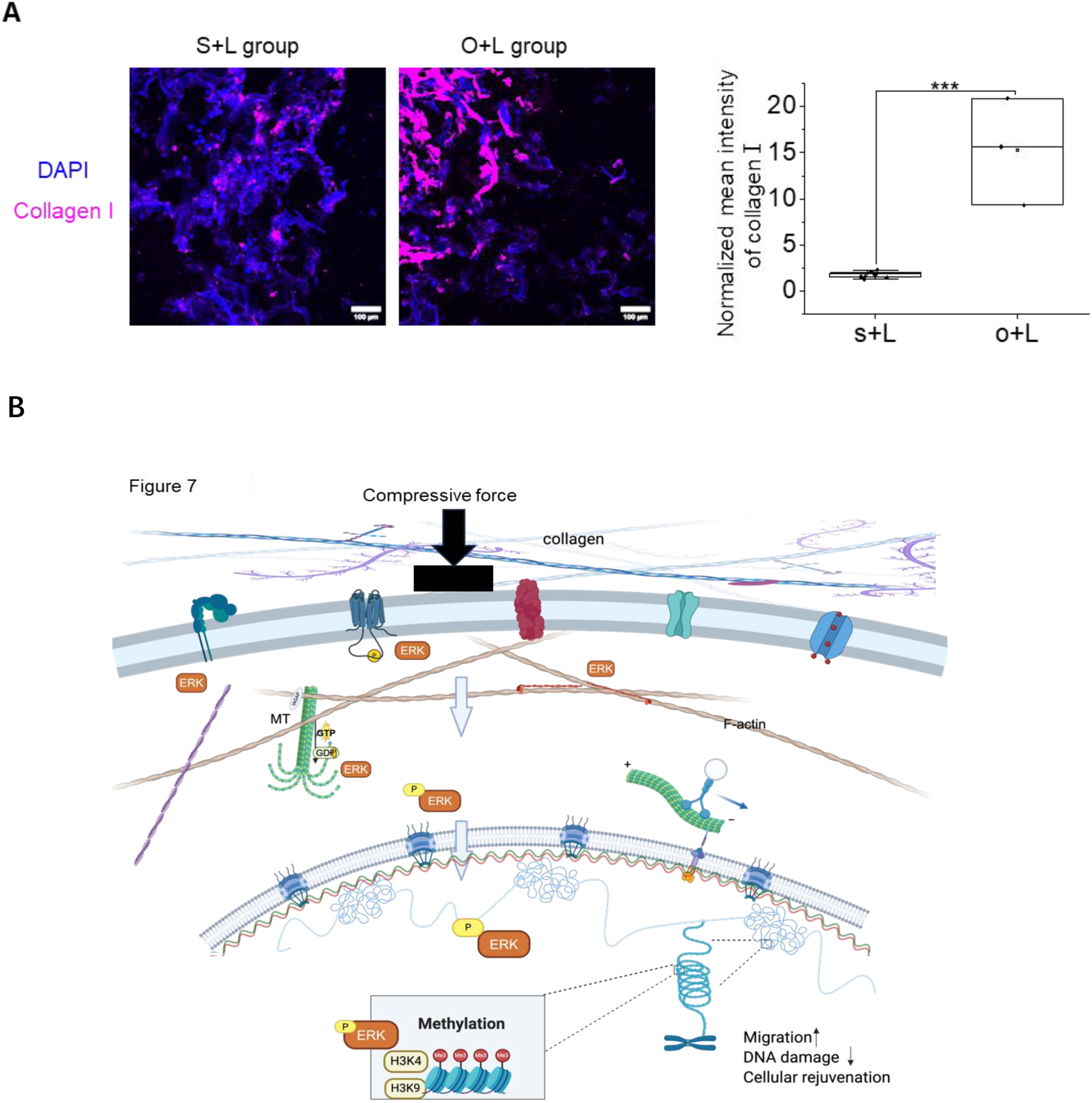
Implanted reprogrammed single cell and spheroids show activation properties under compressive force in an FT AGED skin model. (A) Representative images (20× magnification, Nikon) of collagen I. (Scale bar, 100μm). Normalized intensity plots of collagen I at the cell-implanted regions from at least 3 replicates. S+L group: inject single cell under compressive force; O+L group: inject spheroids under compressive force. (B) Illustration of the mechanism of compressive force in cellular rejuvenation. Figure created with BioRender.com. MT: microtubule.

## Discussion

In summary, this paper presents a new approach for activating/rejuvenating aging dermal fibroblasts using compressive forces. We selected 5% and 15% strain as indicators of compressive force, based on the assumed Young’s modulus of collagen hydrogel around 100 Pa (Figure S1D) (24–26). This differs from other studies that use pressure as an index (10, 11, 13, 27, 28). Our previous work demonstrated that compressive force induces actin depolymerization and leads to transcriptional quiescence at 2D single cell level (13). However, in 3D, compressive forces have been shown to activate cancer cell migration (11). In this paper, we hypothesized that by forming spheroids of aged cells which lead to the depolymerization of cytoskeletal filaments, may trigger activation pathways upon the application of compressive forces.

In particular, cells and nuclei adopt a relatively rounder morphology, compared to polarized single cells, when clumped in the designed pattern (Figure S1C). In this configuration, they exhibit a soft, highly sensitive state, poised to respond to external cues and initiate cellular processes. When such cells experience external forces at a global level, the highly viscoelastic cytoskeletal organization can activate cytoskeletal remodeling pathways (21, 29). This process entails the dynamic reorganization of actin filaments, microtubules, and intermediate filaments, forming networks that serve as mechanotransducers. These networks could facilitate the direct transmission of mechanical signals from the extracellular environment to the nucleus via the linker of nucleoskeleton and cytoskeleton (LINC) complex, ultimately influencing downstream gene transcription (8, 9). Conversely, the nucleus, which exhibits active rheological properties, can function as the primary mechano-sensor when subjected to sudden compressive forces (30). In our model, we observed a clear increase in cell contractility, as indicated by elevated levels of phosphorylated myosin light chain (pMLC). Additionally, we noted a complex reorganization of the microtubule network, accompanied by an increase in Lamin B expression and enhanced chromatin remodeling marked by increased levels of H3K9me3 and HP1a, while H3K4me3 decreased. These findings suggest that mechanical forces trigger cytoskeleton reorganization and chromatin remodeling.

Previous study has demonstrated that mechanical forces increase ɑSMA expression and its incorporation into actin filaments by activating two distinct signaling pathways: Rho/Serum Response Factor (SRF) and Mitogen-Activated Protein Kinase p38 (MAPK p38)/SRF (31). Consistent with these findings, our results indicate an increase in αSMA expression under compressive loading conditions, as shown in Figure 1B. As cells age, it is commonly observed that H3K9me3 levels decrease while H3K4me3 levels increase, accompanied by decreased HP1a levels and reduced LaminB expression (1). Surprisingly, in our novel model, we observed a converse trend. These findings suggest that the clumped HDFs (spheroids), upon sensing compressive forces, initiate fibroblast activation pathways including their potential rejuvenation, a notion further supported by our beta-galactosidase staining results.

Previous research has elucidated the role of mechanical compression in regulating cancer cell migration through the MEK1/ERK1 signaling pathway (12). In this study, compressive force enhanced cell migration, but whether this migration is driven by microtubules is unknown. During migration, cells extend filopodial or lamellipodial protrusions, form focal adhesions, and undergo cytoskeletal reorganization to generate force (32, 33). ERK phosphorylates key proteins involved in these processes, including microtubule-associated proteins (MAPs), focal adhesion kinase (FAK), calpain, and myosin light chain kinase (MLCK), thereby providing essential cues for cell migration (34–37). Additionally, phosphorylated ERK functions as a mechanosensory transcription factor capable of shuttling between the nucleus and cytosol to regulate gene expression, thereby exerting influence over cell migration dynamics (38). In our study, we demonstrated the significance of the force transmission pathway between the cytoskeleton and nucleus, as well as ERK signaling, in influencing cell migration. Employing a targeted approach, we conducted small-scale inhibitor screening and utilized a dominant-negative (DN) KASH construct to dissect the specific pathways involved. Our findings revealed that inhibitors targeting the cytoskeleton, including Blebbistatin, Cytochalasin D, Latrunculin A, Nocodazole, and Withaferin A, significantly impeded cell migration (Figure 3B-E). This underscores the critical role of the cytoskeleton in mediating force transmission essential for cell motility. Moreover, disruption of the LINC complex by the DN KASH (Figure 3B-E) construct corroborated these observations, further emphasizing the importance of the cytoskeleton-nucleus connections in governing cell migration dynamics. Inhibition of ERK1/ERK2 with PD98059 resulted in a notable decrease in cell migration distance. This highlights the pivotal role of ERK signaling in cell migration. In contrast, inhibition of FAK with PF-573228 had minimal effects on cell migration, highlighting the differential roles of focal adhesion-related signaling pathways.

Upon activation, ERK undergoes phosphorylation, resulting in its transformed state, pERK, which then translocates into the nucleus to regulate histone modifications. While a previous study suggested that ERK activation increases H3K4me3 levels in 2D culture conditions (39), interestingly in our 3D model with compressive forces, inhibition of ERK led to an increase in H3K4me3. We propose a possible mechanistic explanation for this observation: under compressive force conditions, perturbation of ERK affects cytoskeletal structure, resulting in reduced cytoskeletal formation and increased softness of the nucleus, as indicated by decreased laminB levels. This could result in higher levels of both H3K4me3 (a euchromatin marker) and H3K9me3 (a heterochromatin marker), reflecting a global change in chromatin architecture in response to ERK perturbation under compressive force conditions.

Our RNA-seq data revealed that under compressive force conditions, the expression of genes such as *INSR, BRAF, EGF, and EGFR* increased. These genes are upstream regulators of the ERK pathway, which initiates ERK signaling cascades in response to various extracellular stimuli (36). Additionally, the expression of *RPS6KA1*, also known as *RSK1*, increased. *RPS6KA1* is a downstream effector of the ERK pathway and is involved in mediating cellular responses to ERK activation. Several studies have highlighted the role of mechanical force in regulating key signaling molecules such as the insulin receptor, *BRAF*, and *EGFR* (40–42). Our study presents novel findings demonstrating that mechanical force activates aged dermal human fibroblasts through the ERK signaling pathway. As mechanosensory genes, we observed an upregulation of *YAP1*, *TRPV4*, and *PIEZO2* gene expression levels under 2x load conditions compared to the unloaded state, while *PIEZO1* was downregulated under 2x load conditions. Intriguingly, when the ERK inhibitor PD98059 was applied, *TRPV4* and *PIEZO1* gene expression levels were further inhibited, indicating a potential link between ERK and *TRPV4/PIEZO1* in our model. Consistently, previous research has demonstrated that mechanical forces can regulate *TRPV4* and *PIEZOs* (43). Moreover, another study has shown that mechanical force can activate TRPV4, subsequently leading to the induction of the ERK signaling pathway (44). These findings align with our observations and support the notion of a mechanistic relationship between mechanical force, *TRPV4/PIEZO1*, and activation of ERK signaling pathway. Upon inhibiting ERK, we observed the upregulation of *CLASP2*, *MAP2*, *MAPT*, and *HSPA1A* genes, which are known to be involved in microtubule dynamics. This suggests the presence of a compensatory response or feedback mechanism triggered by the inhibition of ERK signaling, highlighting the intricate interplay between mechanical force, ERK signaling, and cellular responses related to microtubule dynamics.

We observed a reduction in DNA damage under compressive force conditions, as indicated by decreased γH2AX immunostaining, suggesting enhanced DNA repair mechanisms. Our RNA sequencing data revealed the involvement of the Fanconi anemia pathway, known for anti-oxidative stress (45). This reduction in DNA damage aligns with the activation or rejuvenation required for aging cells. Previous study demonstrated decreased DNA damage under load conditions attributed to heterochromatin organization and low levels of H3K9me3 (16).

However, our results showed an upregulation of H3K9me3, potentially due to differences in cell culture models (2D vs. 3D). Notably, we observed no change in nucleus stiffness, as indicated by Lamin A/C immunostaining data. In our RNA sequencing data, rejuvenation-related genes such as *MBD2*, *KL*, *SIRT1*, *NFE2L2*, *LBR*, and *FOXO3* were upregulated. *MBD2* can reduce CpG methylation levels, delaying aging (46). *KL* has been shown to improve cognitive function, serving as a longevity factor (47). *SIRT* and *FOXO3* are known to slow cellular senescence (48), while *NFE2L2* acts as a transcription factor sensitive to reactive oxygen species (ROS) and nitric oxide (NO), induced by exercise, and protects cells against cytotoxic and oxidative damage (49). Upregulation of LBR can increase cell proliferation and suppress genomic instability, supporting the rejuvenating process (50). Upon load removal, cells maintained their migration behavior, initially exhibiting sustained high levels of αSMA followed by a decrease in our model, along with sustained increased expression of HP1a. The persistent expression of HP1a indicates that chromatin organization remains unchanged even after load removal within 5 days. The sustained expression of αSMA initially followed by a decrease in our model represents an intriguing finding, particularly considering that prolonged αSMA expression is associated with fibrosis. In the skin tissue model, after two days of incubation, the spheroid group exhibited higher secretion of collagen 1 compared to the single-cell group, highlighting the significance of compressive force induced tissue regeneration properties (Figure 6A).

Collectively, the activation of fibroblasts upon compressive force, cytoskeletal and chromatin remodeling, and transition from a mesenchymal to collective migration mode of aged HDFs may signify cellular rejuvenation (Figure 6B), crucial for both tissue regeneration and wound healing and could serve as a valuable platform for drug screening.

## Methods

### Fabrication of micropatterned PDMS stamps and microcontact printing

Polydimethylsiloxane (PDMS, SYLGRAD^TM^ 184 Silicone Elastomer Kit) elastomer is prepared by blending the base and curing agent at a 10:1 ratio. A typical quantity of 20-25g proves sufficient to entirely coat the custom wafer surface (1,800 μm^2^ rectangles, (aspect ratio 1:5), distance between rectangles is 500μm). PDMS is poured onto the wafer, and subjected to degases within a vacuum chamber for 30 minutes until the absence of air bubbles on the surface is achieved. Subsequently, the curing process is initiated at a temperature of 60°C for a duration of 3 hours. Following the cooling phase, the PDMS material is carefully detached from the substrate using a pair of tweezers. It is then sectioned into a round 1cm^2^ square pieces and stored within clean containers to avert the accumulation of particulate matter. The PDMS surface is activated by a Plasma machine (Henniker Plasma, HPT-200). Briefly, O2 gas was used by exposing the stamps to it for 1.5 min at 75% power, with pressure 0.4mbar. A mixture solution was prepared which includes fibronectin (MERK, F1141) and protein labeling kit (invitrogen, A20170A) in PBS at a concentration of 10% and 3%, respectively. For microcontact printing (mCP), a 10ul fibronectin mixture solution is applied onto the PDMS surface and observed under microscope for appropriate drying. After the fibronectin deposited on PDMS was dried, it was then stamped onto the un-coated IBIDI dishes (ididi, µ-Dish 35 mm, high, uncoated Cat.No:81151) and pressed gently using tweezers after which the PDMS is then carefully lifted. Finally these imprinted micro patterns were observed under EVOS fluorescence microscope..After careful selection of the dishes, they were passivated using 0.2% pluronic acid (Sigma, P2443) for 10 min followed by washing with PBS three times before seeding the cell.

### Cell culture

GM08401 (75 years old) and GM09503 (10 years old) healthy human dermal fibroblast cells (male origin) were obtained from the NIGMS Human Genetic Cell Repository at the Coriell Institute for Medical Research. The HDFs are cultured in MEM (Gibco, 11090-081) with 15% FBS (Thermo Fisher, 16141079), 1% P/S (Penicillin and Streptomycin) (PAN BIOTECH, P06-07300), 1% Glutamax (100x, Gibco, 35050-038) and 1% NEAA (100x, Gibco, 11140-035) under 5% CO2 and 37 °C. HEK293T cells (gift from Dr. Deborah Walter) were cultured in high-glucose DMEM supplemented (BioConcept, 1-26F03-1) with 10% (v/v) fetal bovine serum (Dominigue Dutscher, S1900-500B) and 1% P/S.

### Application of static compressive force in 3D culture model

70,000 old HDF cells were seeded on fibronectin-coated micropatterns in an ibidi dish overnight to form spheroids. 1mg/ml Collagen gel mixture was prepared from Collagen type I from rat tail (Gibco, A1048301) according to the manufacturer protocol. 400ul of 1mg/ml collagen hydrogel was applied on top of the spheroid and allowed for the polymerization for 1 hour in the incubator at 37 °C. After 1h incubation metal ring or glass ring was placed on top of collagen gel to avoid shrinkage during prolonged culture conditions. Glass coverslip (VWR, 631-1577, 12mm round) was used for compressive force and placed carefully without disrupting the gel inside the ring. For 5% applied compressive force 3 stacked coverslips and for 15% 7 stacked coverslips were used (Figure S1D).

### Immunostaining

Collagen hydrogel samples were fixed with 4% paraformaldehyde (Merk, F8775-25ML) for 1h and the coverslip, used to apply compressive forces, was then removed carefully. Gels are then washed with 100mM glycine (Roth, Nr.3790.3) three times to prevent excess fixation. Permeabilization was done for 20 minutes with 0.8% Triton X-100 for γH2AX and ERK staining and 0.5%Triton X-100 was used for the rest of the markers. This was followed by washing with 100mM glycine for three times. Samples were then blocked with 10% NGS (Abcam, ab7481) in wash buffer PBS (PanReac Applichem, A0964 9050) containing 0.2%Triton and 0.2% Tween20 (Sigma, SLBZ8563) for three hours at room temperature (For γH2AX and ERK staining PBS containing 0.3%Triton and 0.2% Tween20 wash buffer used for blocking). Primary antibody staining was done with 10% goat serum in the wash buffer for two days at 4°C. Next day, the gels were washed with a buffer (PBS containing 0.2%Triton and 0.2% tween20) for 10-15 minutes each wash three times. Secondary antibody staining 1:300 dilution was done in 5% goat serum in the wash buffer and incubated for three hours at room temperature. Followed by washing with a wash buffer for 15 minutes once and then twice with PBS 15 minutes each. DAPI (Thermo Fisher Scientific, R37605), nucleus stain, and ActinGreen (Thermo Fisher Scientific, R37110), actin stain, was incubated in PBS overnight at 4°C and washed three times with PBS. Finally, 100 ul of PBS was added and imaged using Nikon confocal imaging system. Antibodies used in this paper are listed in Supplementary Table 2. For the beta-galactosidase assay, CellEvent™ Senescence Green Flow Cytometry Assay Kit (Invitrogen, C10840) was used as per manufacturer protocol and confocal images were captured. DRAQ7 (Biolegend, 424001) was used to discriminate between live/dead cells.

### Drug treatment

All the drugs with specific concentrations used in our assays are mentioned in Figure S1E and supplementary table 3. 1 ml medium with the respective drug concentrations were added to the overnight spheroids and incubated for 1h, before covering it with the collagen gel. After 1h of gel polymerisation, 2ml of new complete medium with respective concentration of drug was added and incubated for 2 days. Finally, these gels were processed for immunostaining and imaging.

### Real-time PCR assay

For RNA purification, at least 10 gels from aged fibroblasts (with and without load) were used. Single cells were isolated from these gels using collagenase at a concentration of 2 mg/ml (Merk, C0130) and incubated at 37^0^C for 30 minutes. After centrifugation at 1000 rpm for 4 minutes, the supernatant was removed and pellet was collected to be processed for RNA isolation using RNeasy Plus Micro Kit (QIAGEN, 74034). cDNA was prepared using iScript™ cDNA Synthesis Kit (BIO-RAD, 1708890). Real-time PCR was done using Sso advanced SYBER mix (BIO-RAD, 1725274). Relative fold change was calculated with the 2^-ΔΔCT method using GAPDH as a housekeeping gene for normalization. All primers used in this study are shown in Supplementary Table 1.

### RNA seq analysis

RNA library preparation and sequencing was performed at Genomics Facility, ETH Zurich in Basel. Unload group, 1XL group, and 2XL group in triplicates were performed by NovaSeq S4 PE 2×101bp. The PD98059 group in triplicates was performed by NextSeq PE 2×38bp. Standard pipelines such as DEseq were used for RNA seq analysis (51, 52). To summarize, the paired-end reads were aligned to the human genome GRCh38.84 from UCSC. Reference genomic indexes using the HISAT2 sequence-alignment tool (version 2.2.1) was used. The cloud indexes (grch38_trans) for HISAT2, was accessed on June 25th, 2020 from https://registry.opendata.aws/jhu-indexes. Combining reads from four technical replicates for each biological sample served as the input for HISAT2, utilizing default parameters. Subsequently, single aligned reads were enumerated using htseq-count (version 1.99.2). The counts for all expressed genes were then employed for the differential expression analysis and analyzed using DESeq2 (Version 1.36.0). Batch information was incorporated into the DESeq2 design formula. Differentially expressed genes were identified based on adjusted P values (Benjamini–Hochberg) below 0.1 false discovery rate (FDR) and fold change above or below 2. Enrichment analysis was performed using ShinyGO (version 0.80). Python script was utilized for generating heat maps that compare gene expression across different biological conditions, based on DESeq2 normalized counts.

### In vitro 3D reconstructed skin models under compressive force

In this study, we used the Phenion FT AGED skin model as a substitute for aged human skin. This model includes senescent fibroblasts, reduced ECM proteins (such as collagen and elastin), and elevated MMP secretion due to treatment with mitomycin C. We divided the samples into six groups, as shown in Figure S15A. Single-cell injection refers to the collection of cells from 2D cell cultures. Spheroid injection involves collecting cells following the method described above. The elastic modulus of the reconstructed skin model was approximately 7kPa (53), and deformation of up to 12.6% was achieved under compressive force, as indicated in Figure S15B. Cells were injected at three different points, with a concentration of ∼70,000 cells per point and an injection volume of 50 µL. The injection sites were located approximately 2 mm from the center, and a wound was made at one site to indicate the direction (Figure S15C). The reconstructed skin models were cultured in a specific Air-Liquid Interface Culture Medium provided by the supplier.

### Cryo-sectioning and immunofluorescence of tissue sections

After two days of culture, the tissues were placed in cryomolds and embedded in OCT medium (Leica Biosystem, 14020108926). Samples were cryo-sectioned at a thickness of 20 µm at −15 °C using a cryo-microtome and stored at −80 °C until staining. For immunostaining, the tissue sections were fixed in pre-cooled acetone (VWR, 20063.296) for 15 minutes at −20 °C. After air-drying for 5 minutes, a PAP pen (Sigma-Aldrich) was used to encircle the tissue.

Sections were then blocked with 10% goat serum for 1 hour. Subsequently, the samples were incubated with primary antibodies diluted in 1% BSA and 0.3% Triton-X 100 in PBS overnight at 4 °C. After three washes in PBS (5 minutes each), the sections were incubated with secondary antibodies diluted in 1% BSA and 0.3% Triton-X 100 in PBS overnight at 4 °C. Following another three washes in PBS, the samples were stained with Hoechst 33342 in PBS (one drop per 1ml) for 1 hour at room temperature. Finally, the sections were mounted with ProLong Gold Antifade Reagent (Thermo Fisher Scientific), covered with a coverslip, and sealed at the edges with a thin layer of nail polish. The slides were stored at 4 °C until imaging.

### Image acquisition and analysis

All confocal images were obtained using the Nikon confocal ti2 imaging system. Briefly, collagen hydrogel was imaged using a 40X oil immersion objective NA 1.25 or 60 X oil immersion objective NA 1.4. All bright-field images in this study were acquired using EVOS M5000 (Thermo Fisher Scientific) and slide scanner (sysmex) for skin tissue. For the analysis of mean intensity and the γH2AX foci number, Fiji image tool was used. For nuclear marker analysis, DAPI channel was used to generate the mask, whereas for cytosolic protein, either the actin or protein channel was used for mask generation. Nuclear and chromatin features analysis was done using the code from previously published paper (54). The importance of each attribute was measured by Relief F/Gini/Gain ratio methods via orange software. Internuclear pairwise distance (IPD) analysis was performed using R package dist. Labeled images were processed using the StarDist2D plugin in Fiji. Microtubule meshwork generation and directionality histograms analysis were conducted by SOAX software and Fiji software with plugin directionality. For skin tissue samples, Fiji software was used to measure the mean fluorescence intensity and to count cell numbers using the StarDist2D plugin. Normalization was calculated as the ratio of the mean intensity of the immunofluorescence-positive injected area to the cell number.

### Statistical analysis

All plots and statistical analysis were performed with Origin 2024. For box-and-whisker plots: The box represents the interquartile range (IQR), encompassing the middle 50% of the data. The bottom of the box marks the first quartile (25th percentile), and the top marks the third quartile (75th percentile). The line inside the box indicates the median (50th percentile). The whiskers extend to the smallest and largest values within 1.5 times the IQR, while outliers are represented by asterisks. Unpaired, two-tailed student-t test was used to compare two groups. One-way ANOVA (Tukey test) was employed to compare groups comprising more than two.

## Data and materials availability

All codes used in this paper are available from the corresponding author upon request. All illustration graphs shown in this study were created by Biorender.com. All data are available in the main text or the supplementary materials.

## Acknowledgments

We thank GVS group members for their comments on the manuscript and in particular Drs. Nicholas Lawler and Yagyik Goswami for critical reading of the manuscript. This work was supported by the Swiss National Science Foundation grant 310030_208046; China Scholarship Council (Grant Number: 202008440471).

## Notes

### Competing Interest Statement

The authors have declared no competing interest.

## References

1. C. López-Otín, M. A. Blasco, L. Partridge, M. Serrano, G. Kroemer, Hallmarks of aging: An expanding universe. Cell 186, 243–278 (2023).

2. M. V. Plikus, et al., Fibroblasts: Origins, definitions, and functions in health and disease. Cell 184, 3852–3872 (2021).

3. S. Knoedler, et al., Fibroblasts – the cellular choreographers of wound healing. Front. Immunol. 14, 1233800 (2023).

4. K. Takahashi, S. Yamanaka, Induction of Pluripotent Stem Cells from Mouse Embryonic and Adult Fibroblast Cultures by Defined Factors. Cell 126, 663–676 (2006).

5. B. Roy, et al., Laterally confined growth of cells induces nuclear reprogramming in the absence of exogenous biochemical factors. Proc. Natl. Acad. Sci. 115 (2018).

6. W. Zhang, J. Qu, G.-H. Liu, J. C. I. Belmonte, The ageing epigenome and its rejuvenation. Nat. Rev. Mol. Cell Biol. 21, 137–150 (2020).

7. C. C. DuFort, M. J. Paszek, V. M. Weaver, Balancing forces: architectural control of mechanotransduction. Nat. Rev. Mol. Cell Biol. 12, 308–319 (2011).

8. C. Uhler, G. V. Shivashankar, Regulation of genome organization and gene expression by nuclear mechanotransduction. Nat. Rev. Mol. Cell Biol. 18, 717–727 (2017).

9. G. V. Shivashankar, Mechanical forces and the 3D genome. Curr. Opin. Struct. Biol. 83, 102728 (2023).

10. Y. Cui, et al., Cyclic stretching of soft substrates induces spreading and growth. Nat. Commun. 6, 6333 (2015).

11. J. M. Tse, et al., Mechanical compression drives cancer cells toward invasive phenotype. Proc. Natl. Acad. Sci. 109, 911–916 (2012).

12. M. Kalli, et al., Mechanical Compression Regulates Brain Cancer Cell Migration Through MEK1/Erk1 Pathway Activation and GDF15 Expression. Front. Oncol. 9, 992 (2019).

13. K. Damodaran, et al., Compressive force induces reversible chromatin condensation and cell geometry–dependent transcriptional response. Mol. Biol. Cell 29, 3039–3051 (2018).

14. N. Naetar, S. Ferraioli, R. Foisner, Lamins in the nuclear interior – life outside the lamina. J. Cell Sci. 130, 2087–2096 (2017).

15. F. Napoletano, et al., The prolyl-isomerase PIN1 is essential for nuclear Lamin-B structure and function and protects heterochromatin under mechanical stress. Cell Rep. 36, 109694 (2021).

16. M. M. Nava, et al., Heterochromatin-Driven Nuclear Softening Protects the Genome against Mechanical Stress-Induced Damage. Cell 181, 800–817.e22 (2020).

17. T. Montavon, et al., Complete loss of H3K9 methylation dissolves mouse heterochromatin organization. Nat. Commun. 12, 4359 (2021).

18. X. Liu, et al., Distinct features of H3K4me3 and H3K27me3 chromatin domains in pre-implantation embryos. Nature 537, 558–562 (2016).

19. W. Zeng, A. R. Ball Jr, K. Yokomori, HP1: Heterochromatin binding proteins working the genome. Epigenetics 5, 287–292 (2010).

20. D. M. Rosenbaum, S. G. F. Rasmussen, B. K. Kobilka, The structure and function of G-protein-coupled receptors. Nature 459, 356–363 (2009).

21. N. Wang, J. D. Tytell, D. E. Ingber, Mechanotransduction at a distance: mechanically coupling the extracellular matrix with the nucleus. Nat. Rev. Mol. Cell Biol. 10, 75–82 (2009).

22. Y. Pan, et al., Mechanosensor Piezo1 mediates bimodal patterns of intracellular calcium and FAK signaling. EMBO J. 41, e111799 (2022).

23. F. K. Noubissi, et al., Detection and quantification of γ-H2AX using a dissociation enhanced lanthanide fluorescence immunoassay. Sci. Rep. 11, 8945 (2021).

24. T. Jin, L. Li, R. C. Siow, K.-K. Liu, Collagen matrix stiffness influences fibroblast contraction force. Biomed. Phys. Eng. Express 2, 047002 (2016).

25. C. S. Vidmar, M. Bazzi, V. K. Lai, Computational and experimental comparison on the effects of flow-induced compression on the permeability of collagen gels. J. Mech. Behav. Biomed. Mater. 128, 105107 (2022).

26. C. R. I. Lam, et al., A 3D Biomimetic Model of Tissue Stiffness Interface for Cancer Drug Testing. Mol. Pharm. 11, 2016–2021 (2014).

27. Y. Tang, Y. Fan, Q. Luo, G. Song, Pressure Loading Induces DNA Damage in Human Hepatocyte Line L02 Cells via the ERK1/2–Dicer Signaling Pathway. Int. J. Mol. Sci. 23, 5342 (2022).

28. T. Kanazawa, et al., Biological Responses of Three-Dimensional Cultured Fibroblasts by Sustained Compressive Loading Include Apoptosis and Survival Activity. PLoS ONE 9, e104676 (2014).

29. I. Andreu, et al., The force loading rate drives cell mechanosensing through both reinforcement and cytoskeletal softening. Nat. Commun. 12, 4229 (2021).

30. Y. Song, J. Soto, B. Chen, L. Yang, S. Li, Cell engineering: Biophysical regulation of the nucleus. Biomaterials 234, 119743 (2020).

31. J. Wang, R. Zohar, C. A. McCulloch, Multiple roles of α-smooth muscle actin in mechanotransduction. Exp. Cell Res. 312, 205–214 (2006).

32. C. Le Clainche, M.-F. Carlier, Regulation of Actin Assembly Associated With Protrusion and Adhesion in Cell Migration. Physiol. Rev. 88, 489–513 (2008).

33. K. M. Yamada, M. Sixt, Mechanisms of 3D cell migration. Nat. Rev. Mol. Cell Biol. 20, 738–752 (2019).

34. V. J. Fincham, Active ERK/MAP kinase is targeted to newly forming cell-matrix adhesions by integrin engagement and v-Src. EMBO J. 19, 2911– 2923 (2000).

35. R. E. Harrison, E. A. Turley, Active Erk Regulates Microtubule Stability in H-ras-Transformed Cells. Neoplasia 3, 385–394 (2001).

36. H. Lavoie, J. Gagnon, M. Therrien, ERK signalling: a master regulator of cell behaviour, life and fate. Nat. Rev. Mol. Cell Biol. 21, 607–632 (2020).

37. C. Huang, K. Jacobson, M. D. Schaller, MAP kinases and cell migration. J. Cell Sci. 117, 4619–4628 (2004).

38. D. A. Berti, R. Seger, “The Nuclear Translocation of ERK” in ERK Signaling, Methods in Molecular Biology., G. Jimenez, Ed. (Springer New York, 2017), pp. 175–194.

39. C. Esnault, et al., ERK-Induced Activation of TCF Family of SRF Cofactors Initiates a Chromatin Modification Cascade Associated with Transcription. Mol. Cell 65, 1081–1095.e5 (2017).

40. J. Kim, D. Bilder, T. P. Neufeld, Mechanical stress regulates insulin sensitivity through integrin-dependent control of insulin receptor localization. Genes Dev. 32, 156–164 (2018).

41. A. Hollósi, K. Pászty, M. Kellermayer, G. Charras, A. Varga, BRAF Modulates Stretch-Induced Intercellular Gap Formation through Localized Actin Reorganization. Int. J. Mol. Sci. 22, 8989 (2021).

42. B. Sullivan, et al., Mechanical disruption of E-cadherin complexes with epidermal growth factor receptor actuates growth factor–dependent signaling. Proc. Natl. Acad. Sci. 119, e2100679119 (2022).

43. M. Zhang, N. Meng, X. Wang, W. Chen, Q. Zhang, TRPV4 and PIEZO Channels Mediate the Mechanosensing of Chondrocytes to the Biomechanical Microenvironment. Membranes 12, 237 (2022).

44. P. S. Nayak, et al., Mechanotransduction via TRPV4 regulates inflammation and differentiation in fetal mouse distal lung epithelial cells. Respir. Res. 16, 60 (2015).

45. W. Du, Z. Adam, R. Rani, X. Zhang, Q. Pang, Oxidative Stress in Fanconi Anemia Hematopoiesis and Disease Progression. Antioxid. Redox Signal. 10, 1909–1921 (2008).

46. A. A. Johnson, et al., The Role of DNA Methylation in Aging, Rejuvenation, and Age-Related Disease. Rejuvenation Res. 15, 483–494 (2012).

47. C. Park, et al., Platelet factors are induced by longevity factor klotho and enhance cognition in young and aging mice. Nat. Aging 3, 1067–1078 (2023).

48. K. A. Nath, The role of Sirt1 in renal rejuvenation and resistance to stress. J. Clin. Invest. 120, 1026–1028 (2010).

49. T. L. Merry, M. Ristow, Nuclear factor erythroid-derived 2-like 2 (NFE2L2, Nrf2) mediates exercise-induced mitochondrial biogenesis and the anti-oxidant response in mice. J. Physiol. 594, 5195–5207 (2016).

50. A. En, K. Takemoto, Y. Yamakami, K. Nakabayashi, M. Fujii, Upregulated expression of lamin B receptor increases cell proliferation and suppresses genomic instability: implications for cellular immortalization. FEBS J. febs.17113 (2024). 10.1111/febs.17113.

51. B. Roy, T. Pekec, L. Yuan, G. V. Shivashankar, Implanting mechanically reprogrammed fibroblasts for aged tissue regeneration and wound healing. Aging Cell 23, e14032 (2024).

52. M. I. Love, W. Huber, S. Anders, Moderated estimation of fold change and dispersion for RNA-seq data with DESeq2. Genome Biol. 15, 550 (2014).

53. D. Malhotra, et al., Linear viscoelastic and microstructural properties of native male human skin and in vitro 3D reconstructed skin models. J. Mech. Behav. Biomed. Mater. 90, 644–654 (2019).

54. S. Venkatachalapathy, D. S. Jokhun, M. Andhari, G. V. Shivashankar, Single cell imaging-based chromatin biomarkers for tumor progression. Sci. Rep. 11, 23041 (2021).

