## Supplementary File for "Compressive forces induce epigenetic activation of aged human dermal fibroblasts through ERK signaling pathway"

**Abstract**

Age-related changes in human dermal fibroblasts (HDFs) contribute to impaired wound healing and skin aging. While these changes result in altered mechanotransduction, the epigenetic basis of rejuvenating aging cells remains a significant challenge. This study investigates the effects of compressive forces on nuclear mechanotransduction and epigenetic rejuvenation in aged HDFs. Using a compressive force application model, the activation of HDFs through alpha-smooth muscle actin (α-SMA) is demonstrated. Sustained compressive forces induce significant epigenetic modifications, including chromatin remodeling and altered histone methylation patterns. These epigenetic changes correlate with enhanced cellular migration and rejuvenation. Small-scale drug screening identifies the extracellular signal-regulated kinase (ERK) signaling pathway as a key mediator of compression-induced epigenetic activation. Furthermore, implanting aged cell spheroids to an aged skin model and subjecting the tissue with compressing forces resulted in increased collagen I protein levels. Collectively, these findings demonstrate that applying compressive force to aged fibroblasts activates global epigenetic changes through the ERK signaling pathway, ultimately rejuvenating cellular functions with potential applications for wound healing and skin tissue regeneration.

### Methods

#### Lentiviral Transfection assay.

DN KASH GFP-KAN (gift from Prof. Brian Burke's lab) E. coli strain was cultured on a plate with 50 μg/ml kanamycin sulfate (GERBU, #1091.0250) overnight. One clone was chosen and transferred into 100 ml Luria Broth (LB) for overnight culture. The NucleoBond Xtra Midi Kit (Macherey-nagel, #740410.50) was used to purify the plasmid DNA. Based on the DN KASH sequence, we designed DN KASH primers (supplementary table 1) to expand the region of interest using the PCR method and purified it through agarose gel extraction.

pLV-EGFP/pPLV-mcherry (Addgene #36083, #Addgene 36084) were cut using AgeI HF/SalI HF, and the DN KASH PCR product was cut using AgeI HF/SalI HF/Dpnl. Finally, primer pLV-EGFP was selected to ligate with the DN KASH product. Transformation was carried out in E.Coli using the addition of 1ul purified plasmid DNA to E.coli cells by gentle tap and immediately placed on ice and incubated for 30 minutes. After incubation, the tubes were then placed in a 37°C water bath for 45 seconds and immediately placed in ice for 2 minutes. Followed by addition of the SOC (invotrogen, P/N 46-0700) medium and cultured in a 37°C bacterial shaker incubator for 1 hour with agitation of 250rpm. 100ul of Competent E. coli was spread on LB (Luria-Bertani) agar plate containing 100 mg/ml Ampicillin (GERBU, #1091.0250) and incubated at room temperature for 2 days. After incubation five clones were picked and cultured in 50 ml LB medium overnight in a 37°C bacterial shaker incubator with agitation of 250rpm. Next day plasmid DNA was purified from these five clones using NucleoBond Xtra Midi Kit (Macherey-nagel, #740588.250). Finally, the plasmid presence was verified through enzymatic digestion and run on agarose gel electrophoresis and plasmids were sequenced to verify the sequence (Supplementary List 2).

A total of 48 μg of DNA mixture was prepared, containing lentivirus plasmids in a ratio of 3:1:1:1. These plasmids include the lentiviral gene carrier pRRLSIN.cPPT.PGK-GFP.WPRE with RabGAP insert (Addgene plasmid #12252), pMD2.G (Addgene plasmid #12259, provided by Didier Trono), pMDLg/pRRE, and pRSV-Rev. HEK293T cells were used as packaging cells and further details were indicated by a previously established protocol (Xie et al., 2019). GM08401 fibroblast, which acted as host cell, was seeded in a 12-well plate at a concentration of $1x10^5$ cells per well. Different volumes of concentrated lentivirus suspension (100μl, 50μl, 25μl, 10μl) were tested to optimize conditions. Finally, 100μl and 50μl were chosen to transfect the host cell and transfected cells were propagated for 2 weeks before 3D gel application. The transfection efficiency was shown in Supplementary Figure S5A-C. The lentiviral construct design and transfection experiments were carried out with the help of Dr. Phillip Berger and Lulu Yang.

Supplementary figure 1

A

Metal ring

Glass ring

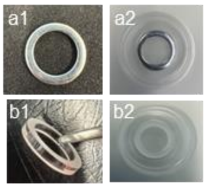


| | Outer diameter | Inner diameter | Height | Weight | Density |
|---|---|---|---|---|---|
| Metal ring | 20mm | 14mm | 2mm | 2g | 7.85g/cm$^3$ |
| Glass ring | 20mm | 15mm | 3mm | 0.88g | 2.5g/cm$^3$ |

B

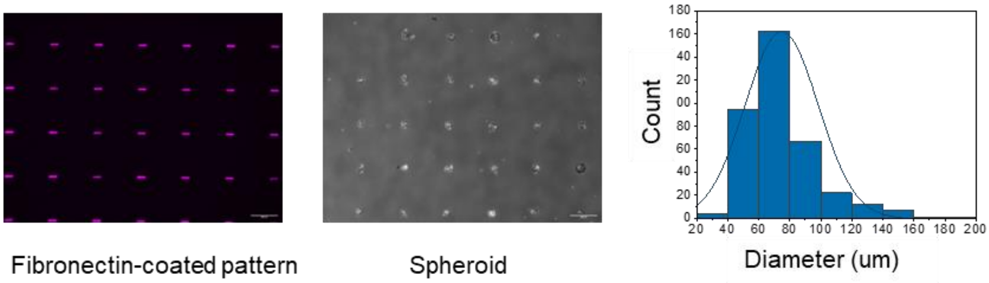


Fibronectin-coated pattern

Spheroid

C

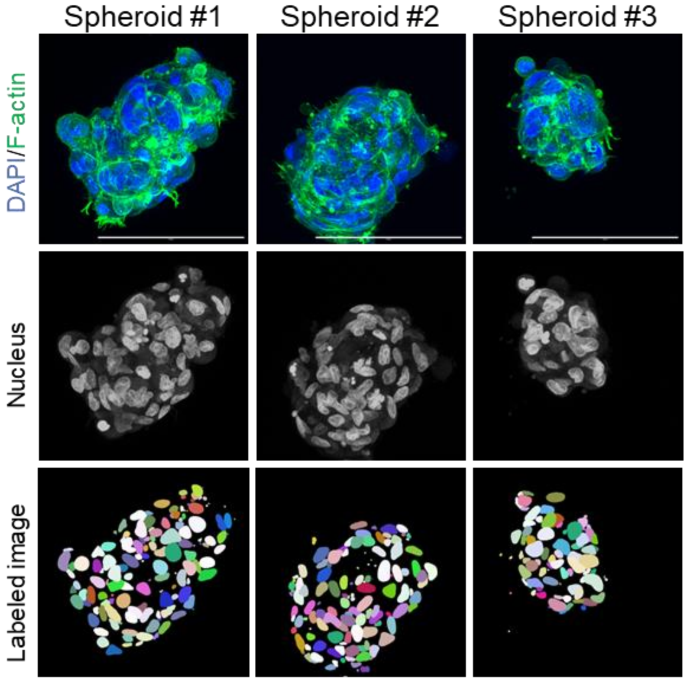


D

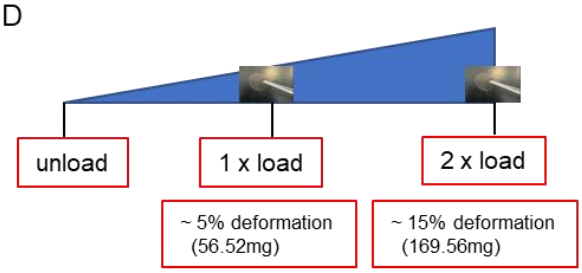


Coverslip weight in medium:

First, $Volumn = M/\rho = \frac{40mg}{2.5g/cm^3} = 1.6x10^{-8}m^3$

Then, $Weight = mg - buoyant\ force$

| | Diameter | Weight | Weight in medium |
|---|---|---|---|
| coverslip | 12mm | 40mg | 24mg |

### Supplementary figure 1

Young's modulus equation:

$$E = \frac{tensile\ stress}{extensional\ strain} = \frac{\sigma}{\varepsilon} = \frac{F/A_0}{\Delta L/L_0} = \frac{FL_0}{A_0\Delta L} = \frac{\frac{F}{A_0}}{\text{strain}} = \frac{\frac{mg}{A_0}}{strain}$$

where

$E$ is the Young's modulus (modulus of elasticity)

$F$ is the force exerted on an object under tension;

$A_0$ is the actual cross-sectional area through which the force is applied;

$\Delta L$ is the amount by which the length of the object

$L_0$ is the original length of the object.

| |
|---|
| Young modulus: around 100Pa (1mg/ml collagen) |
| Strain : 5% —15% |
| Area : $A = \pi r^2 = \pi(6mm)2$ |
| Weight : 56.52mg—169.56mg |
| Number of coverslip: 3 — 7 |

E

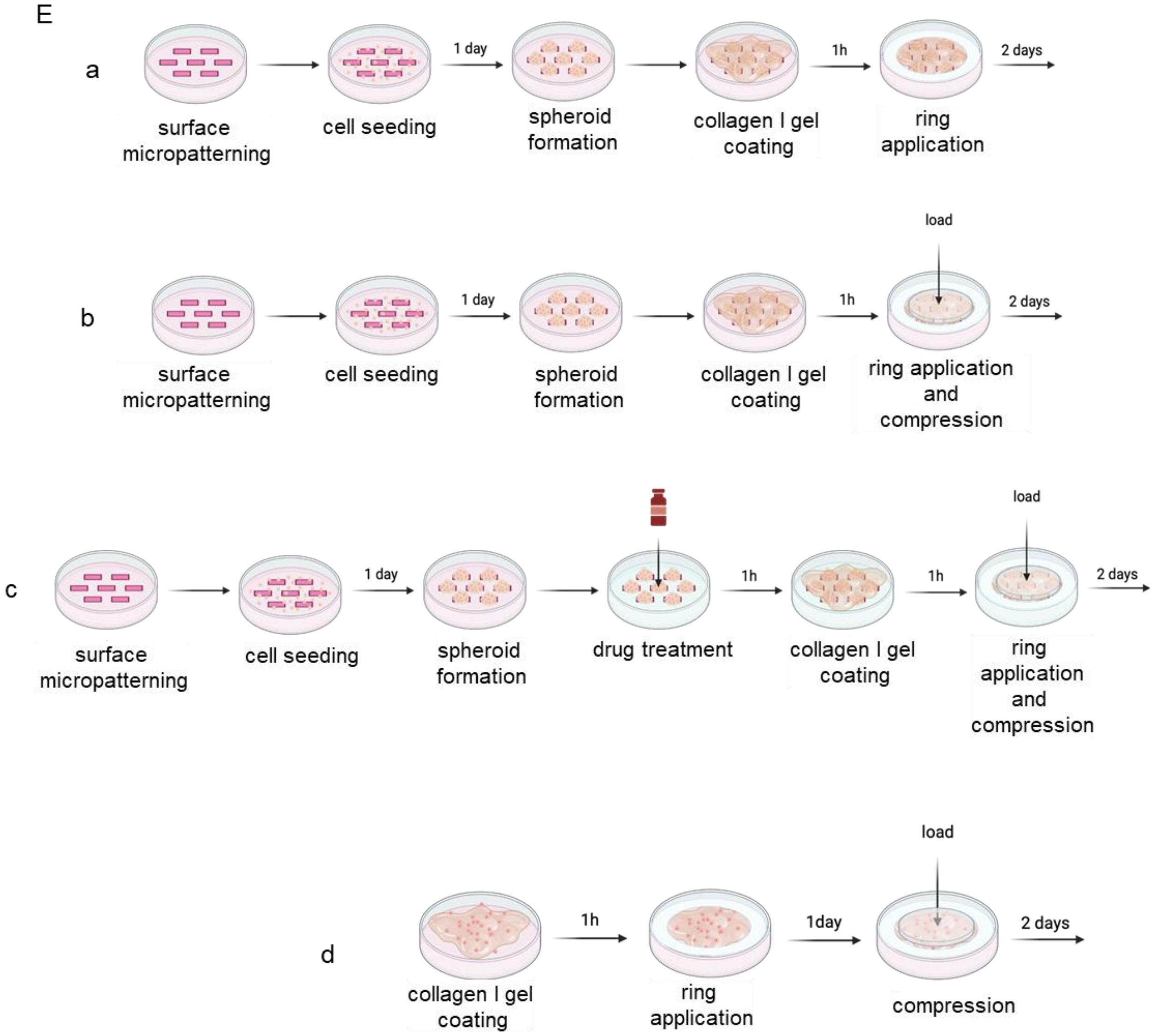

Supplementary Figure.1. Illustration of material properties, spheroid formation methods and compressive force calculation. (A) Photograph of metal ring and glass ring, and a table for their description. (B) Fibronectin-coated micropatterns of 1800 $\mu m^2$ area on IBIDI 35*μm* dish. (Scale bar, 300*μm*); The brightfield images of spheroid formation of fibroblasts cultured on micropatterns overnight. (Scale bar, 300*μm*); The distribution of spheroid diameter. (C) Representative confocal images of spheroid, 3D construct images and labeled image by StarDist2D, F-actin(green) and nucleus (blue). (Scale bar, 100*μm*). (D) The process of calculating the applied compressive forces. (E) Step by step process for sample preparation. (a) Sample preparation for the control group. (b) Sample preparation for load group. (c) Sample preparation for drug treatment. (d) single cell embedding in collagen hydrogel.

Supplementary figure 2

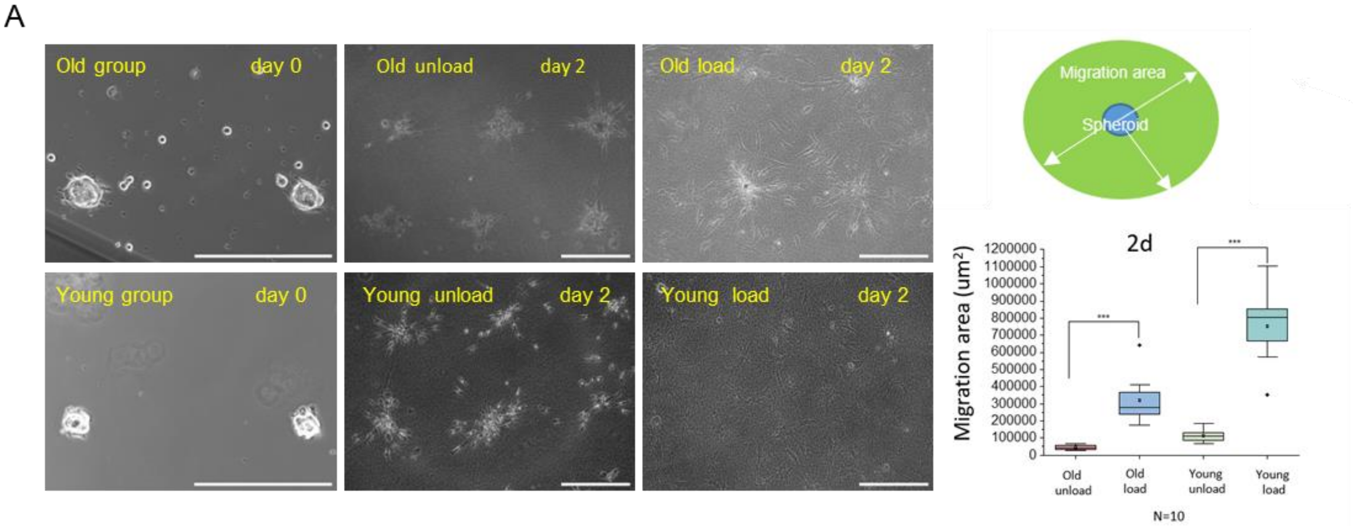


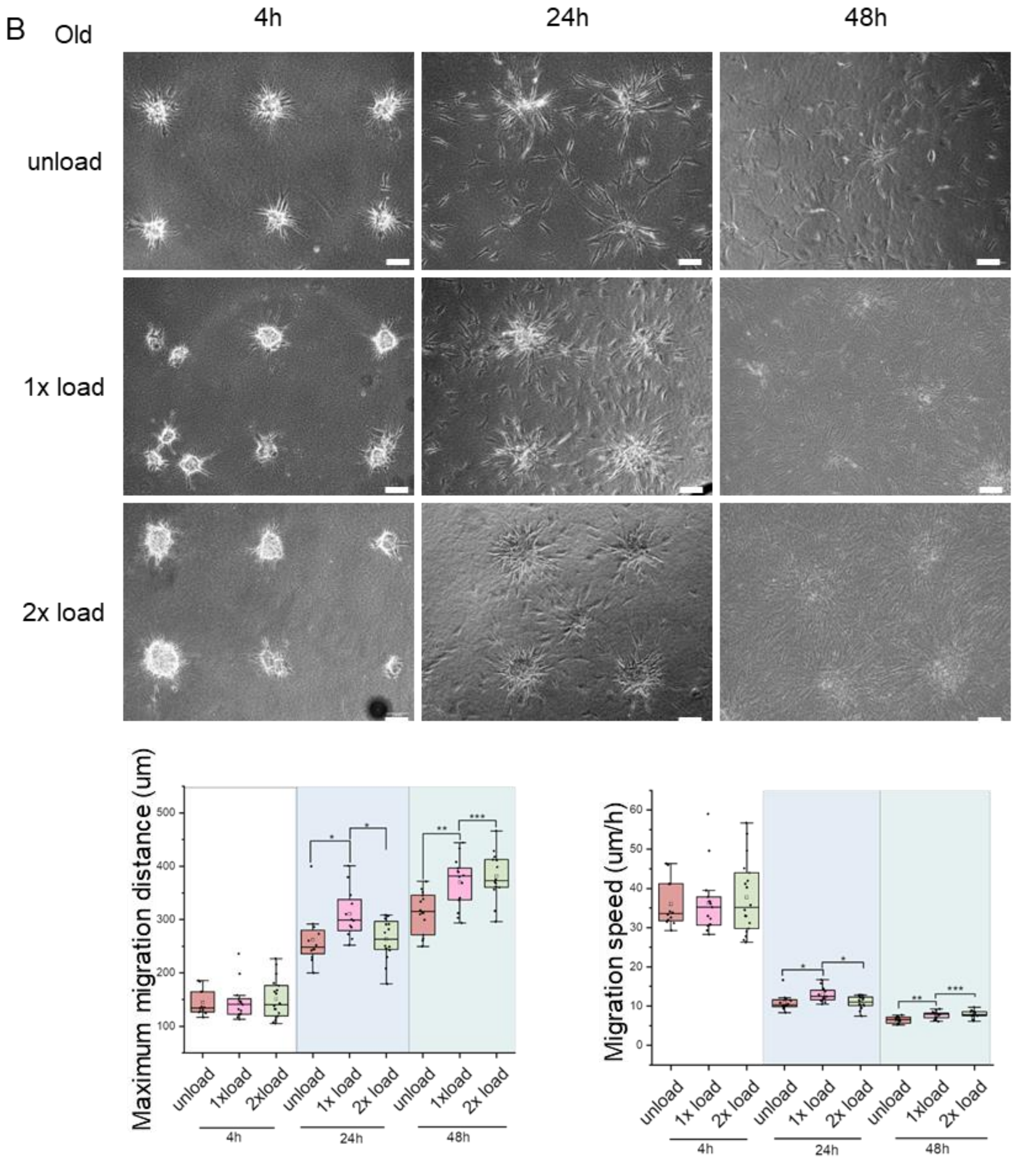

Supplementary figure 2

C

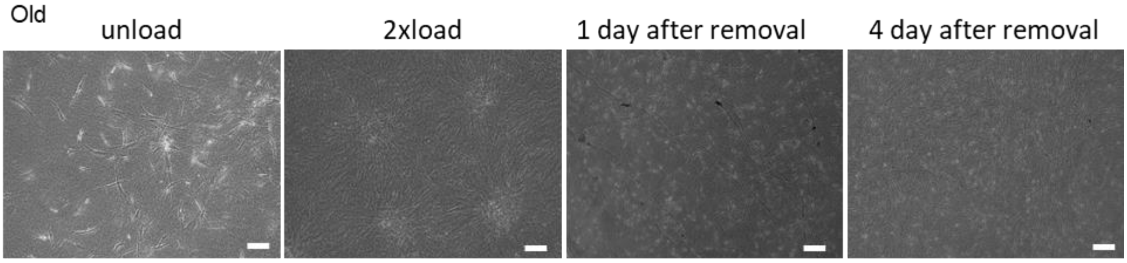


D

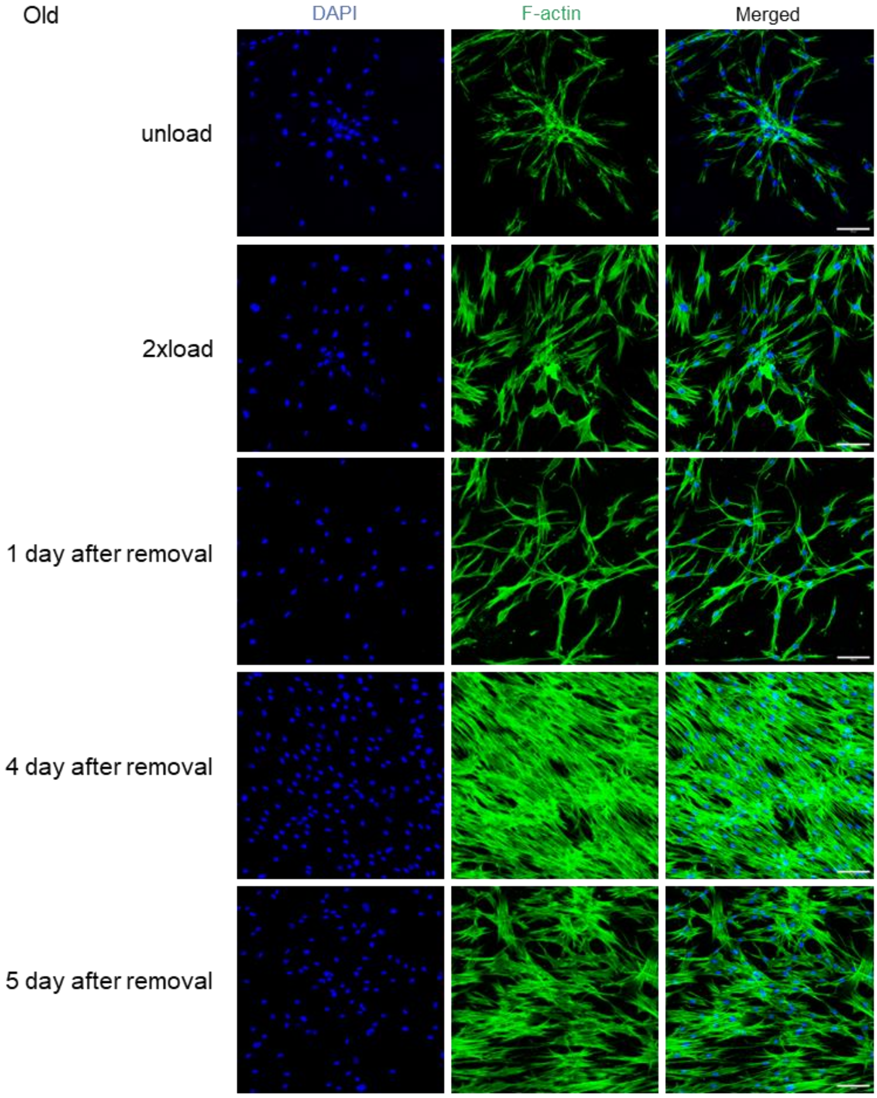


E

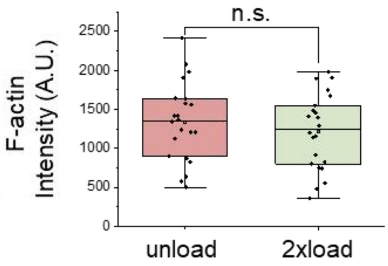

Supplementary Figure.2. Cell migration behavior and status check under mechanical loading and load removal conditions (metal ring and glass ring). (A) The brightfield images of GM08401 human (75 years old, old group) fibroblasts and GM09503 human (10 years old, young group) fibroblasts cultured for 2 days by using metal rings. (Scale bar, 300*μm*); Quantification data of migration area. (B) The brightfield images of GM08401 human (75 years old, old group) culture for 4h, 24h and 48h by using glass rings. (Scale bar, 100*μm*); Quantification data of maximum migration distance and migration speed. (C) The brightfield images of load removal. (Scale bar, 100*μm*). (D) Representative confocal images in different groups (unload, 2xload, remove for 1 day, remove for 4 days and remove for 5 days). GM08401 fibroblasts are used in this study. F-actin(green) and nucleus (blue). (Scale bar, 100*μm*). (E) Quantification data per image of F-actin in unload and 2xload group. *All the experiments were repeated at least three times independently with similar results. P values in Figure (A) were calculated by unpaired, two-tailed Student's t test. old unload group vs old load group; young unload group vs young load group. P values in Figure (B) were calculated by the one-way ANOVA method with Tukey's post hoc test at 4h, 24h and 48h separately. P values in Figure (E) were calculated by unpaired, two-tailed Student's t test. *P<0.05; **<0.01; ***P<0.001; No asterisks mean not significant.*

Supplementary figure 3

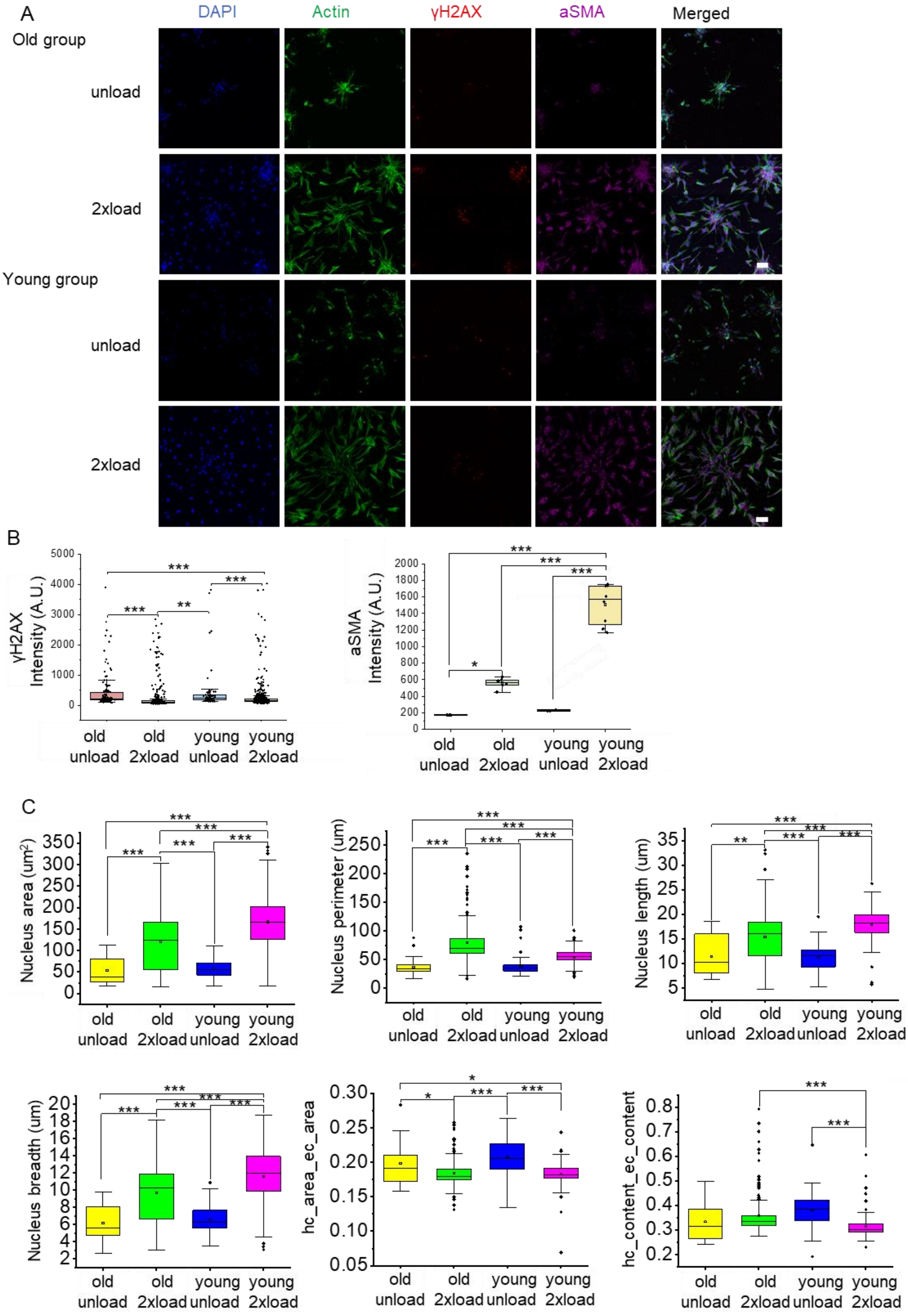

Supplementary figure 3

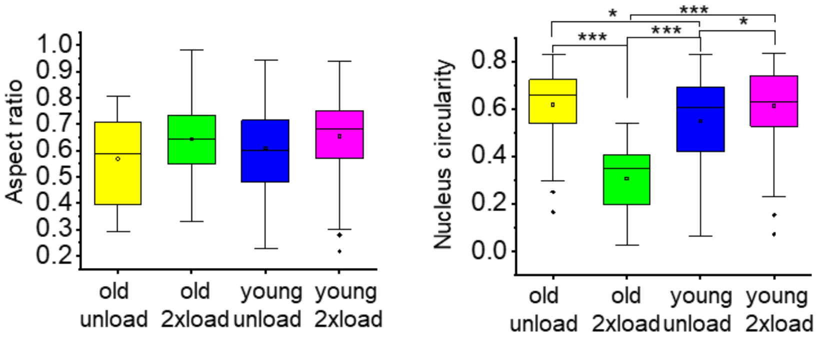


D

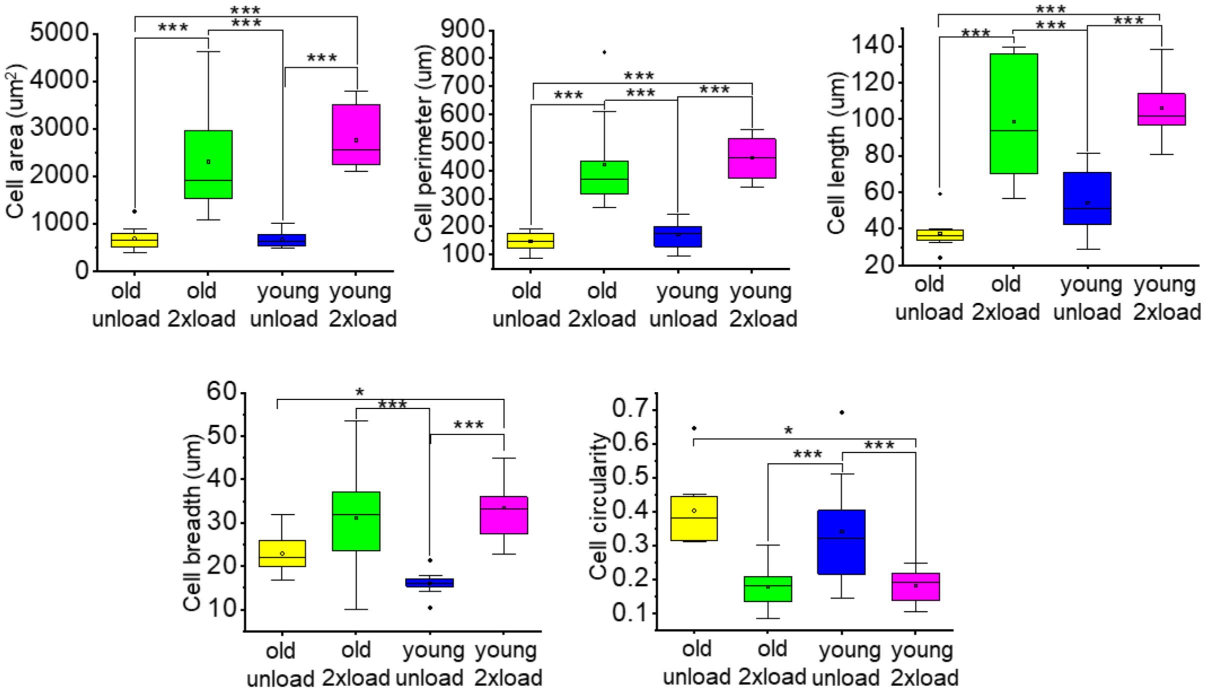

Supplementary Figure.3. Reduced DNA damage, cellular activation, and altered cell and nucleus morphology in old and young groups (metal ring). (A) Representative actin, γH2AX and αSMA immunofluorescence confocal images in young (GM09503 human fibroblasts, 10 years old) and old group (GM08401 human fibroblasts, 75 years old). Nucleus is labeled in blue. (Scale bar, 100*μm*). (B) Quantification data per image of γH2AX and αSMA mean intensity in old and young groups. (C) Quantification data of nucleus morphology (area, perimeter, length, breadth and circularity) and the ratio of heterochromatin/euchromatin (Metal ring). (D) Quantification data of cell morphology (area, perimeter, length, breadth and circularity). (Metal ring). *All the experiments were repeated at least three times independently with similar results. P values in Figure (B-D) were calculated by the one-way ANOVA method with Tukey's post hoc test. *P<0.05; **<0.01; ***P<0.001; No asterisks mean not significant.*

Suppmentary figure 4

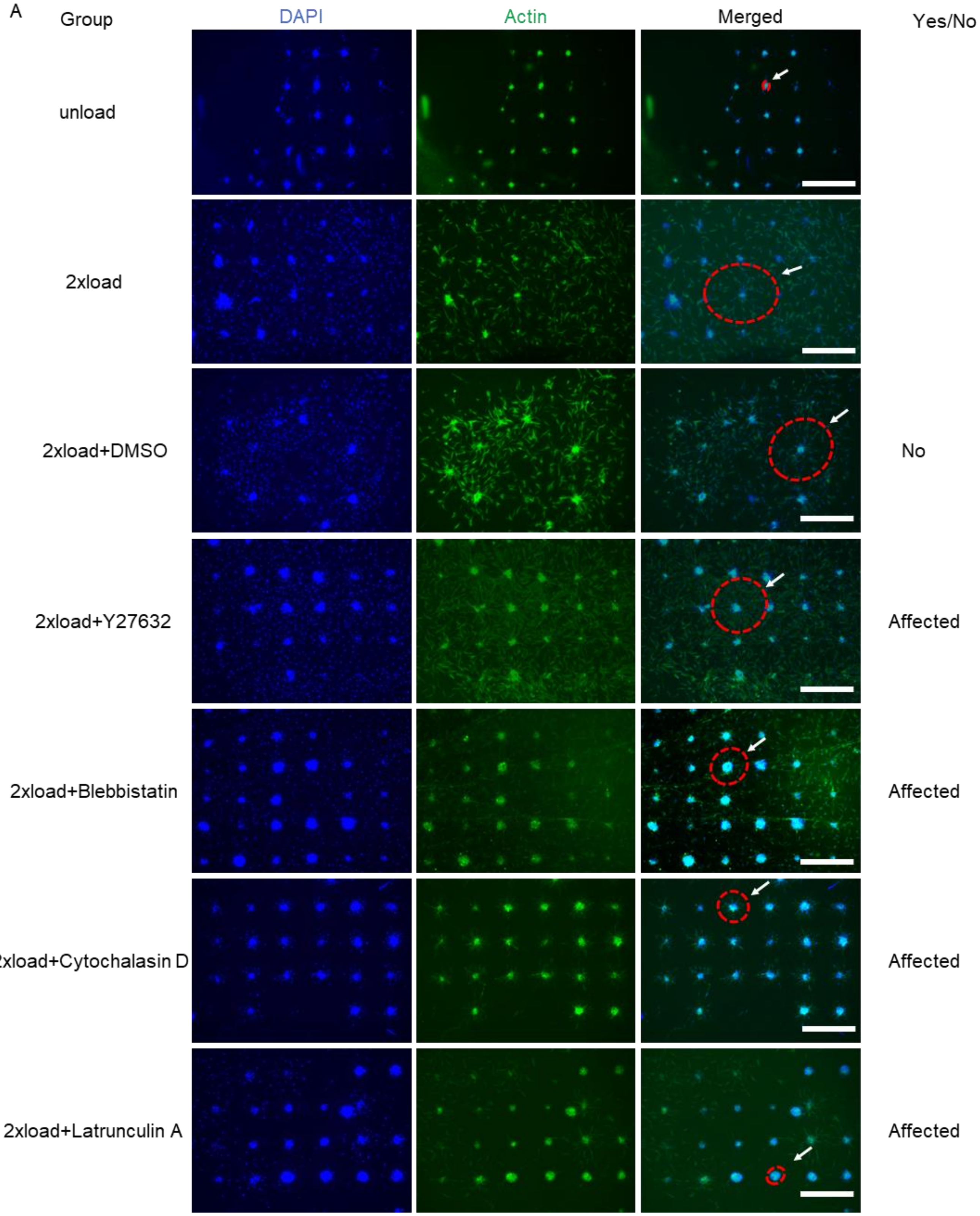

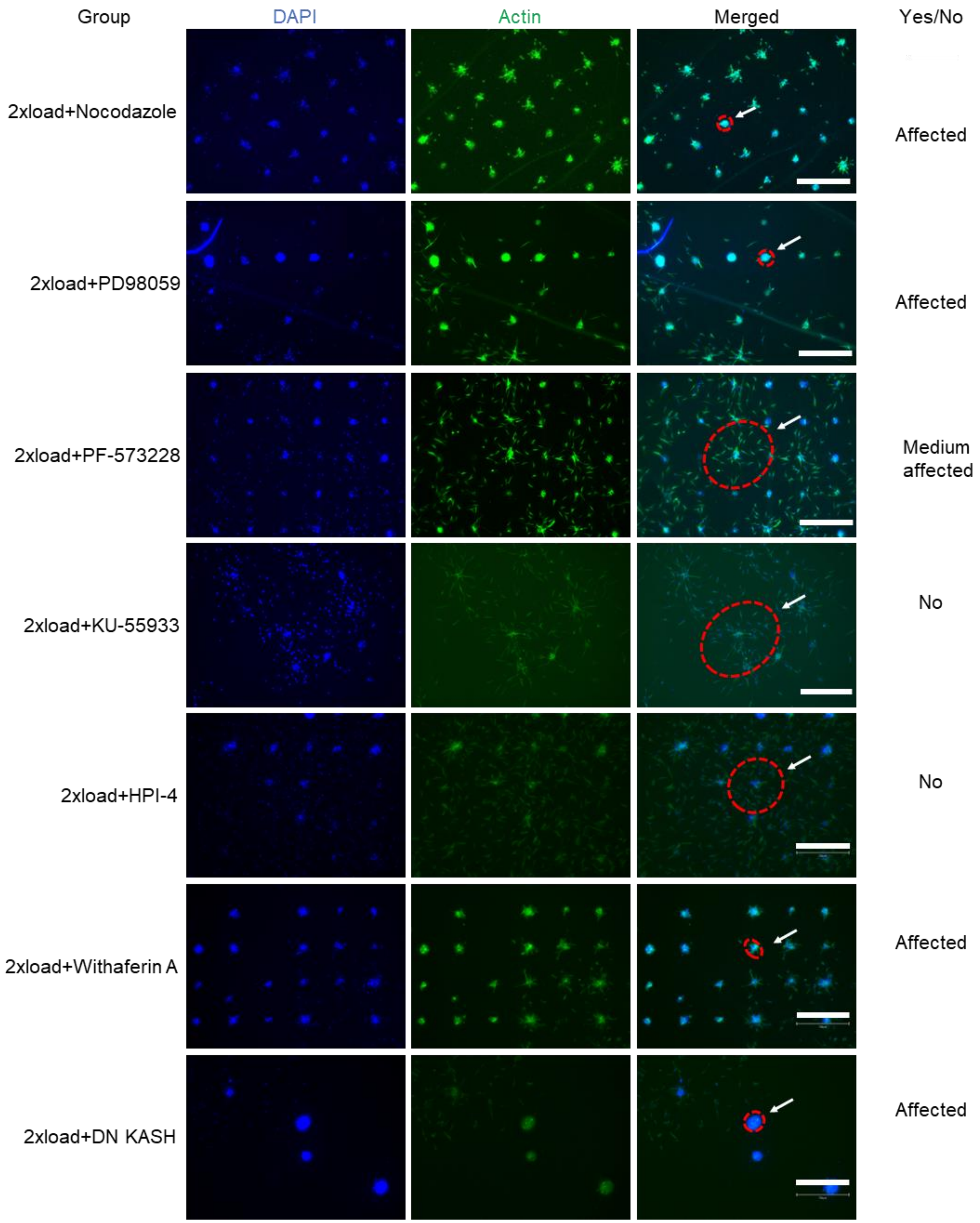

Group
DAPI
Actin
Merged
Yes/No
2xload+Nocodazole
Affected
2xload+PD98059
Affected
2xload+PF-573228
Medium affected
2xload+KU-55933
No
2xload+HPI-4
No
2xload+Withaferin A
Affected
2xload+DN KASH
Affected

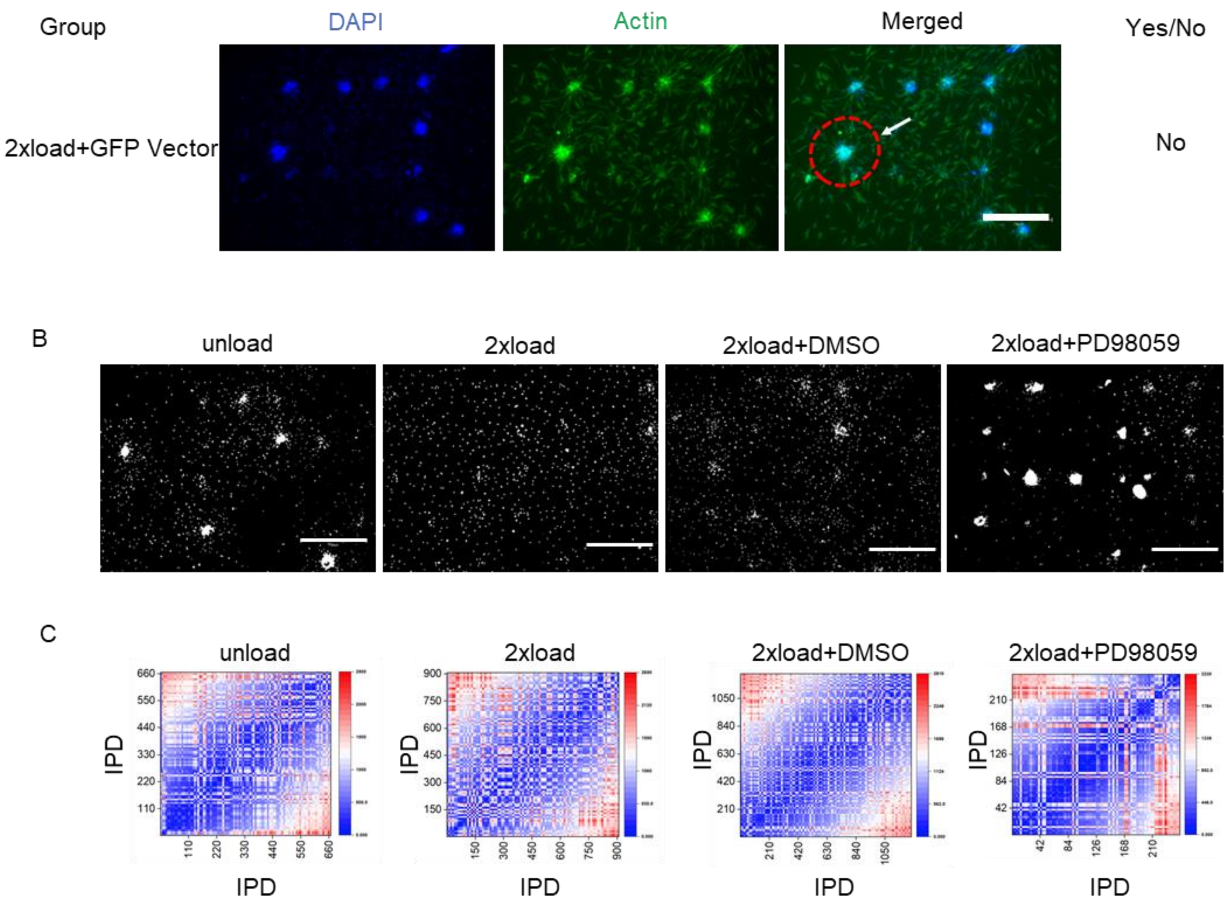

Group
DAPI
Actin
Merged
Yes/No
2xload+GFP Vector
No
B
unload
2xload
2xload+DMSO
2xload+PD98059
C
unload
2xload
2xload+DMSO
2xload+PD98059
IPD
IPD
IPD
IPD
IPD
IPD
IPD
IPD

Supplementary Figure.4. Pharmacological perturbations to identify critical signaling intermediate in the old group via metal ring or glass ring. (A) DMSO as control group; Y27632 is ROCK inhibitor; Blebbistatin is non-muscle myosin II inhibitor; Cytochalasin D disrupts actin microfilaments; Latrunculin A inhibits actin polymerization; Nocodazole is microtubules inhibitor; PD98059 is MEK1 and MEK2 inhibitor; PF-573228 is FAK inhibitor; KU-55933 is ATM inhibitor; Above inhibitor experiments were carried out using the metal ring and inhibitor experiments below was performed by glass ring. HPI-4 is dynein inhibitor; Withaferin A is vimentin inhibitor; DN KASH is Nesprin 4 inhibitor. (Scale bar, 750$\mu m$). Red circle labels spheroid spreading. Green color (actin), Blue color (DAPI). (B) Representative gray images for doubly checking interesting inhibitors effect by glass ring. (Scale bar, 750$\mu m$). (C) Internuclear pairwise distance (IPD) analysis of images from B.

Suppmentary figure 5

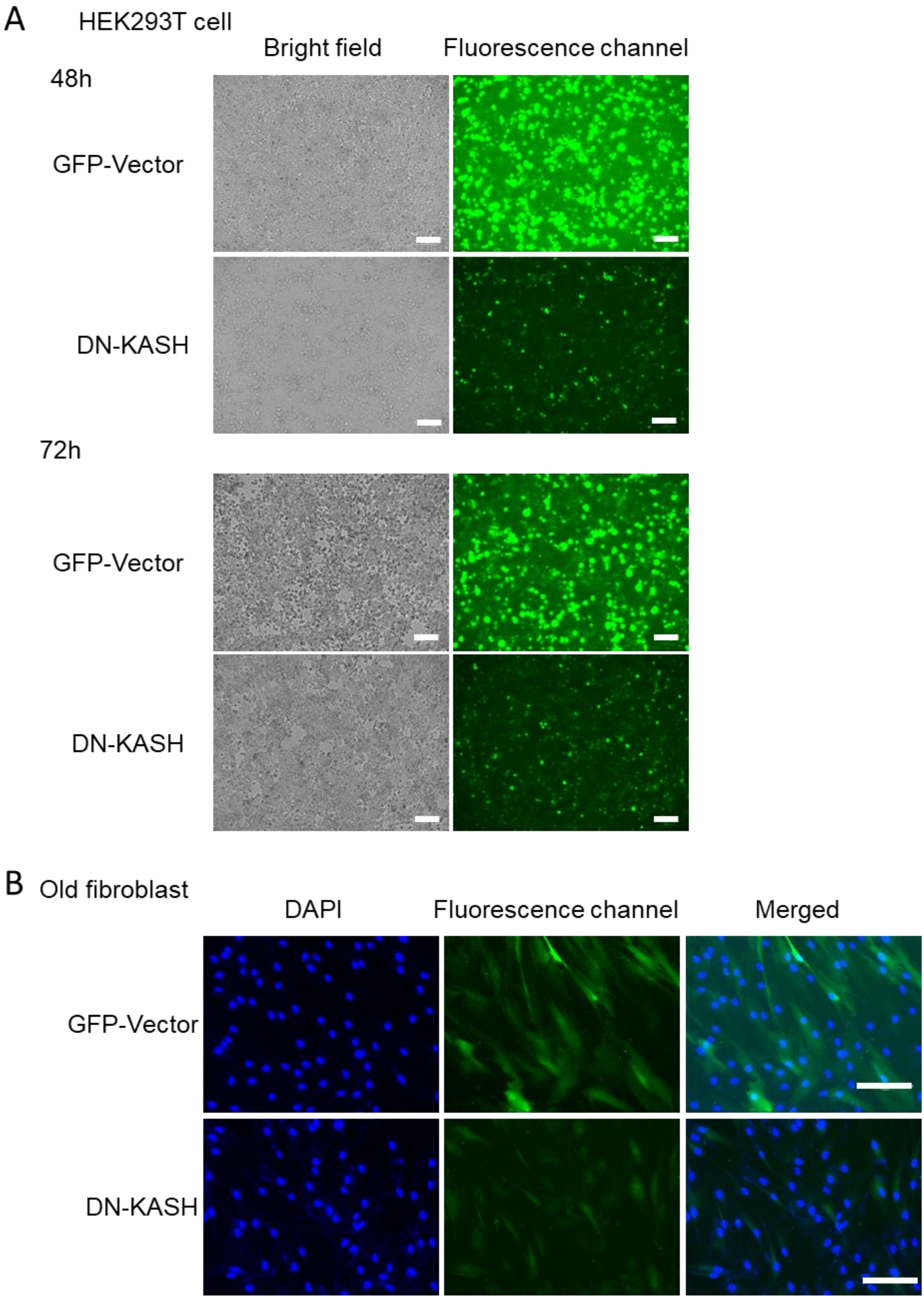

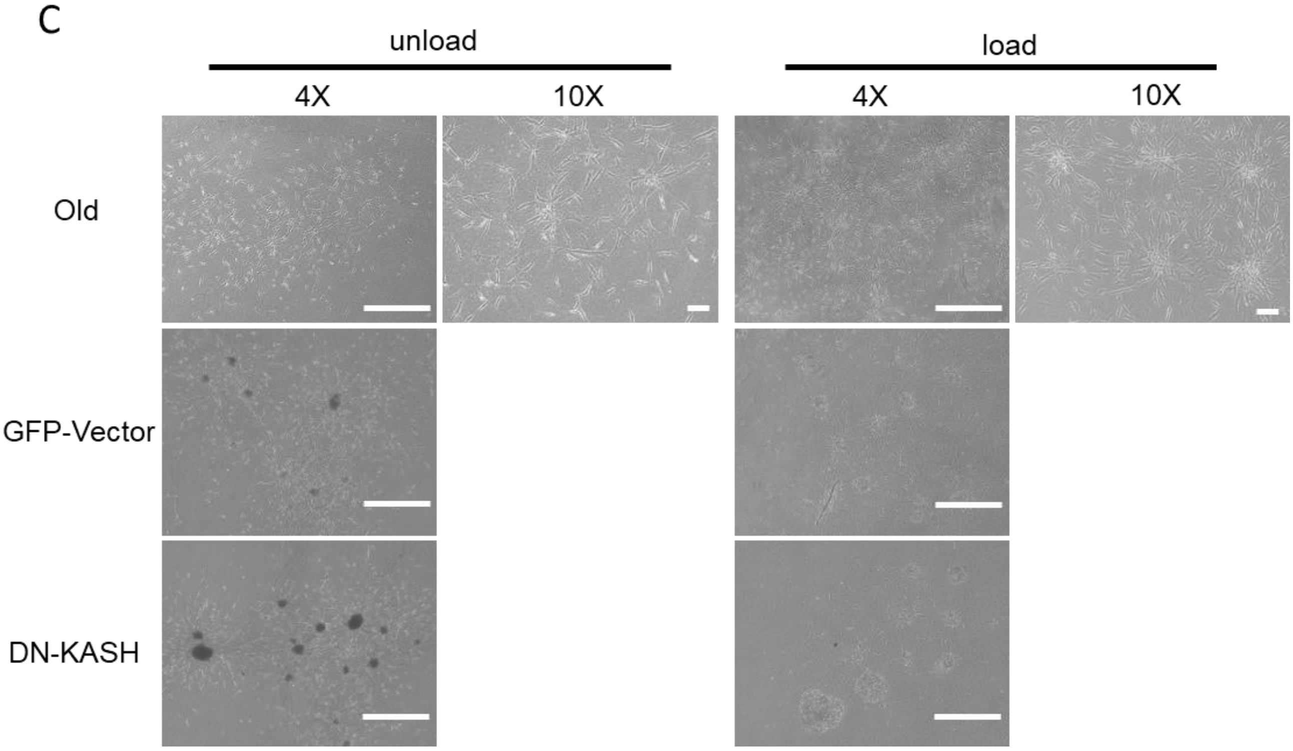
C
unload
load
4X
10X
4X
10X
Old
GFP-Vector
DN-KASH

Supplementary Figure .5. Lentivirus transfection efficiency. (A) Representative bright field images and green fluorescence images. The HEK293 cell lines were used to produce lentivirus. (Scale bar, 100*μm*). (B) Purified lentivirus was used to transfect old fibroblast. (Scale bar, 150*μm*). (C) Representative bright field images of fibroblast in a 3D model under compressive conditions. GFP-Vector alone is transfected with lentivirus vectors and the other group were transfected by lentivirus with DN-KASH-GFP construct. (Scale bar, 4X objective is 750*μm*, 10X objective is 100*μm*).

Suppmentary figure 6

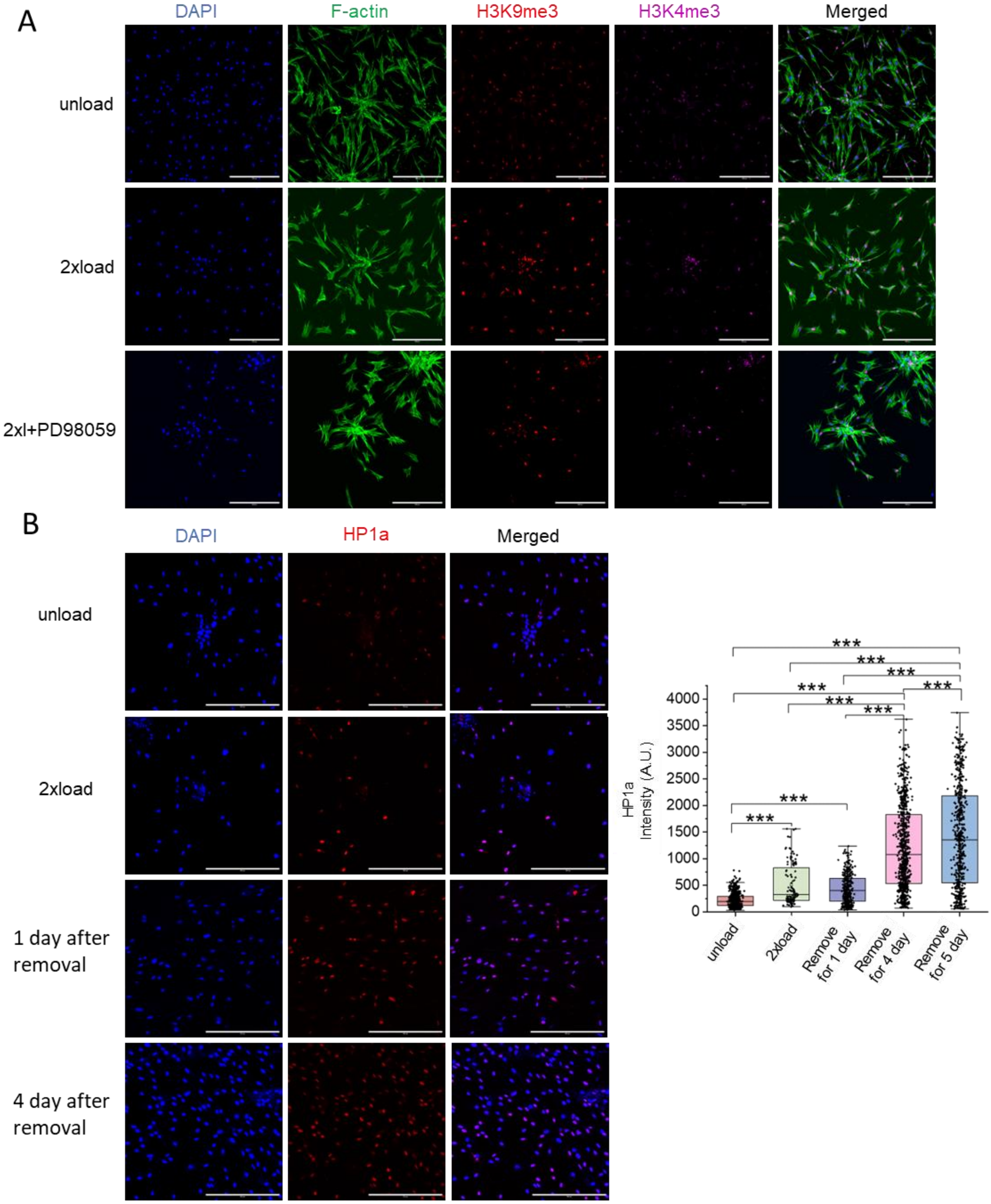

Suppmentary figure 6

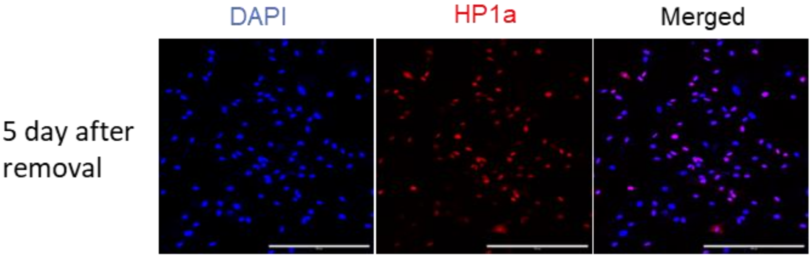


C

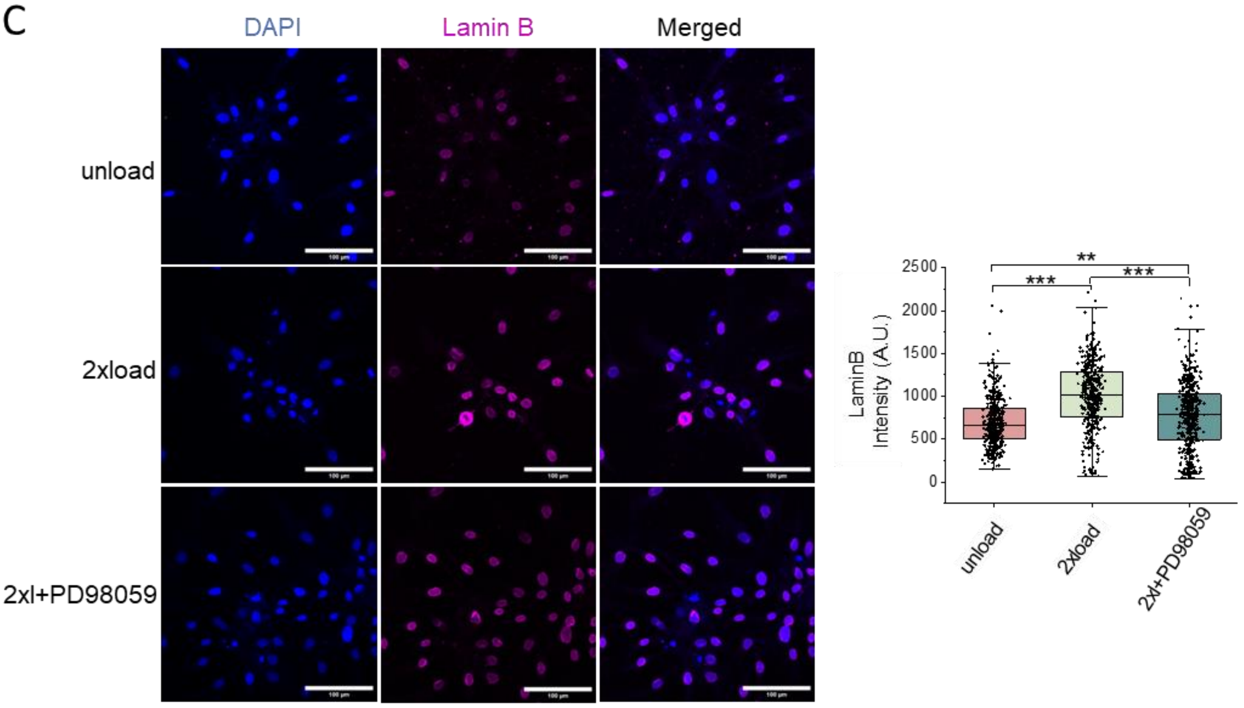


D

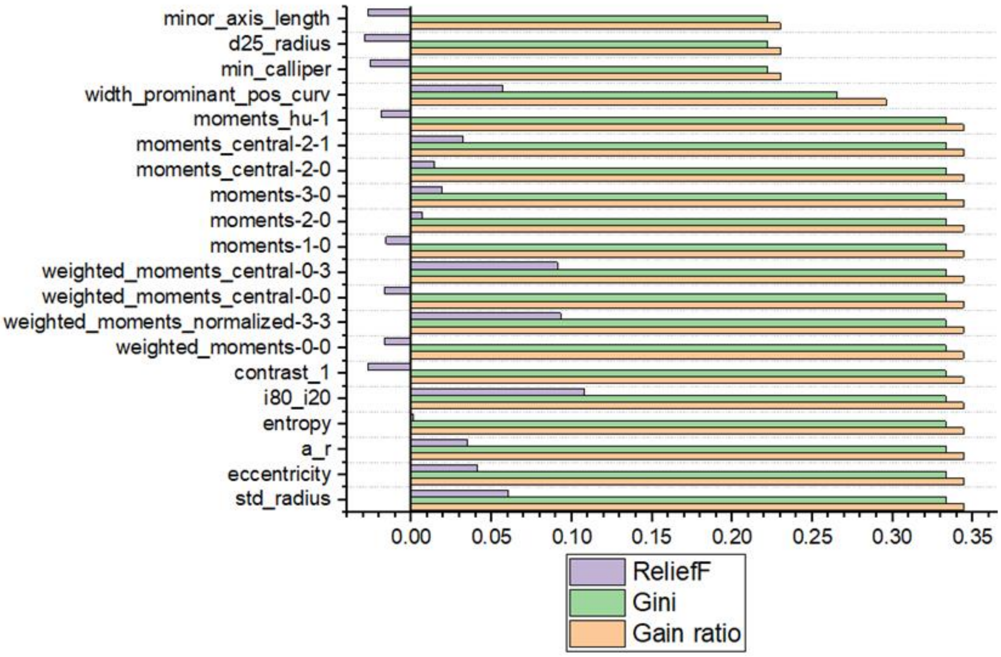

Supplementary Figure.6. Individual channels of immunofluorescence staining images in Figure 2 and Figure 4. (A) Representative H3K9me3 and H3K4me3 immunofluorescence confocal images in the unload group, 2xload group and 2xl+PD98059 group. (Scale bar, 300*μm*). (B) Representative HP1a immunofluorescence confocal images in unload group, 2xload group, 1 day after removal group, 4 day after removal group and 5 day after removal group. (Scale bar, 300*μm*). Quantification data per nucleus of HP1a mean intensity. *P<0.05; **<0.01; ***P<0.001; (C) Representative Lamin B immunofluorescence confocal images in unload group, 2xload group and 2xl+PD98059 group. (Scale bar, 100*μm*). Quantification data per nucleus of Lamin B mean intensity. (D) Importance analysis of each attribute Figure 2D. *Outliers are symbolized using asterisks. All the experiments were repeated at least three times independently with similar results. P values in Figure (B and C) were calculated by the one-way ANOVA method with Tukey's post hoc test. *P<0.05; **<0.01; ***P<0.001; No asterisks mean not significant.*

Suppmentary figure 7

A

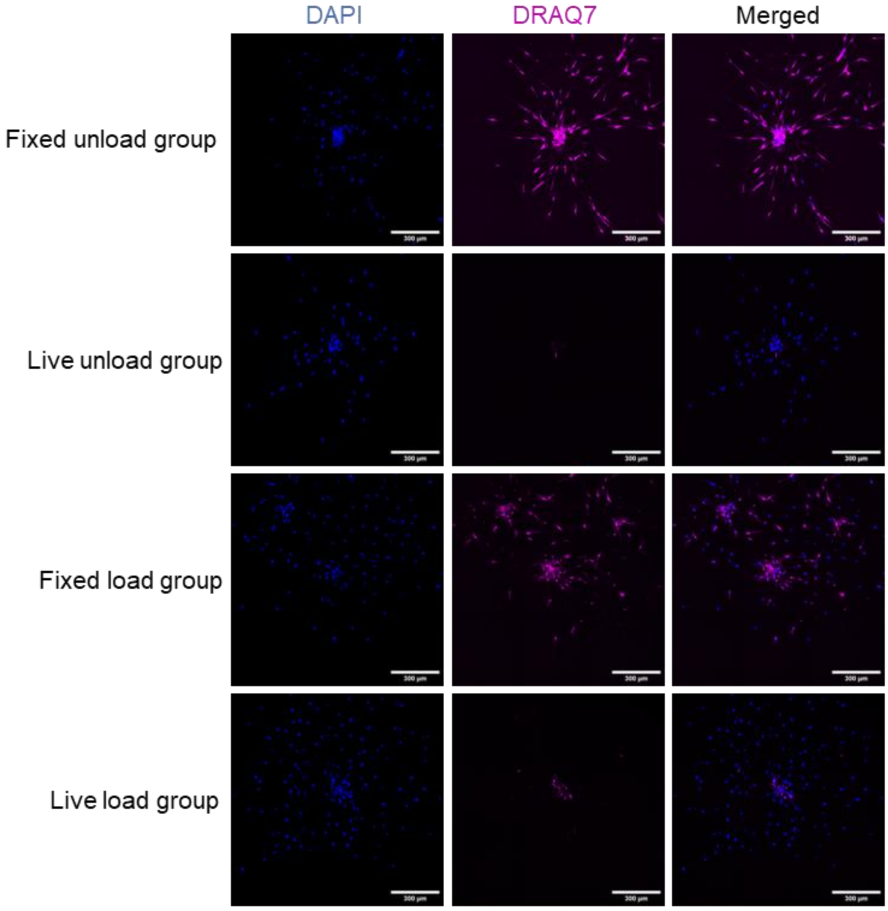


B

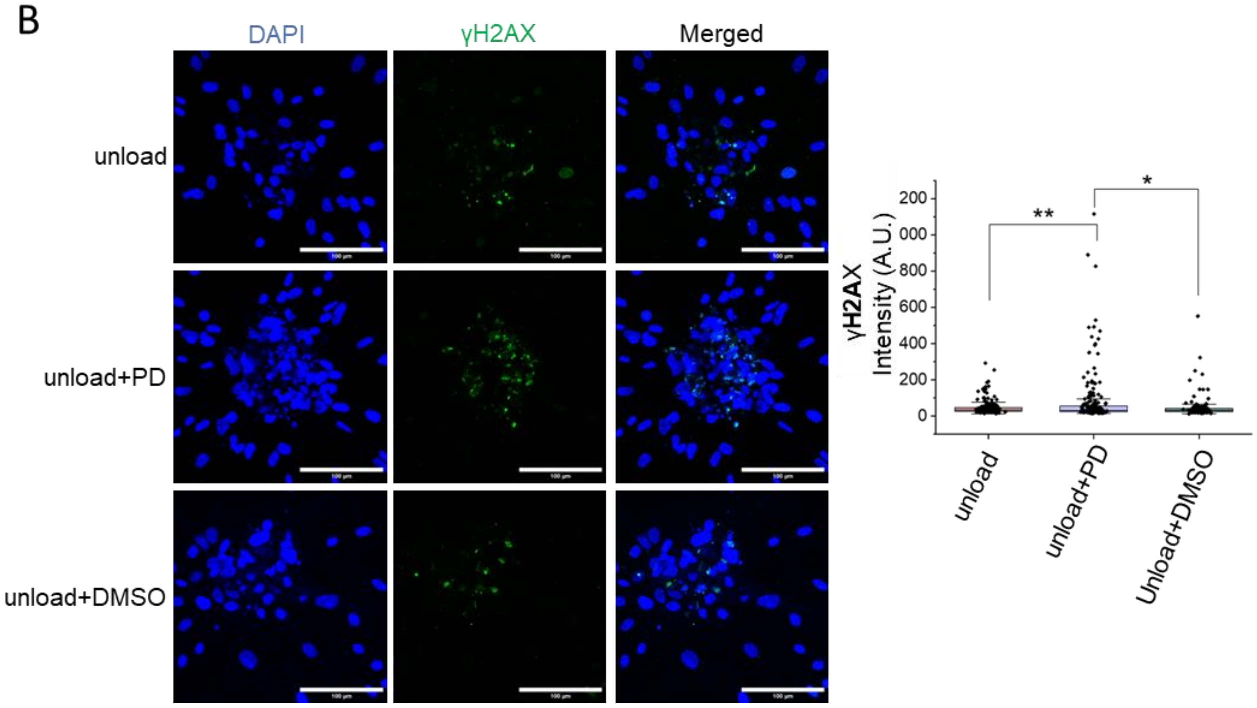

Suppmentary figure 7

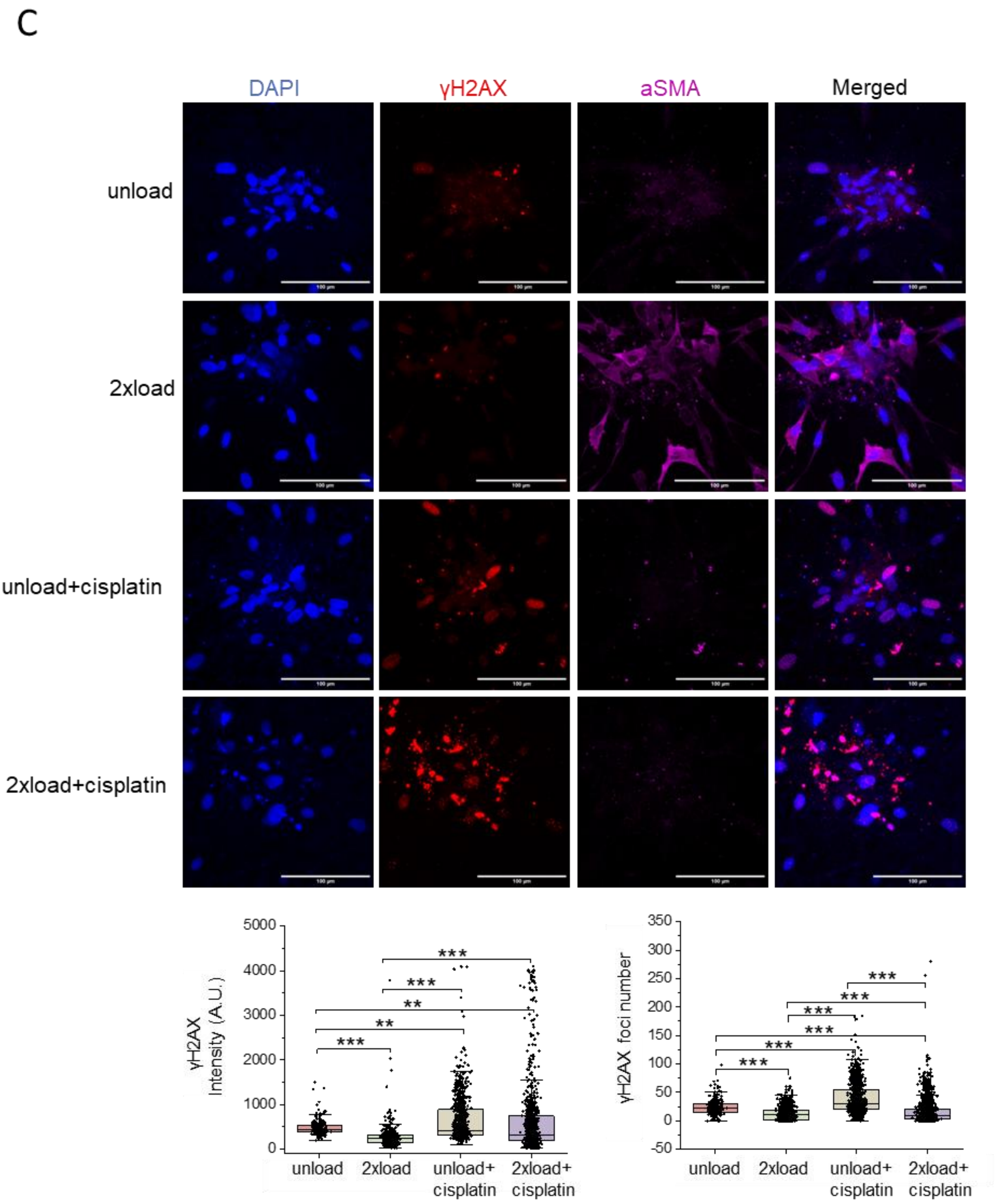

Supplementary Figure.7. Cell viability and γH2AX expression in different conditions. (A) DRAQ7 Dye staining for dead cells in different conditions. Fixed unload group and fixed load group are regarded as control group. (Scale bar, 300*μm*). (B) Representative γH2AX immunofluorescence confocal images in unload group, unload+PD98059 group and unload+DMSO group. Quantification data per nucleus of γH2AX mean intensity. (Scale bar, 100*μm*). (C) Individual channels of γH2AX, αSMA immunofluorescence staining images in Figure 4E. Quantification data per nucleus of γH2AX mean intensity and γH2AX foci number. (Scale bar, 100*μm*). *All the experiments were repeated at least three times independently with similar results. P values in Figure (B and C) were calculated by the one-way ANOVA method with Tukey's post hoc test. *P<0.05; **<0.01; ***P<0.001; No asterisks mean not significant.*

Suppmentary figure 8

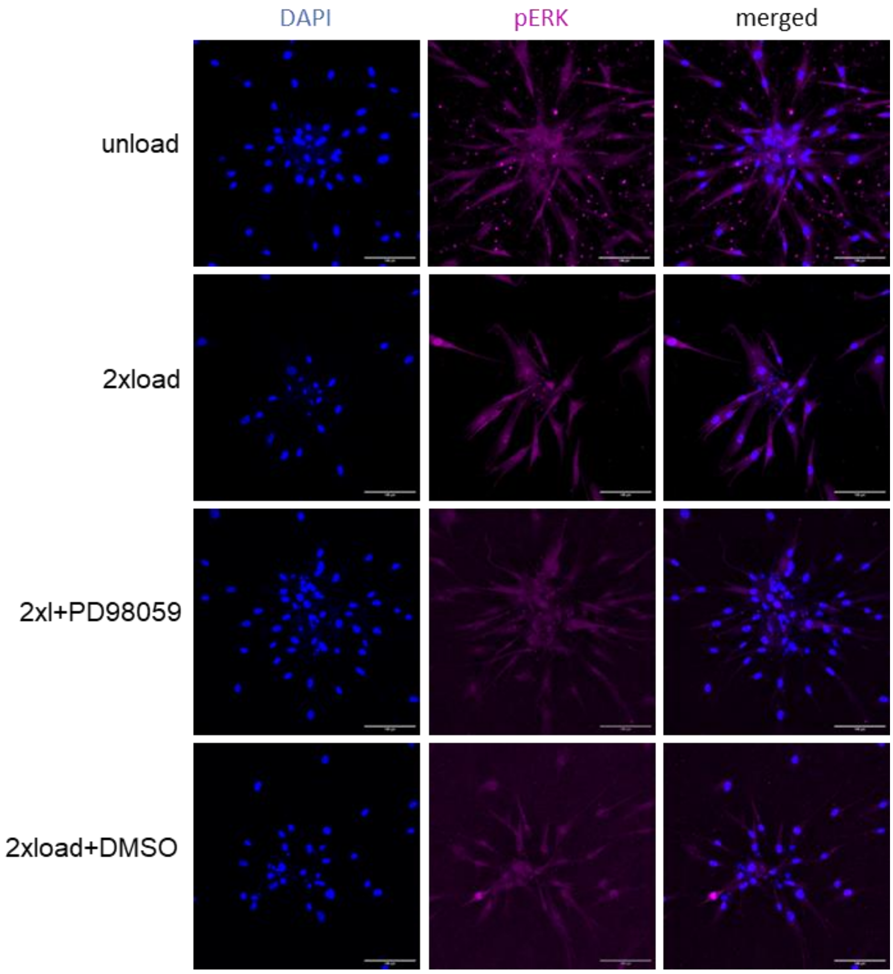

Supplementary Figure.8. Representative pERK immunofluorescence confocal images in different groups. (Scale bar, 100*μm*).

Suppmentary figure 9

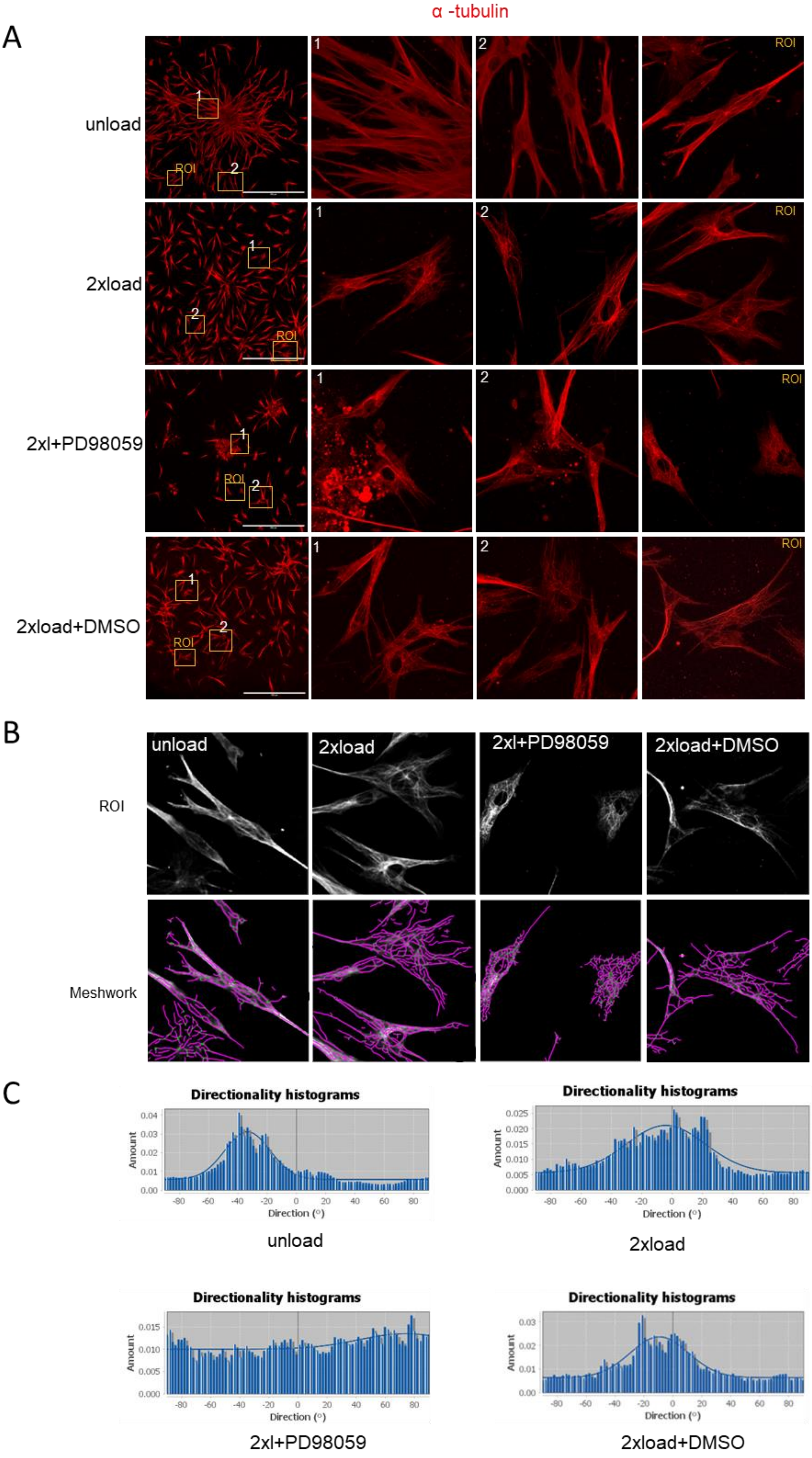

Supplementary Figure.9. Microtubule network analysis in different groups. (A) Representative *α*-tubulin immunofluorescence confocal images and selected regions of interest. (Scale bar, 500*μm*). (B) ROI meshwork construct. (C) directionality histograms for ROI meshwork.

Suppmentary figure 10

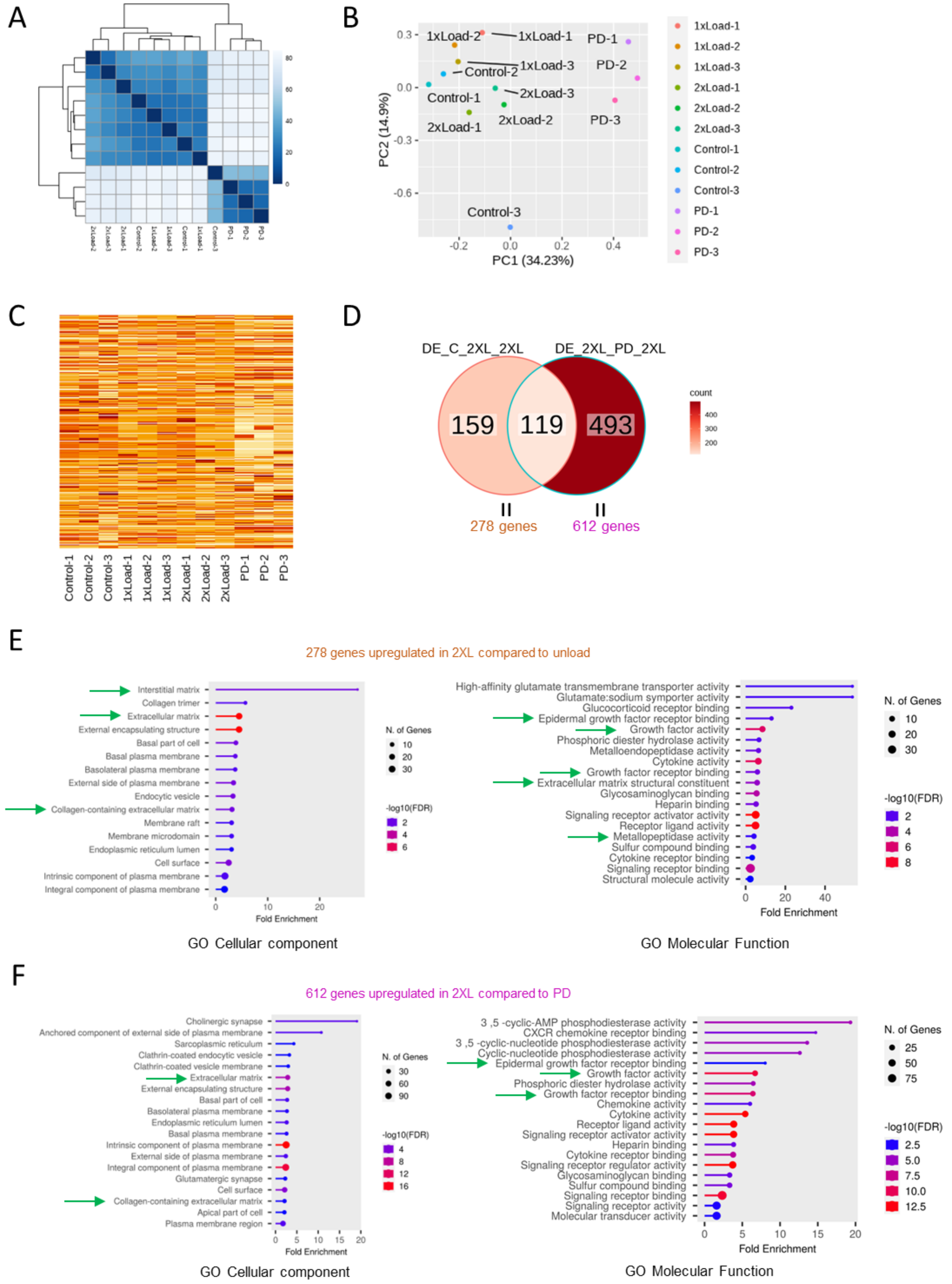

Suppmentary figure 10

G

119 shared genes upregulated in 2XL compared to unload and PD

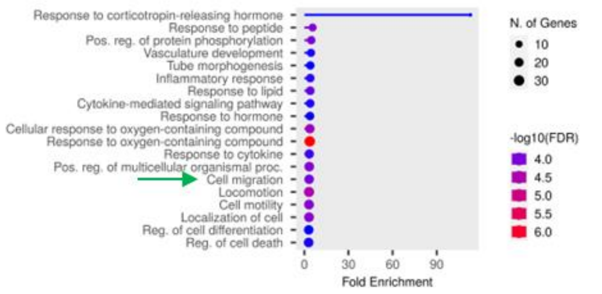


GO Biological process

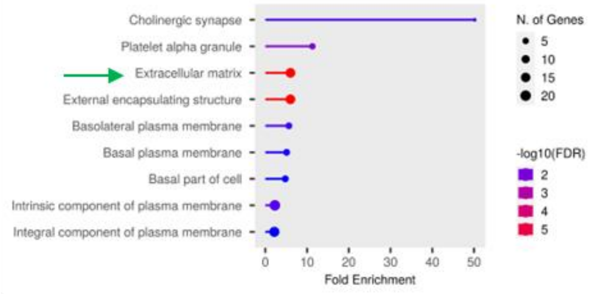


GO Cellular component

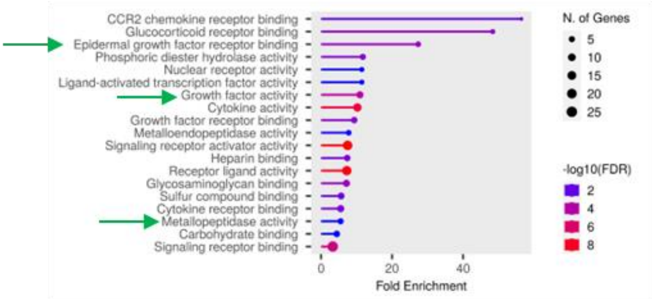


GO Molecular Function

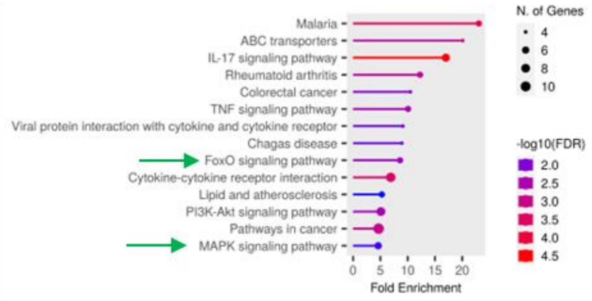


KEGG

Supplementary Figure.10. RNAseq data analysis. (A) Hierarchical clustering analysis for different groups. Control-1, control-2 and control-3 belong to the unload group (three biology replicates). 1xLoad-1, 1xLoad-2 and 1xLoad-3 belong to the 1xload group (three biology replicates). 2xLoad-1, 2xLoad-2 and 2xLoad-3 belong to the 2xload group (three biology replicates). PD-1, PD-2 and PD-3 belong to the 2xload+PD98059 group (three biology replicates). (B) Principal Component Analysis of the gene expression profile in four groups. (C) The heat maps show gene expression profiles of all the four groups. (D) Venn diagram showing the gene overlap between different groups. 278 genes upregulated in the 2xload group compared to the unload group. 612 genes upregulated in the 2xload group compared to the PD group. 119 genes overlap between 278 DEG and 612 DEG. (E) GO Cellular component plot and GO Molecular Function plot for 278 DEG list. (F) GO Cellular component plot and GO Molecular Function plot for 612 DEG list. (G) GO Biological process plot, GO Cellular component plot, GO Molecular Function plot and KEGG plot for 119 DEG list.

Suppmentary figure 11

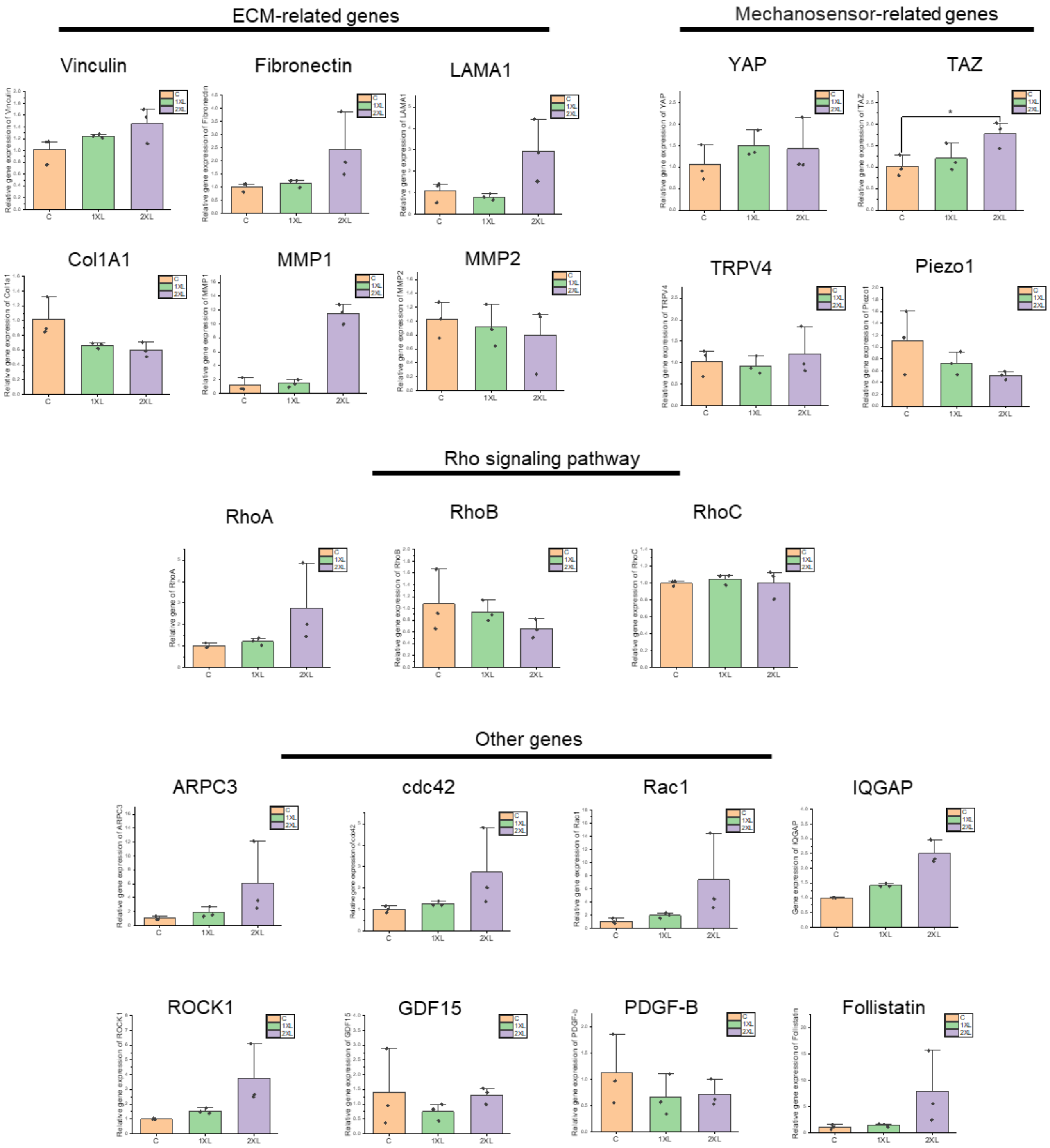

Supplementary Figure.11. Validation of gene expression related to cellular migration using qRT-PCR method. *Outliers are symbolized using asterisks. All the experiments were repeated at least three times independently with similar results. P values in Figure were calculated by the one-way ANOVA method with Tukey's post hoc test. *P<0.05; **<0.01; ***P<0.001; No asterisks mean not significant. Source data are provided as a Source Data file.*

Suppmentary figure 12

A

MAPK pathway (278 genes upregulated in 2XL compared to unload )

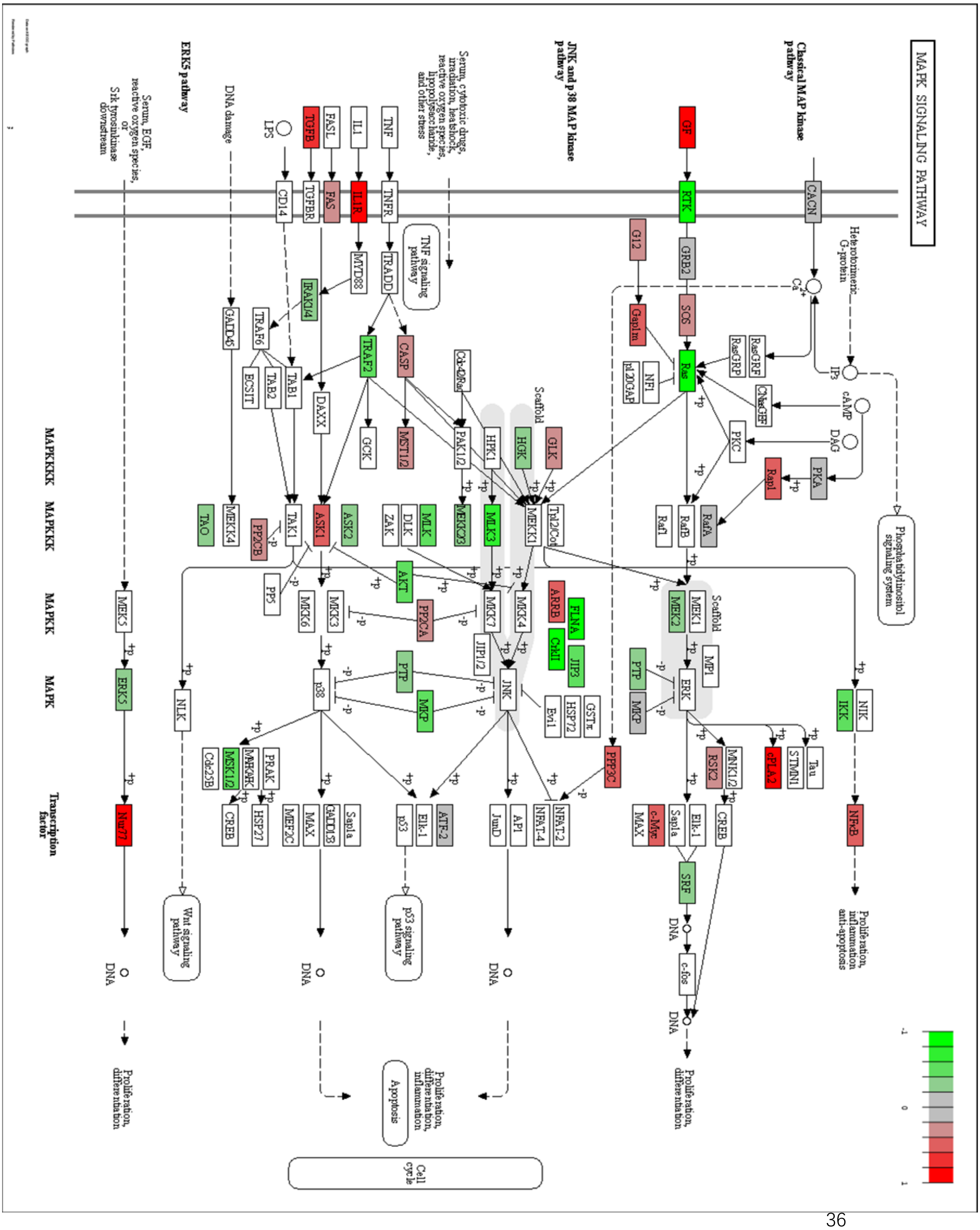

B

MAPK pathway ( 612 genes upregulated in 2XL compared to PD)

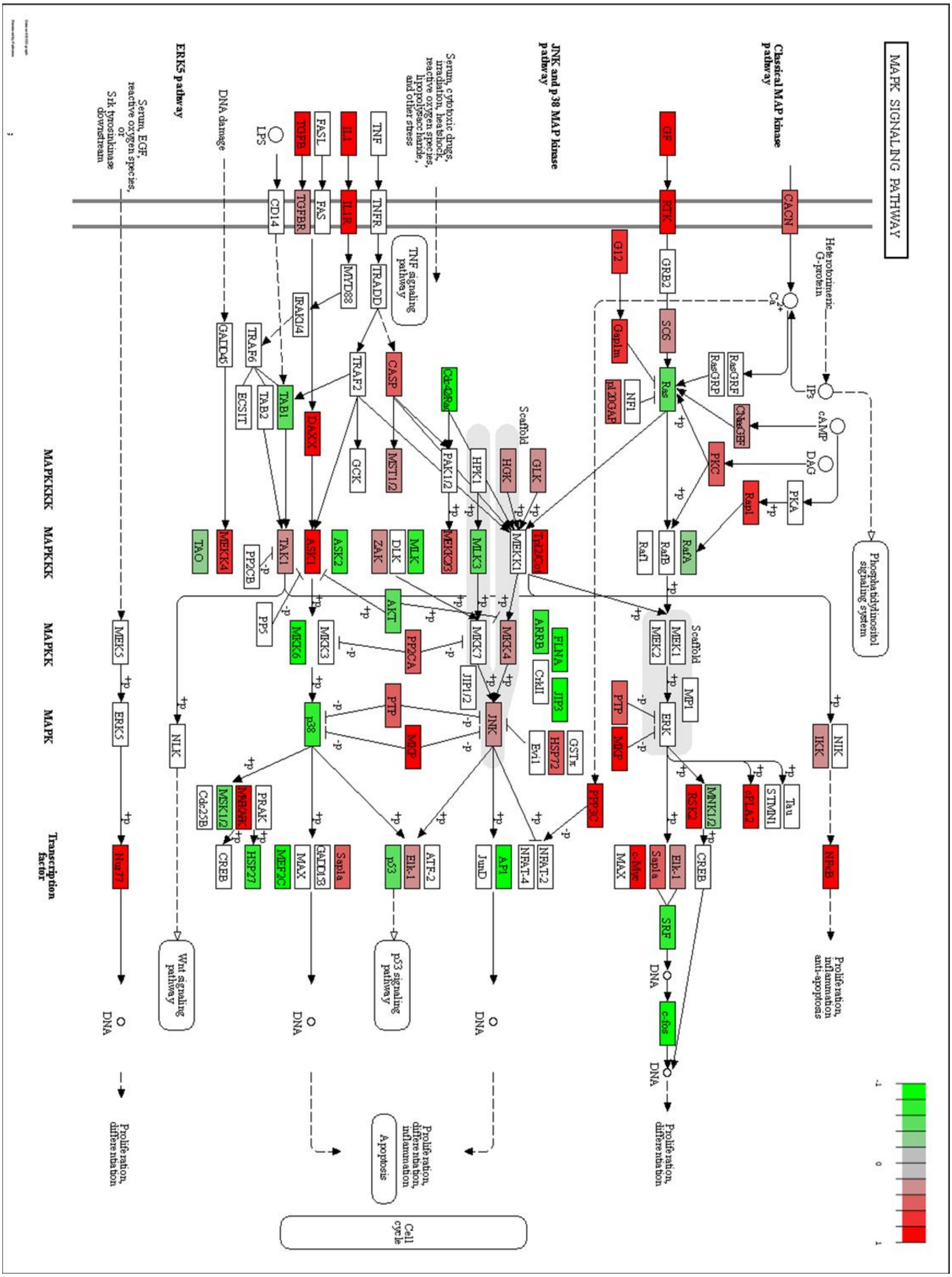

Supplementary Figure.12. MAPK KEGG pathway plot (A) MAPK KEGG pathway plot for 278 DEG list. (B) MAPK KEGG pathway plot for 612 DEG list.

Suppmentary figure 13

DNA repair related pathway (278 genes upregulated in 2XL compared to unload)

A Non-homologous end-joining - Homo sapiens (human)

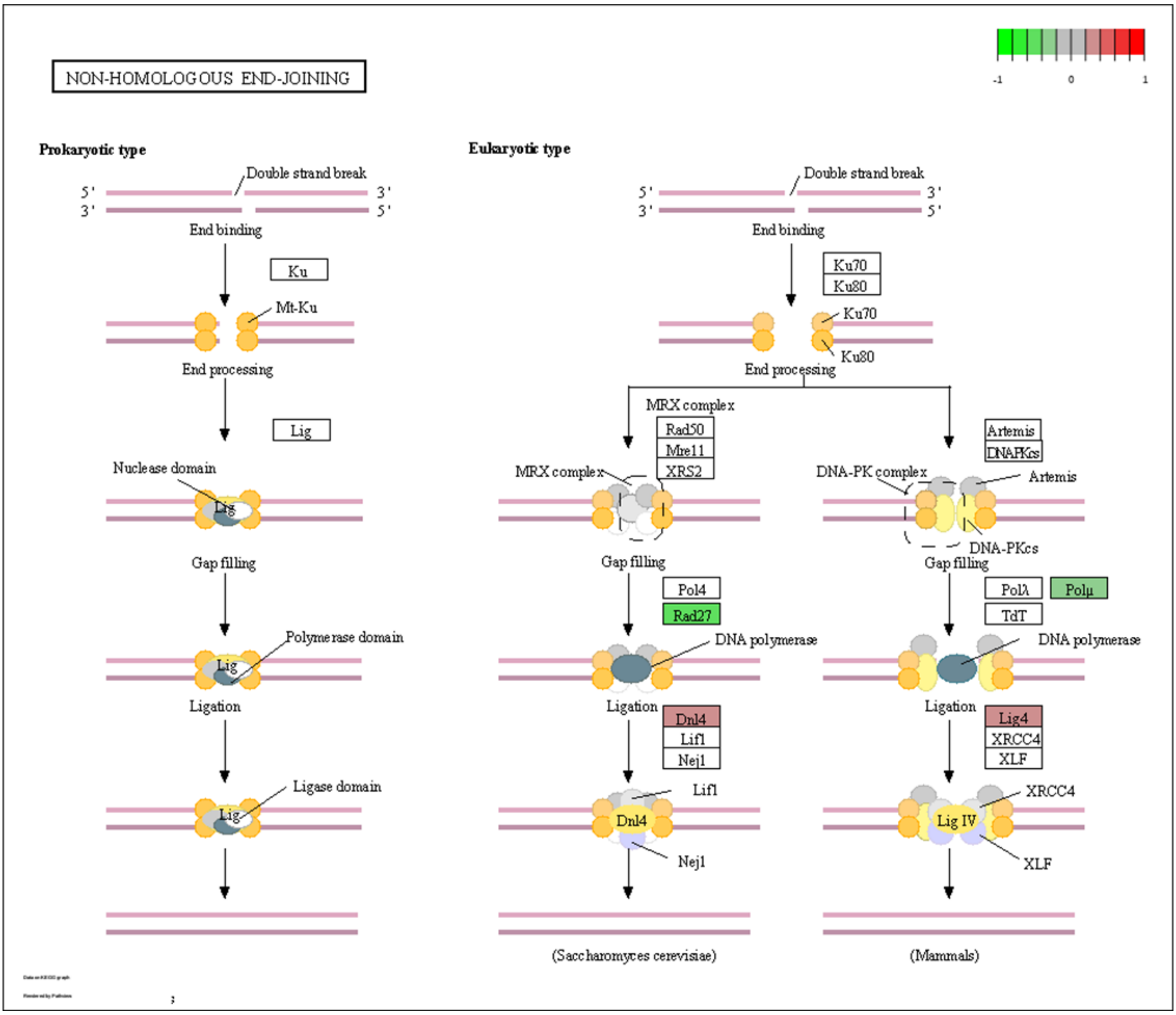

B

Fanconi anemia pathway - Homo sapiens (human)

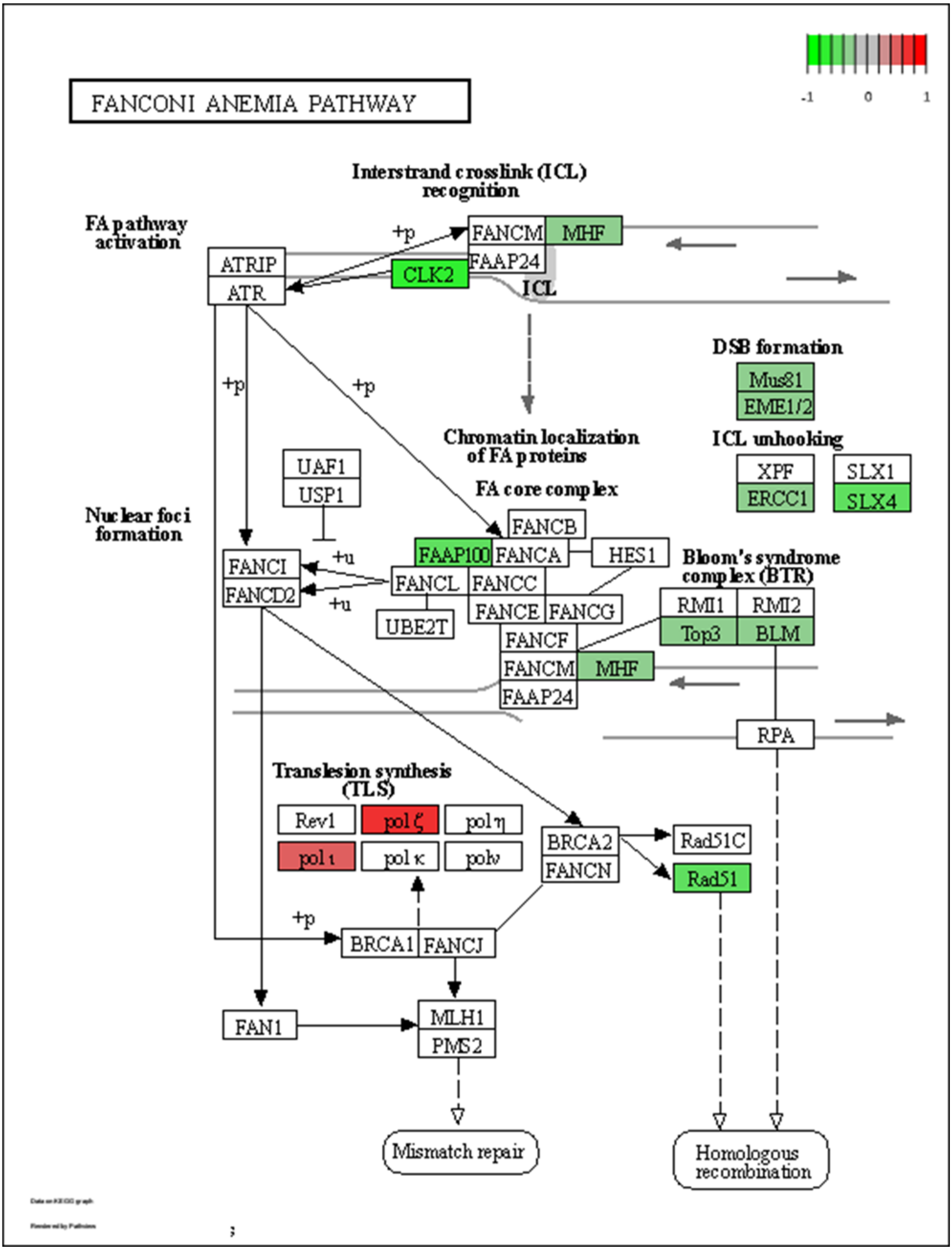

Suppmentary figure 13

C Mismatch repair - Homo sapiens (human)

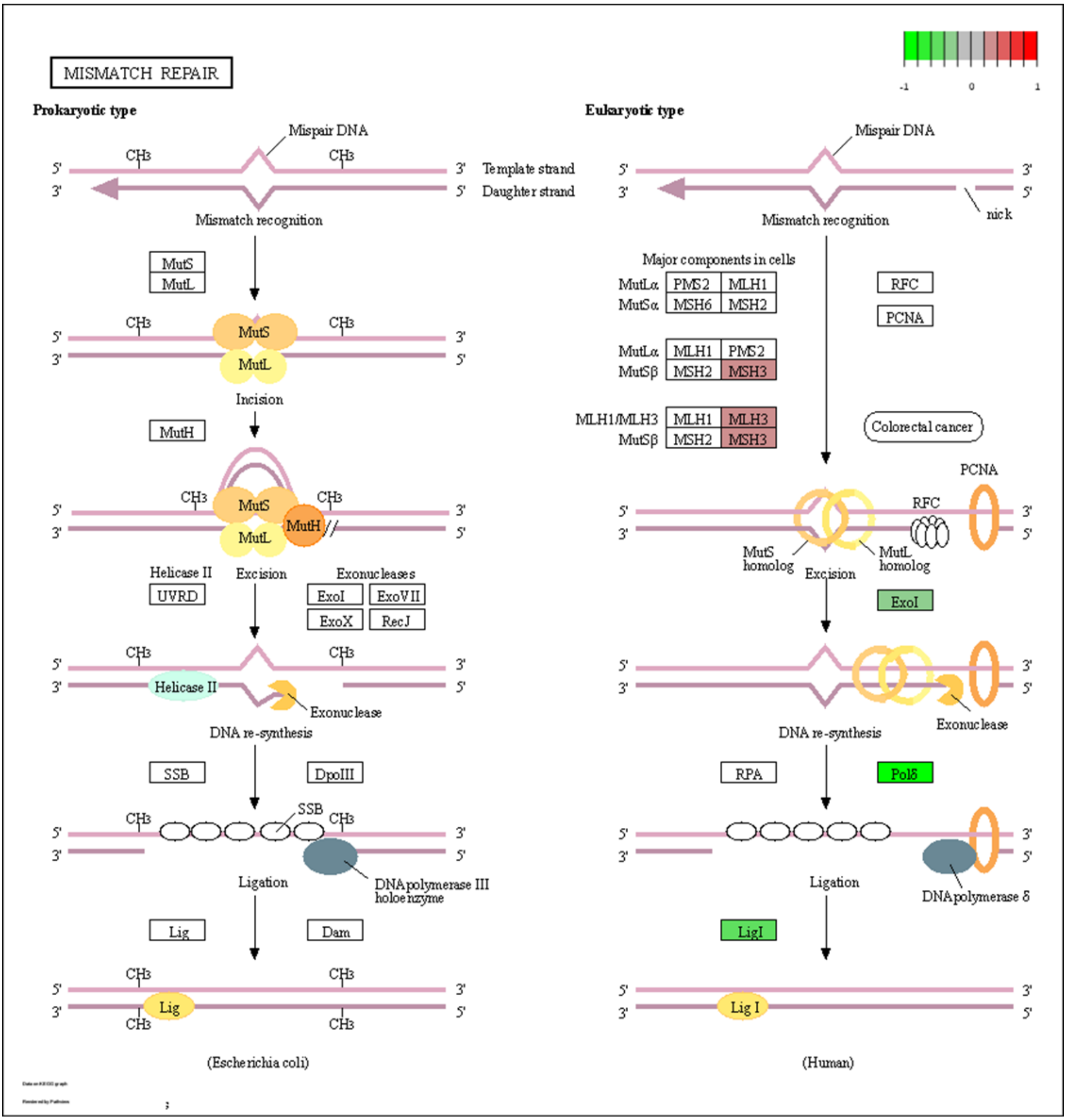

D

Homologous recombination - Homo sapiens (human)

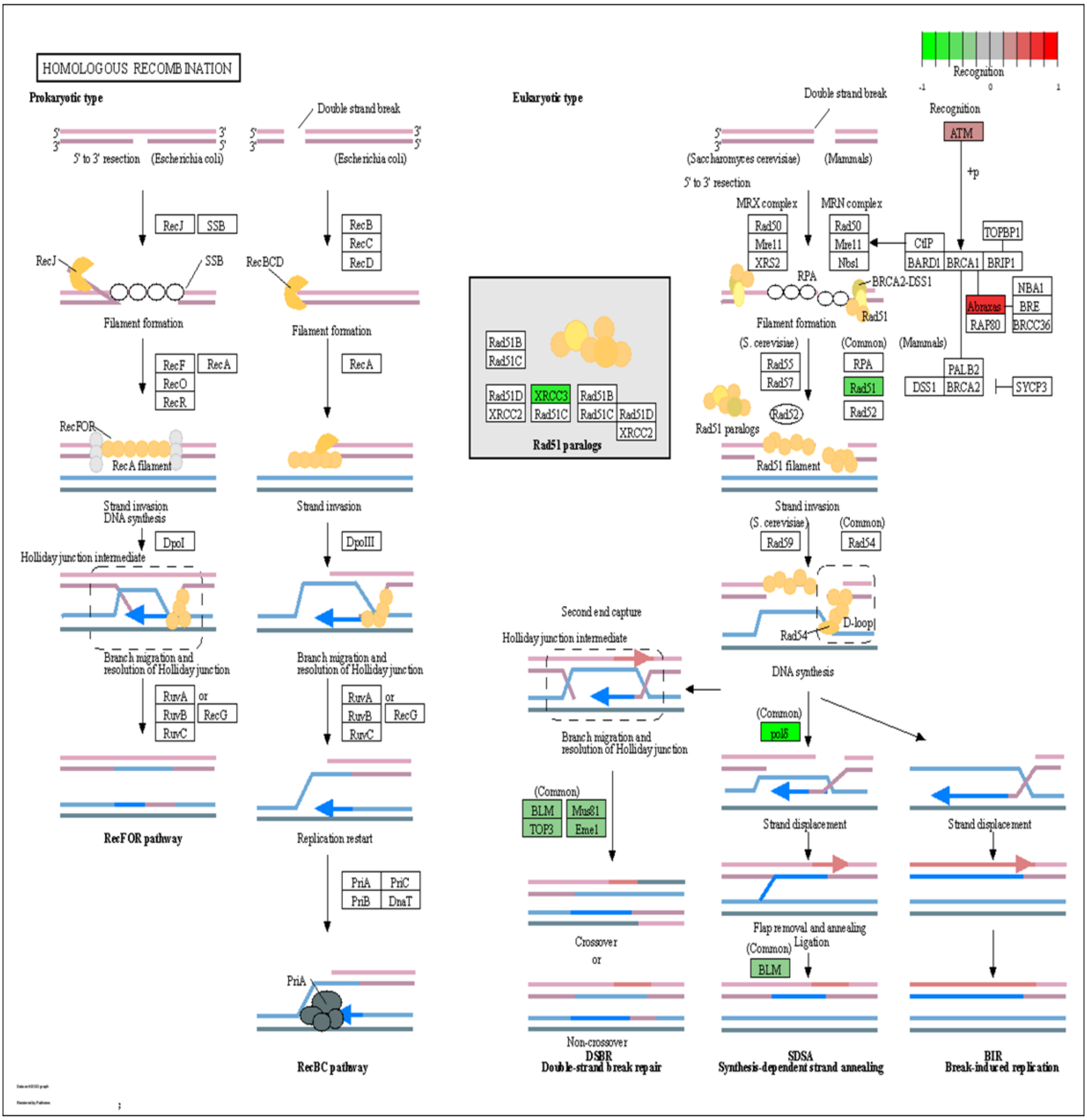

Suppmentary figure 13

E Nucleotide excision repair - Homo sapiens (human)

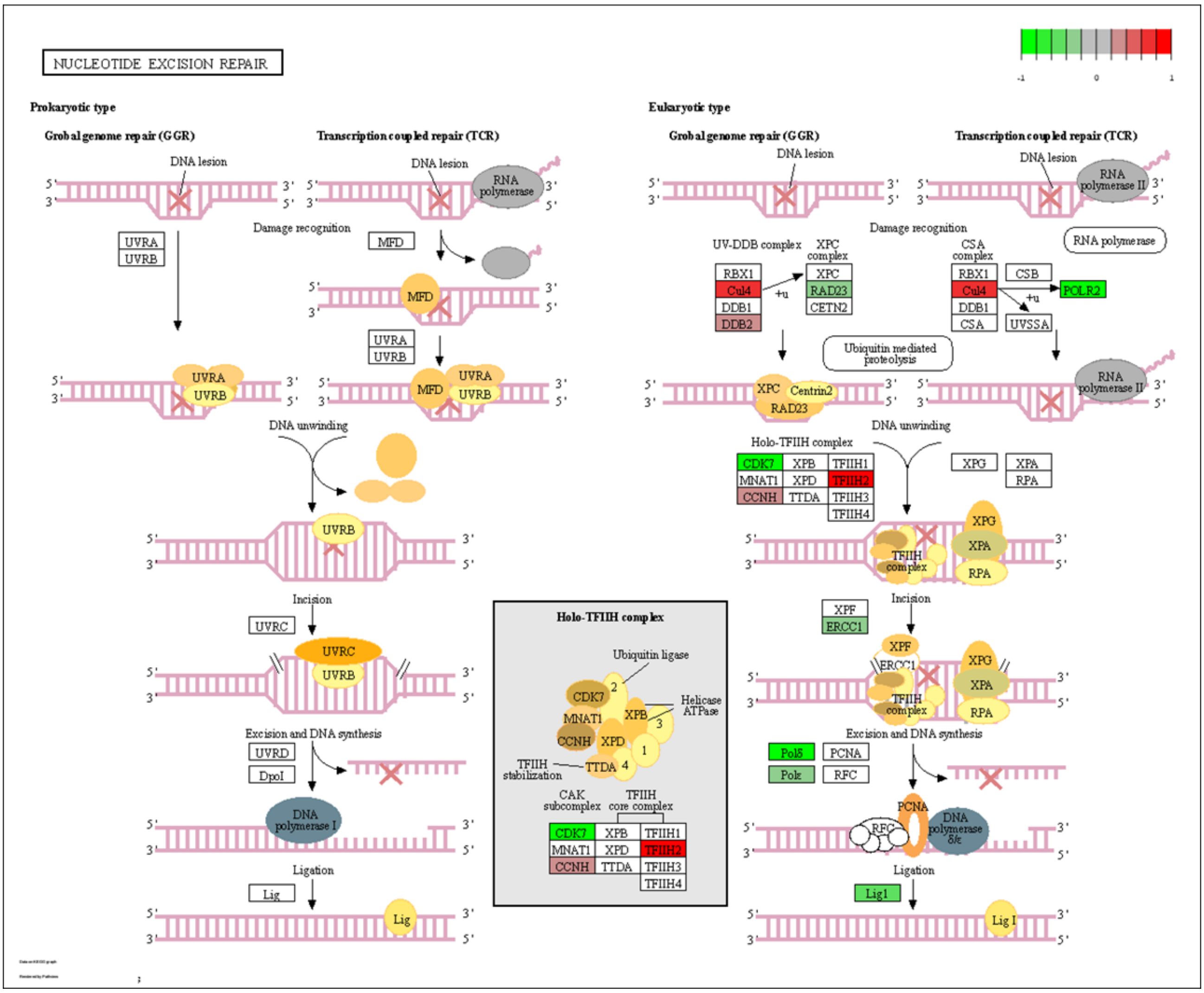

F

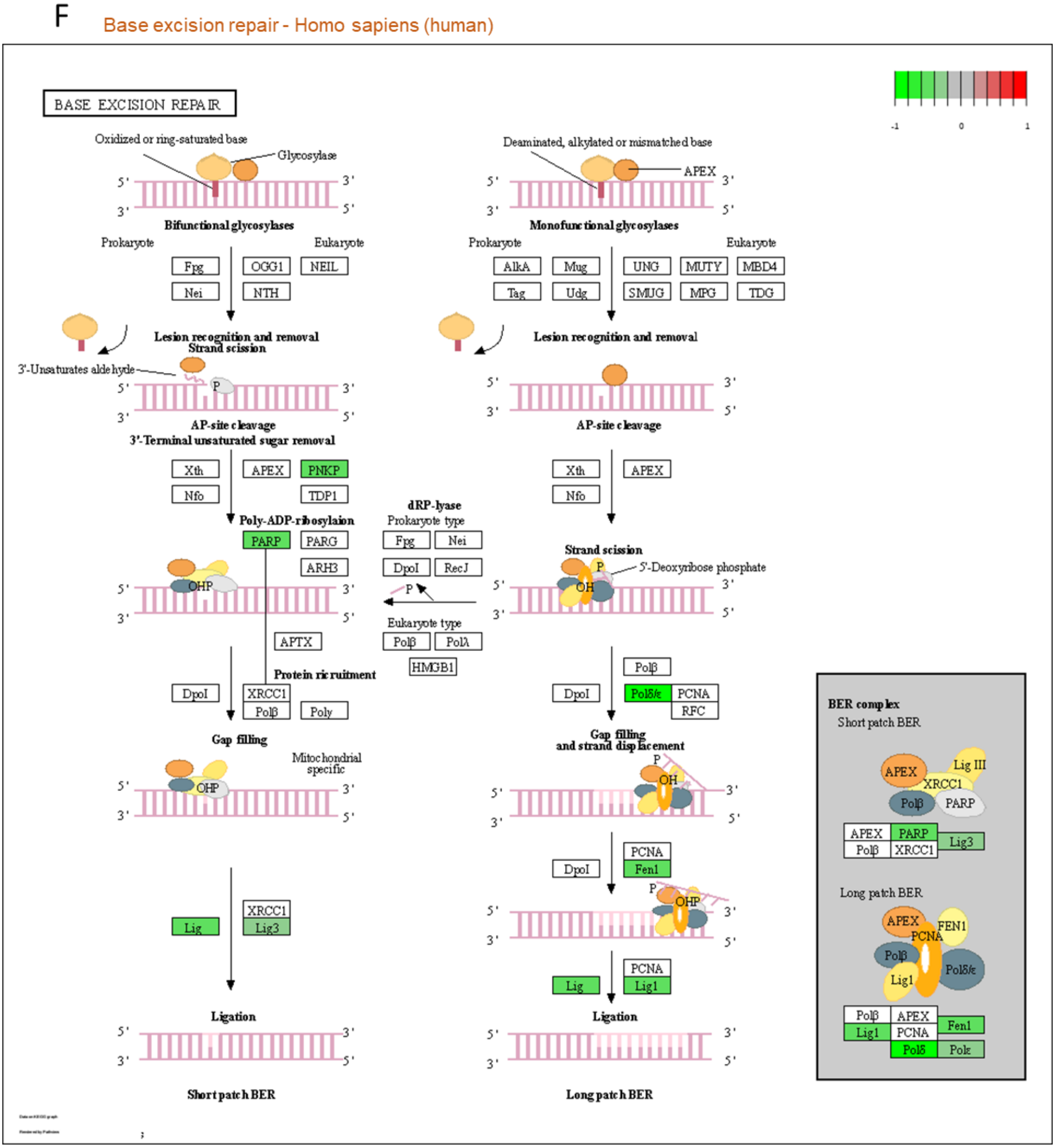
Base excision repair - Homo sapiens (human)
BASE EXCISION REPAIR
Oxidized or ring-saturated base
Glycosylase
Bifunctional glycosylases
Prokaryote
Eukaryote
Fpg
OGG1
NEIL
Nei
NTH
Lesion recognition and removal
Strand scission
3'-Unsaturates aldehyde
AP-site cleavage
3'-Terminal unsaturated sugar removal
Xth
APEX
PNKP
Nfo
TDP1
Poly-ADP-ribosylaion
PARP
PARG
ARH3
APTX
Protein ricruitment
DpoI
XRCC1
Polβ
Poly
Gap filling
Mitochondrial specific
Lig
Lig3
Ligation
Short patch BER
Deaminated, alkylated or mismatched base
APEX
Monofunctional glycosylases
AlkA
Mug
UNG
MUTY
MBD4
Tag
Udg
SMUG
MPG
TDG
dRP-lyase
Prokaryote type
RecJ
Eukaryote type
PolΛ
HMGB1
Strand scission
5'-Deoxyribose phosphate
Polδ/ε
PCNA
RFC
Gap filling and strand displacement
Fen1
Lig1
Long patch BER
BER complex
Short patch BER
Lig III
PARP
Long patch BER
FEN1
Polε
Polδ

Supplementary Figure.13. DNA repair related KEGG pathway plot for 278 DEG list. (A) Non-homologous end-joining KEGG pathway plot. (B) Fanconi anemia KEGG pathway plot. (C) Mismatch repair KEGG pathway plot. (D) Homologous recombination KEGG pathway plot. (E) Nucleotide excision repair KEGG pathway plot. (F) Base excision repair KEGG pathway plot.

Suppmentary figure 14

Rejuvenation related pathway (278 genes upregulated in 2XL compared to unload)

A mTOR signaling pathway

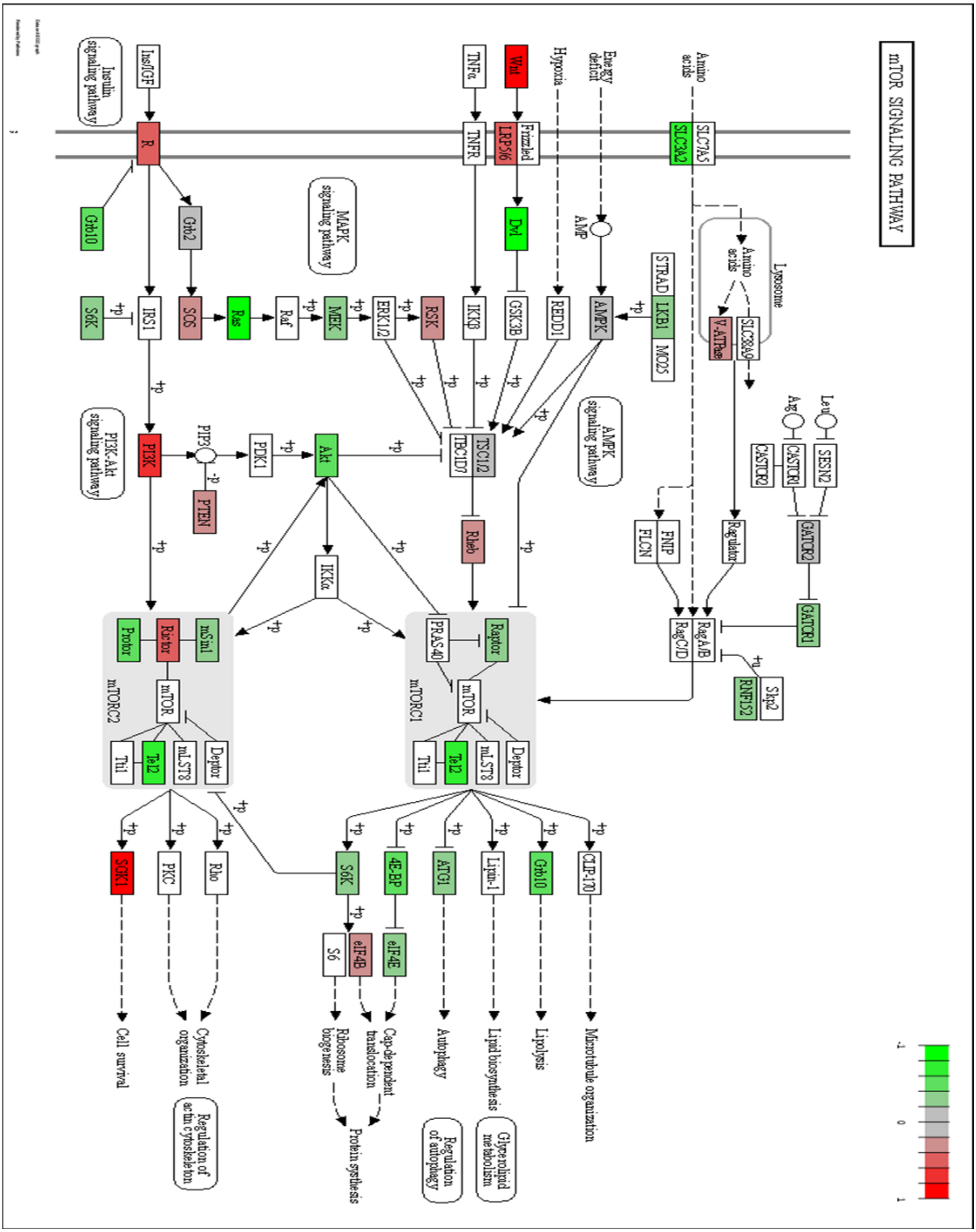

### B Longevity signaling pathway

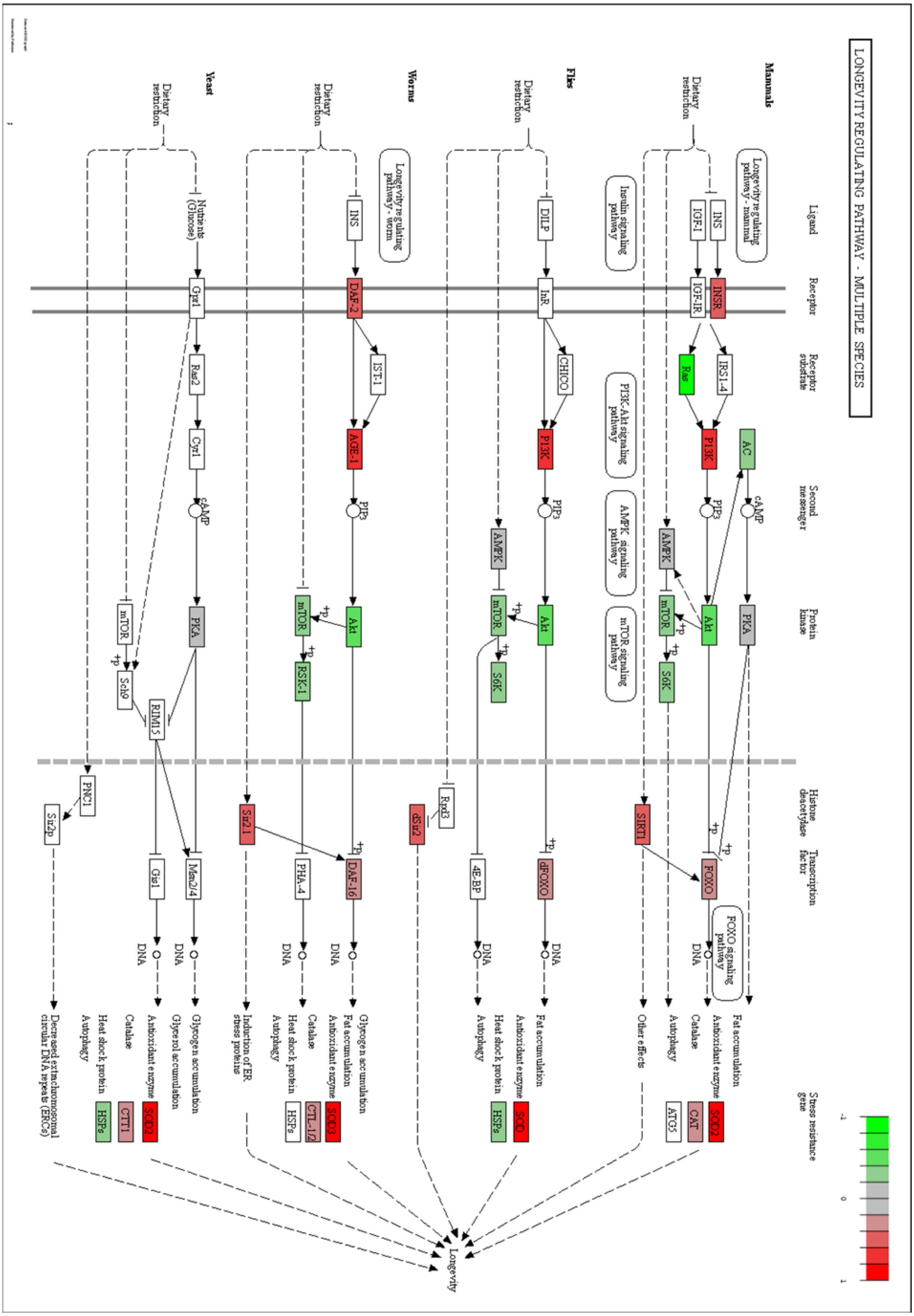

Suppmentary figure 14

C FOXO signaling pathway

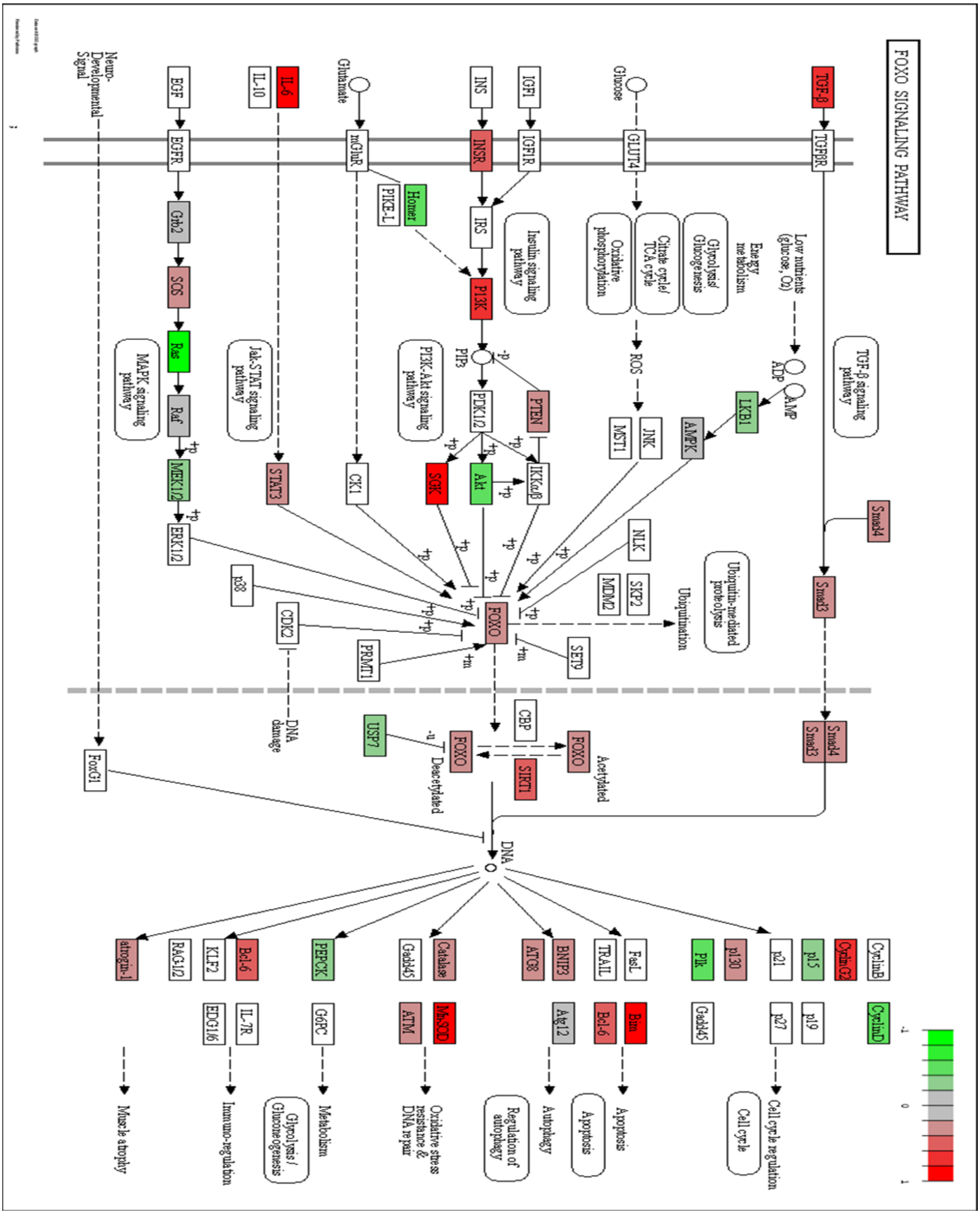

D AMPK signaling pathway

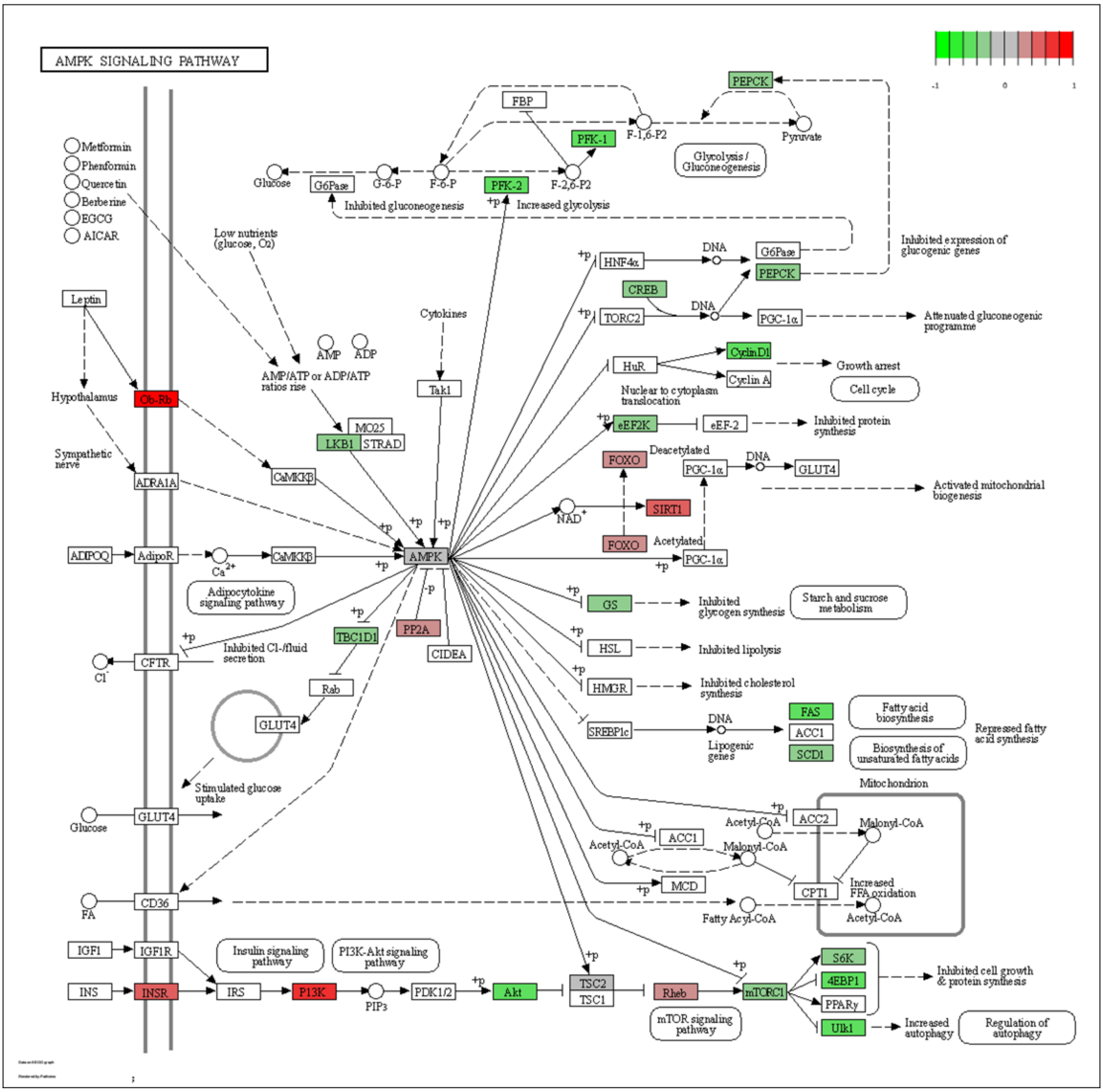

Supplementary Figure.S14. Rejuvenation related KEGG pathway plot for 278 DEG list. (A) mTOR KEGG pathway plot. (B) Longevity KEGG pathway plot. (C) FOXO KEGG pathway plot. (D) AMPK KEGG pathway plot.

Supplementary figure 15

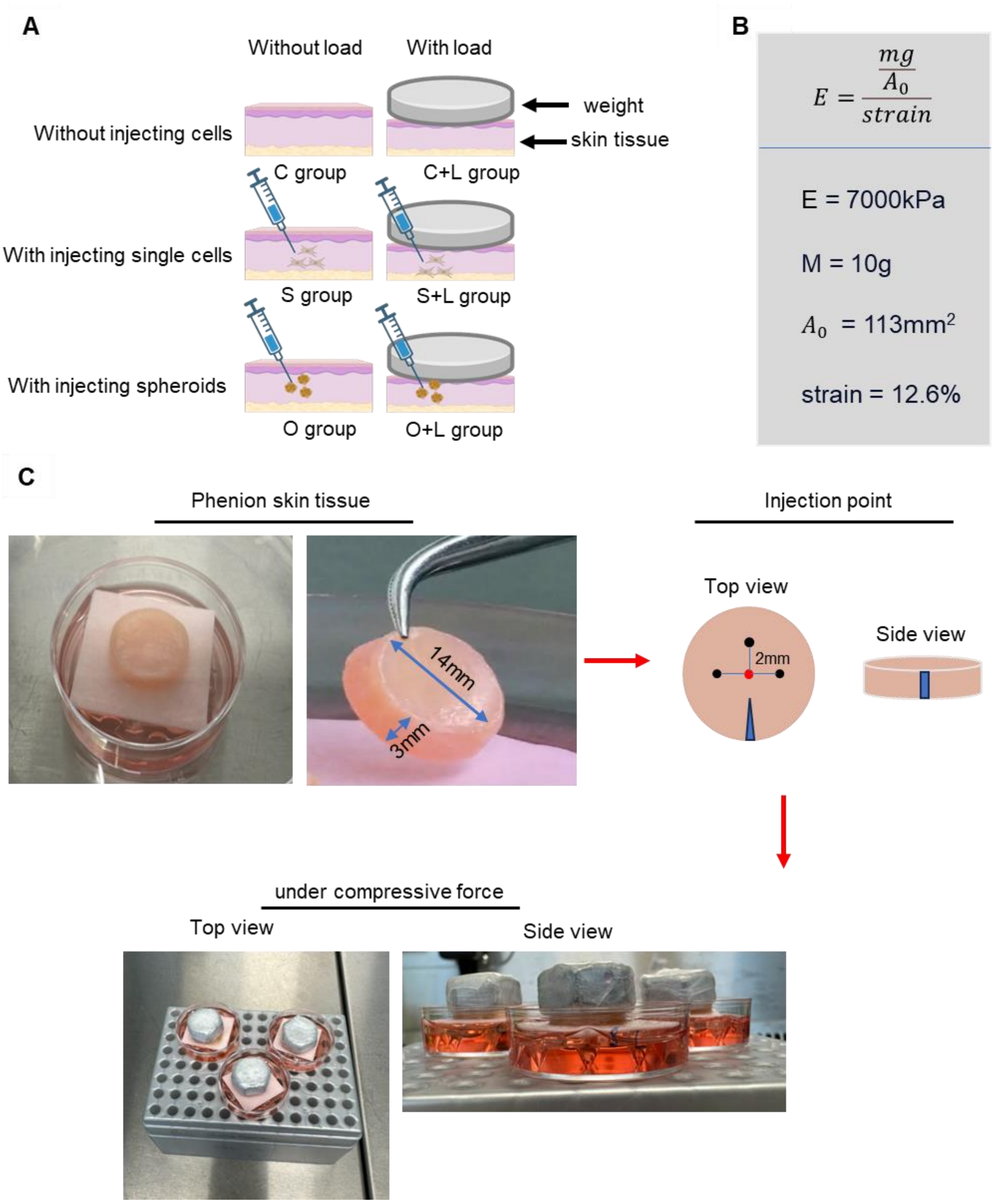

Supplementary Figure.S15. Illustration of experimental procedure. (A) Schematic representation of injecting single cell or spheroids with load or without load. (B) Calculation of deformation under compressive force. (C) Representative pictures of FT AGED skin tissue, injection point and under compressive force. Red dot means the center of skin tissue. Black dots mean injection point. Blue triangle means wounded region to identify the location for cryosection.

Suppmentary figure 16

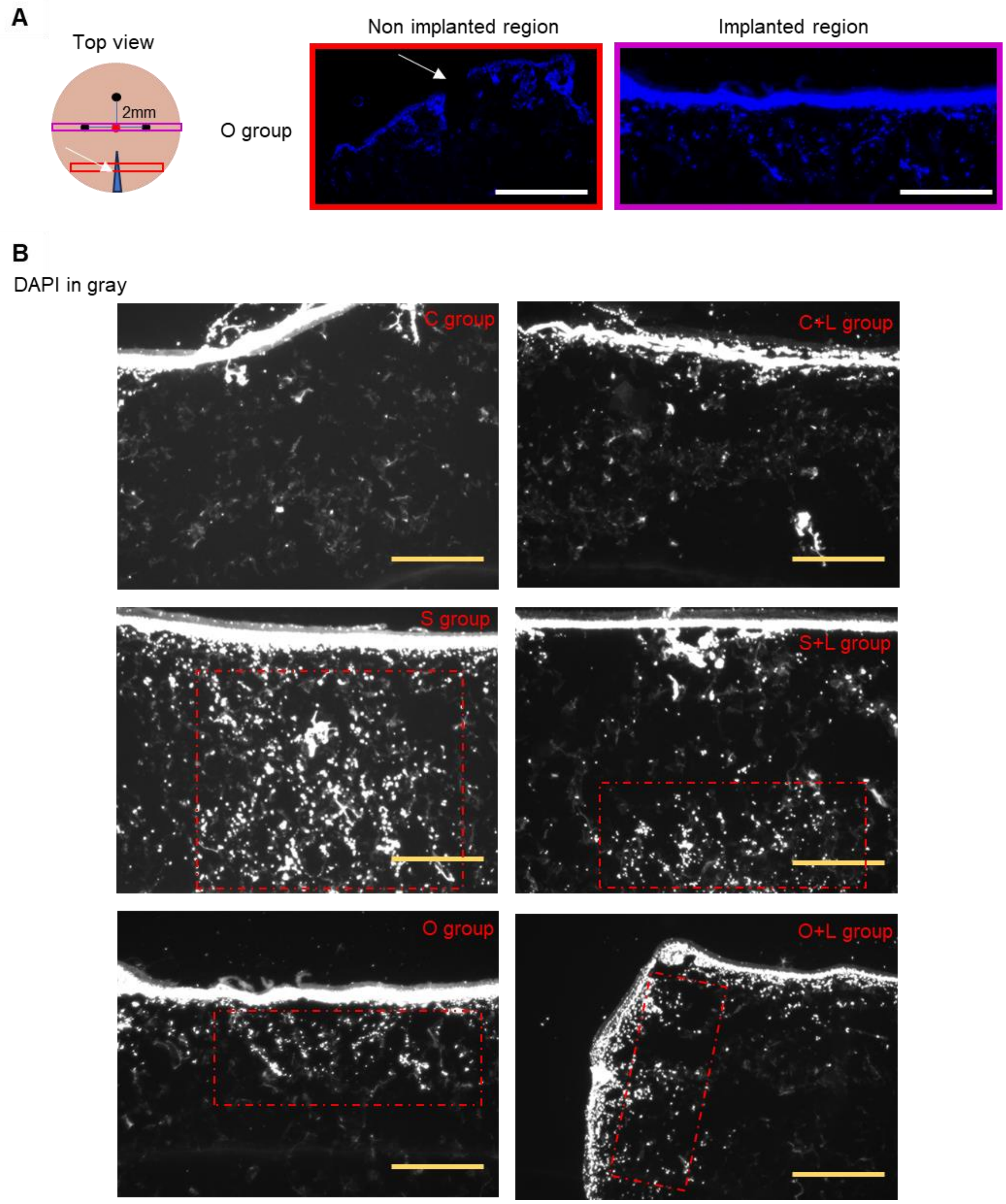


Supplementary Figure.S16. Check for injection point. (A) DAPI staining to check the injection point. Scale bar in red box is 1550 μ m. Scale bar in purple box is 750 μ m. (B) DAPI staining in gray color to show the injection point in every group. Scale bar: 750 μm. Red box means injection area.

Suppmentary figure 17

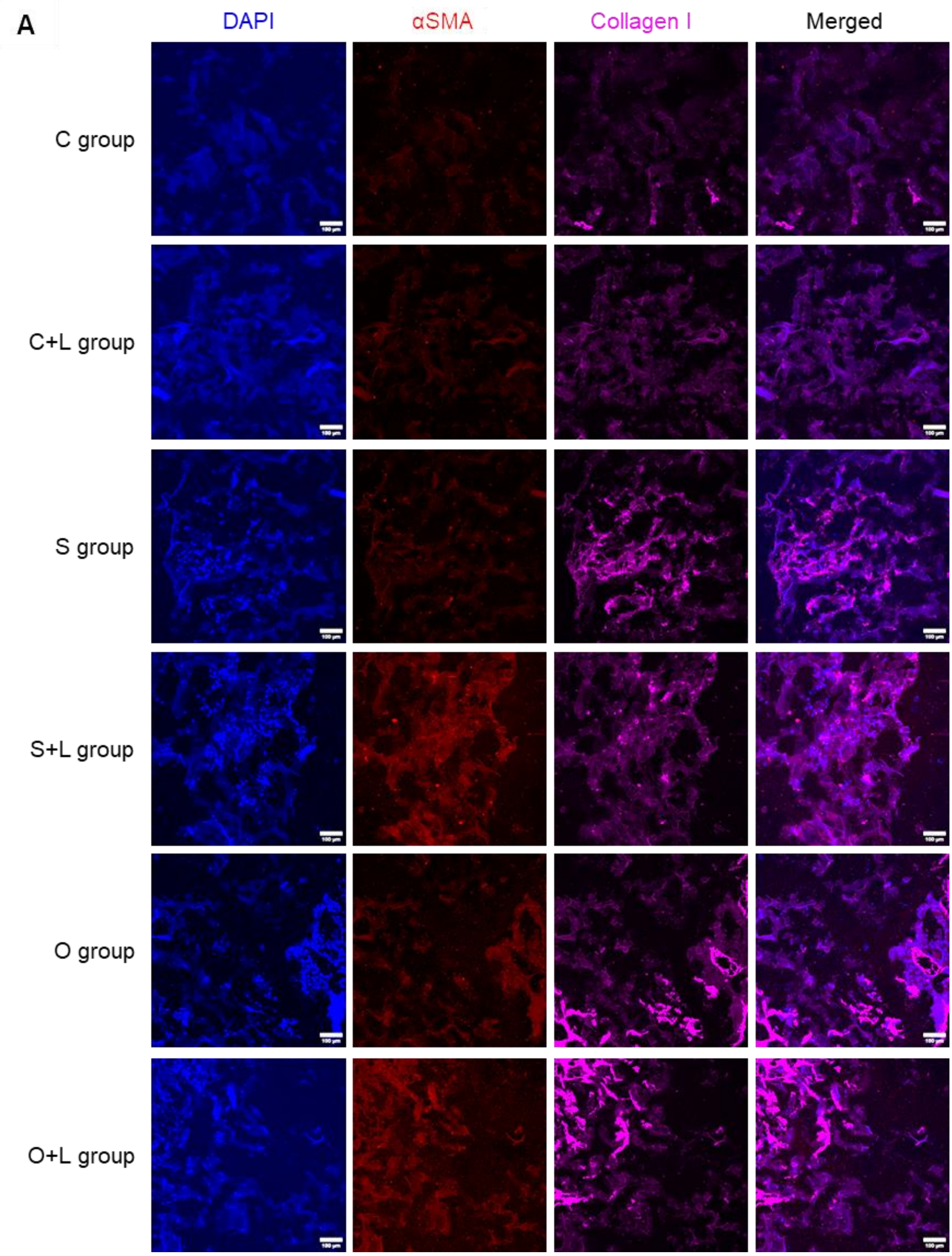

B

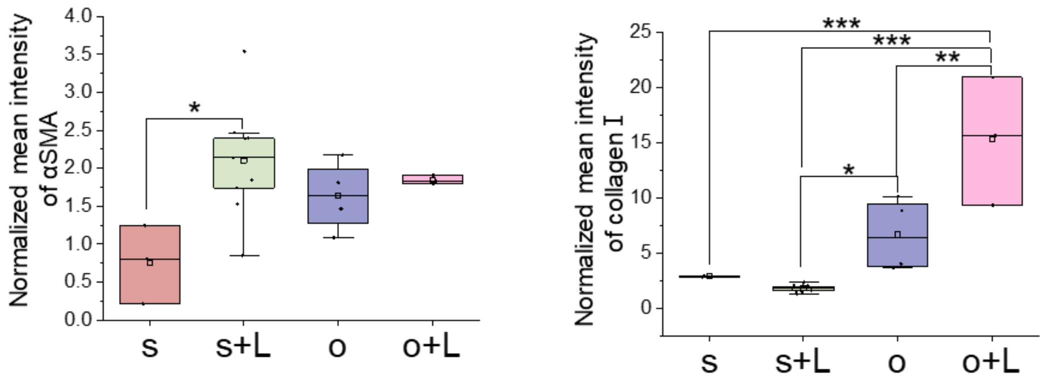


Supplementary Figure.S17. Individual channels for αSMA and collagen I. (A) Representative immunofluorescence confocal. (B) Quantification data of mean intensity per cell *from at least 3 replicates*. (Scale bar, 100μm). *P values were calculated by the one-way ANOVA method with Tukey's post hoc test. *P<0.05; **<0.01; ***P<0.001; No asterisks mean not significant.*

Suppmentary figure 18

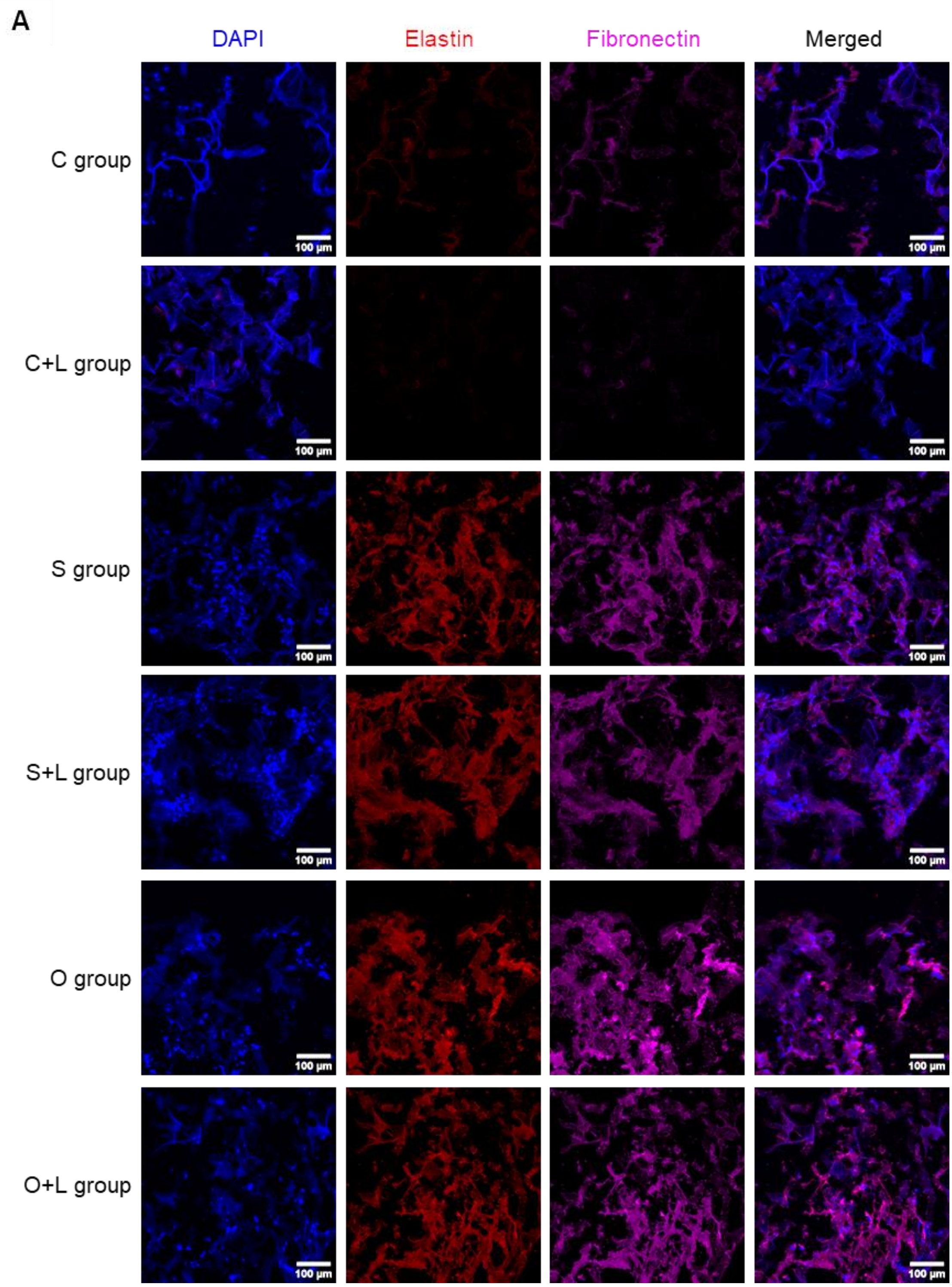

**B**

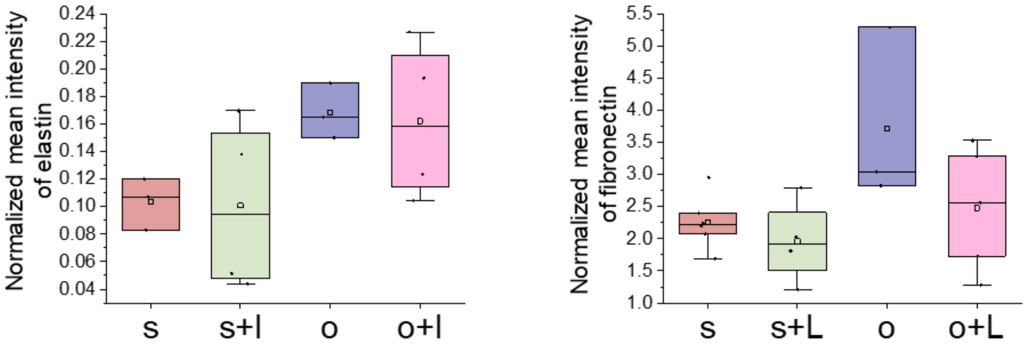


Supplementary Figure.S18. Individual channels for elastin and fibronectin. (A) Representative immunofluorescence confocal. (B) Quantification data of mean intensity per cell *from at least 3 replicates*. (Scale bar, 100μm). *P values were calculated by the one-way ANOVA method with Tukey's post hoc test. *P<0.05; **<0.01; ***P<0.001; No asterisks mean not significant.*

**Table S1.** List of primers (12).

| Primer (human) | Sequence (5'-3') |
|---|---|
| TRPV4-F | TCC ACC CTA TAT GAG TCC TCG |
| TRPV4-R | TAG GTG CCG TAG TCA AAC AGT |
| GDF15-F | TCA AGG TCG TGG GAC GTG ACA |
| GDF15-R | GCC GTG CGG ACG AAG ATT CT |
| RhoA-F | CGG GAG CTA GCC AAG ATG AAG |
| RhoA-R | CCT TGC AGA GCA GCT CTC GTA |
| RhoB-F | TGC TGA TCG TGT TCA GTA AG |
| RhoB-R | AGC ACA TGA GAA TGA CGT CG |
| RhoC-F | TCC TCA TCG TCT TCA GCA AG |
| RhoC-R | GAG GAT GAC ATC AGT GTC CG |
| Rac-1-F | AAC CAA TGC ATT TCC TGG AG |
| Rac-1-R | CAG ATT CAC CGG TTT TCC AT |
| ROCK1-F | AAG AGG GCA TTG TCA CAG CA |
| ROCK1-R | AGC ATC CAA TCC ATC CAG CA |

| | |
|---|---|
| Cdc42-F | GCC CGT GAC CTG AAG GCT GTC A |
| Cdc42-R | TGC TTT TAG TAT GAT GCC GAC ACC A |
| YAP-F | CTT CAA CGC CGT CAT GAA CC |
| YAP-R | GCA TCA GTA CTG GCC TGT CG |
| TAZ-F | CTT TCC TCA ATG GAG GGC CA |
| TAZ-R | GAA GTC CTC CGG AGT TGT GG |
| MMP1-F | AGA GCA GAT GTG GAC CAT GC |
| MMP1-R | TTG TCC CGA TGA TCT CCC CT |
| MMP2-F | TGA ACC AAC CAG CTG GCC TA |
| MMP2-R | AAG GTG TTC AGG TAT TGC ATG TG |
| PDGF-B-F | TGA GAA AGA TCG AGA TTG TGC G |
| PDGF-B-R | GGG CTT CGG GTC ACA GG |
| ARPC3-F | CCA AGA TGC CGG CTT ACC A |
| ARPC3-R | TCT GAT AGG CAA CAG TGC CA |
| Fibronectin-F | CCG CCG AAT GTA GGA CAA GA |
| Fibronectin-R | AGG GTT CTT CAT CAG TGC CA |
| COL1A1-F | GTC GAG GGC CAA GAC GAA |

| | |
|---|---|
| COL1A1-R | GTC TCG GTC ATG GTA CCT |
| follistatin-F | CCT GAG AAA GGC TAC CTG CC |
| follistatin-R | CAC AGG ACT TTG CTT TGA TAC AC |
| Vinculin -F | GGA TGA AGA GTT CCC TGA GCA |
| Vinculin -R | GAT GTC ATT GCC CTT GCT GG |
| PIEZO1-F | CAT AGG GGT CAC AAG GCT GG |
| PIEZO1-R | GAG GAG ACC ACC AAG ATG CC |
| IQGAP1-F | CGA AAT CTG GGC TCC ATT GC |
| IQGAP1-R | GCA GTT TGG AAA AAC CGT CTG A |
| LAMA1-F | AGT TTC GAA CCT CCT CGC AG |
| LAMA1-R | ATG GAA CAA GAC CTT GCC GT |
| GFP-KASH For | CG CTA GCG CTA CCG GTC GCC |
| GFP-KASH Rev | GATC GTCGAC CAG TTA TCT AGA TCC GGT G |
| GAPDH For | AGAAGGCTGGGGCTCATTTG |
| GAPDH Rev | AGGGGCCATCCACAGTCTTC |

**Table S2.** Antibody list.

| Antibody name | Brand & catalogue number | Final concentration |
|---|---|---|
| γH2AX | Rabbit, CST, 2577S | 1:250 |
| H3K9me3 | Rabbit, abcam, ab176916 | 1:2000 |
| H3K4mes | Rabbit, abcam, ab213224 | 1:500 |
| ɑSMA | Mouse, Merk, A2547 | 1:250 |
| ɑSMA | Mouse, ab7817 | 1:100 (skin tissue) |
| HP1a | Rabbit, Cell Signaling Technology, 2616s | 1:200 |
| *α*tubulin | Mouse,Merk , T5168 | 1 :2000 |
| pERK | Rabbit,Invitrogen, 36-8800 | 1:500 |
| LaminA/C | Mouse, Cell Signaling Technology, 4777S | 1:200 |
| LaminB | Rabbit, abcam, ab229025 | 1:500 |
| pMLC | Mouse, CTS, 3675S | 1:200 |
| Fibronectin | Rabbit, abcam, ab268020 | 1:100 (skin tissue) |

| | | |
|---|---|---|
| Collagen I | Rabbit, abcam, ab316222 | 1:200 (skin tissue) |
| Elastin | Mouse, abcam, ab9519 | 1:200 (skin tissue) |
| Goat anti-Rabbit IgG (H+L) Highly Cross-Adsorbed Secondary Antibody, Alexa Fluor™ Plus 647 | Invitrogen, A32733 | 1:300<br>or 1:500 in skin tissue sample |
| Goat anti-Mouse IgG (H+L) Highly Cross-Adsorbed Secondary Antibody, Alexa Fluor™ Plus 647 | Invitrogen, A32728 | 1:300 |
| Donkey anti-Rabbit IgG (H+L) Highly Cross-Adsorbed Secondary Antibody, Alexa Fluor™ Plus 555 | Invitrogen, A32794 | 1:300 |
| Donkey anti-Mouse IgG (H+L) Highly Cross-Adsorbed Secondary Antibody, Alexa Fluor™ Plus 555 | Invitrogen, A32773 | 1:300<br>or 1:500 in skin tissue sample |

| | | |
|---|---|---|
| Goat anti-Rabbit IgG (H+L) Highly Cross-Adsorbed Secondary Antibody, Alexa Fluor™ Plus 488 | Invitrogen, A32731 | 1:300 |
| Goat anti-Mouse IgG (H+L) Highly Cross-Adsorbed Secondary Antibody, Alexa Fluor™ Plus 488 | Invitrogen, A32723 | 1:300 |

**Table S3.** List of inhibitors.

| Drug name | Brand & catalogue number | Final concentration |
|---|---|---|
| Y27632 | Sigma, Y0503 | 20*μM* (Venkatachalapathy et al., 2022) |
| Blebbistatin | Sigma, B0560 | 25*μM* (Makhija et al., 2016) |
| Cytochalasin D | Sigma, C2618 | 5*μM* (Potolitsyna et al., 2024) |
| Latrunculin A | SEA, L5163 | 200nM (Venkatachalapathy et al., 2022) |
| Nocodazole | Sigma, M1404 | 10*μ*g/ml (Maharana et al., 2012; Makhija et al., 2016) |
| PD98059 | Merk, 513000 | 20*μM* (Kalli et al., 2019) |
| PF-573228 | Sigma, PZ0117 | 1*μM* (Mukherjee et al., |

| | | |
|---|---|---|
| | | 2022) |
| KU-55933 | Sigma, SML1109 | 20uM (Demelash et al., 2017) |
| HPI-4 | Selleckchem, S8249 | 5uM (Chang et al., 2019) |
| Withaferin A | Merk, W4394 | 5uM (Sliogeryte & Gavara, 2019) |
| Cisplatin | Sigma, PHR1624 | 100uM (Pekeč et al., 2023) |

**Table S4.** Description of heatmap names in Figure 2D.

| Short name for feature | Description |
|---|---|
| std_radius | Standard deviation of the radial distances |
| eccentricity | Eccentricity of an ellipsoid with an equal second-order moment |

| | |
|---|---|
| a_r | Ratio of the minor to the major axis length |
| entropy | Entropy of the intensity distribution |
| i80_i20 | Ratio of the 80%-to-20%-tile of the intensity distribution |
| contrast_1 | Contrast of the GLCM matrix for a lag of 1 pixel |
| weighted_s-0-0 | Weighted spatial moment for p=0 and q=0 |
| weighted_n-3-3 | Weighted normalized spatial moment for p=3 and q=3 |
| weighted_c-0-0 | Weighted central spatial moment for p=0 and q=0 |
| weighted_ c-0-3 | Weighted central spatial moment for p=0 and q=3 |
| moments_s_1-0 | Spatial moment for p=1 and q=0 |
| moments_s_2-0 | Spatial moment for p=2 and q=0 |
| moments_s_3-0 | Spatial moment for p=3 and q=0 |
| moments_c_2-0 | Central spatial moments for p=2 and q=0 |

| | |
|---|---|
| moments_ c_2-1 | Central spatial moments for p=2 and q=1 |
| moments_hu-1 | Hu spatial moments of order 1 |
| width_pos_curv | Average width of the prominent peaks of the positive curvature distribution of the boundary |
| min_calliper | Minimal caliper distance |
| d25_radius | 25%-tile of the radial distances |
| minor_length | Length of the minor axis of an ellipsoid of equal area |

**Movies S1 to S3**

Video 1 displays the z-stack images presented in Figure S1C Spheroid #1 (nucleus number is 62).

Video 2 displays the z-stack images presented in Figure S1C Spheroid #2 (nucleus number is 55).

Video 3 displays the z-stack images presented in Figure S1C Spheroid #3 (nucleus number is 36).

**SI Reference**


Chang, W., Wang, Y., Luxton, G. W. G., Östlund, C., Worman, H. J., & Gundersen, G. G. (2019). Imbalanced nucleocytoskeletal connections create common polarity defects in progeria and physiological aging. *Proceedings of the National Academy of Sciences*, *116*(9), 3578–3583. https://doi.org/10.1073/pnas.1809683116

Demelash, A., Pfannenstiel, L. W., Liu, L., & Gastman, B. R. (2017). Mcl-1 regulates reactive oxygen species via NOX4 during chemotherapy-induced senescence. *Oncotarget*, *8*(17), 28154–28168. https://doi.org/10.18632/oncotarget.15962

Kalli, M., Voutouri, C., Minia, A., Pliaka, V., Fotis, C., Alexopoulos, L. G., & Stylianopoulos, T. (2019). Mechanical Compression Regulates Brain Cancer Cell Migration Through MEK1/Erk1 Pathway Activation and GDF15 Expression. *Frontiers in Oncology*, *9*, 992. https://doi.org/10.3389/fonc.2019.00992

Maharana, S., Sharma, D., Shi, X., & Shivashankar, G. V. (2012). Dynamic Organization of Transcription Compartments Is Dependent on Functional Nuclear Architecture. *Biophysical Journal*, *103*(5), 851–859. https://doi.org/10.1016/j.bpj.2012.06.036

Makhija, E., Jokhun, D. S., & Shivashankar, G. V. (2016). Nuclear deformability and telomere dynamics are regulated by cell geometric constraints. *Proceedings of the National Academy of Sciences*, *113*(1). https://doi.org/10.1073/pnas.1513189113

Mukherjee, D., Hao, J., Lu, H., Lahiri, S. K., Yu, L., & Zhao, J. (2022). KLF8 promotes invasive outgrowth of breast cancer by inducing filopodium-like protrusions via CXCR4. *American Journal of Translational Research*, *14*(2), 1220–1233.

Pekeč, T., Venkatachalapathy, S., Shim, A. R., Paysan, D., Grzmil, M., Schibli, R., Béhé, M., & Shivashankar, G. V. (2023). Detecting radio- and chemoresistant cells in 3D cancer co-cultures using chromatin biomarkers. *Scientific Reports*, *13*(1), 20662. https://doi.org/10.1038/s41598-023-47287-2

Potolitsyna, E., Pickering, S. H., Bellanger, A., Germier, T., Collas, P., & Briand, N. (2024). Cytoskeletal rearrangement precedes nucleolar remodeling during adipogenesis. *Communications Biology*, *7*(1), 458. https://doi.org/10.1038/s42003-024-06153-1

Sliogeryte, K., & Gavara, N. (2019). Vimentin Plays a Crucial Role in Fibroblast Ageing by Regulating Biophysical Properties and Cell Migration. *Cells*, *8*(10), 1164. https://doi.org/10.3390/cells8101164

Venkatachalapathy, S., Sreekumar, D., Ratna, P., & Shivashankar, G. V. (2022). Actomyosin contractility as a mechanical checkpoint for cell state transitions. *Scientific Reports*, *12*(1), 16063. https://doi.org/10.1038/s41598-022-20089-8

Xie, Y., Mansouri, M., Rizk, A., & Berger, P. (2019). Regulation of VEGFR2 trafficking and signaling by Rab GTPase-activating proteins. *Scientific Reports*, *9*(1), 13342. https://doi.org/10.1038/s41598-019-49646-4
